# Evolutionary and ontogenetic changes of the anatomical organization and modularity in the skull of archosaurs

**DOI:** 10.1101/2020.02.21.960435

**Authors:** Hiu Wai Lee, Borja Esteve-Altava, Arkhat Abzhanov

## Abstract

Comparative anatomy studies of the skull of archosaurs provide insights on the mechanisms of evolution for the morphologically and functionally diverse species of crocodiles and birds. One of the key attributes of skull evolution is the anatomical changes associated with the physical arrangement of cranial bones. Here, we compare the changes in anatomical organization and modularity of the skull of extinct and extant archosaurs using an Anatomical Network Analysis approach. We show that the number of bones, their topological arrangement, and modular organization can discriminate birds from non-avian dinosaurs, and crurotarsans. We could also discriminate extant taxa from extinct species when adult birds were included. By comparing within the same framework, juveniles and adults for crown birds and alligator *(Alligator mississippiensis),* we find that adult and juvenile alligator skulls are topologically similar, whereas juvenile bird skulls have a morphological complexity and anisomerism more similar to those of non-avian dinosaurs and crurotarsans than of their own adult forms. Clade-specific ontogenetic differences in skull organization, such as extensive postnatal fusion of cranial bones in crown birds, can explain this pattern. The fact that juvenile and adult skulls in birds do share a similar anatomical integration suggests the presence of a specific constraint to their ontogenetic growth.

## INTRODUCTION

The skulls of archosaurs are morphologically and functionally diverse, with clade-specific specialized features that set apart crurotarsans (extant crocodilians and their stem lineage) from avemetatarsalians (birds and non-avian dinosaurs)^1–7^, as reviewed by Brusatte and colleagues^8^. The evolution and diversification of the skull of archosaurs have been associated with changes in the patterns of phenotypic integration and modularity^9–13^. For more information on integration and modularity in shape, see the review by Klingenberg^14^. Different regions of the skull may act as anatomical modules that can evolve, function, and develop semi-independently from one another. Bones within a same module tend to co-vary in shape and size more with each other than with bones from other such variational modules^15–18^. In addition, the bones of the skull can also modify their physical articulations so that some groups of bones are more structurally integrated than others, and, hence, we can recognize them as distinct anatomical-network modules, which had been defined by Eble as a type of organizational modules^15,19,20^. The relationship between anatomical-network modules and variational modules is not yet fully understood, but it is thought that network anatomy constrain growth patterns and shape variation^21–23^.

Changes in the anatomical organization of the skull in archosaurs have been concomitant with a broader evolutionary trend in tetrapods toward a reduction in the number of skull bones due to loses and fusions, a phenomenon known as the Williston’s law^24–26^. Understanding how the bones are globally arranged to each other allows us to measure the anatomical complexity and organization of body parts, and explain how structural constraints might have influenced the direction of evolution^25–28^. Werneburg and colleagues compared the skull network-anatomy of a highly derived *Tyrannosaurus rex, Alligator mississippiensis* and *Gallus gallus* with that of an opossum, a tuatara, and a turtle^29^. They found that the tyrannosaur has the most modular skull organization among these amniotes, with a modular separation of the snout in upper and lower sub-modules and the presence of a lower adductor chamber module. However, the specific anatomical changes in the organization of the archosaur skull during their evolutionary transitions more generally have never been characterized. More recently, Plateau and Foth used anatomical network analysis to study postnatal ontogenetic changes in the skulls of crown bird and non-avian theropods^30^. They found that early juvenile crown birds have skulls that are less integrated and more modular than those of more derived birds, resembling their non-avian theropod ancestors.

Here, we compared the anatomical organization and modularity of the skull of archosaurs using Anatomical Network Analysis (AnNA)^31^ to highlight how skull topology has changed in evolutionary and developmental scales. We chose AnNA over more conventional methods, such as geometric morphometrics, to understand how major re-organizations of the skull (i.e., loss and fusion of bones) affect the overall anatomy regardless of shape. We created network models of the skull for 21 species of archosaurs, including taxa representing key evolutionary transitions from early pseudosuchians to crocodiles, from non-avian theropods to modern birds, and from paleognath birds to neognaths (Fig. 2). Our dataset also includes a representative ornithischian, a sauropodomorph, and a basal saurischian (Supplementary Information 1) for comparison. To understand the significance of the ontogenetic transitions in archosaur skulls, we provided our dataset with juvenile skulls for extant birds and alligator. Network models of the skull were built by coding individual cranial bones and their articulations with other bones as the nodes and links of a network, respectively (Fig. 1). Network modules, defined as a group of bones with more articulations among them than to other bones outside the module, were identified heuristically using a community detection algorithm. We compared skull architectures using topological variables (i.e. network parameters) that capture whole-skull anatomical feature (modelling and analysis of anatomical networks were detailed previously^20,25,31^).

**Figure 1.**
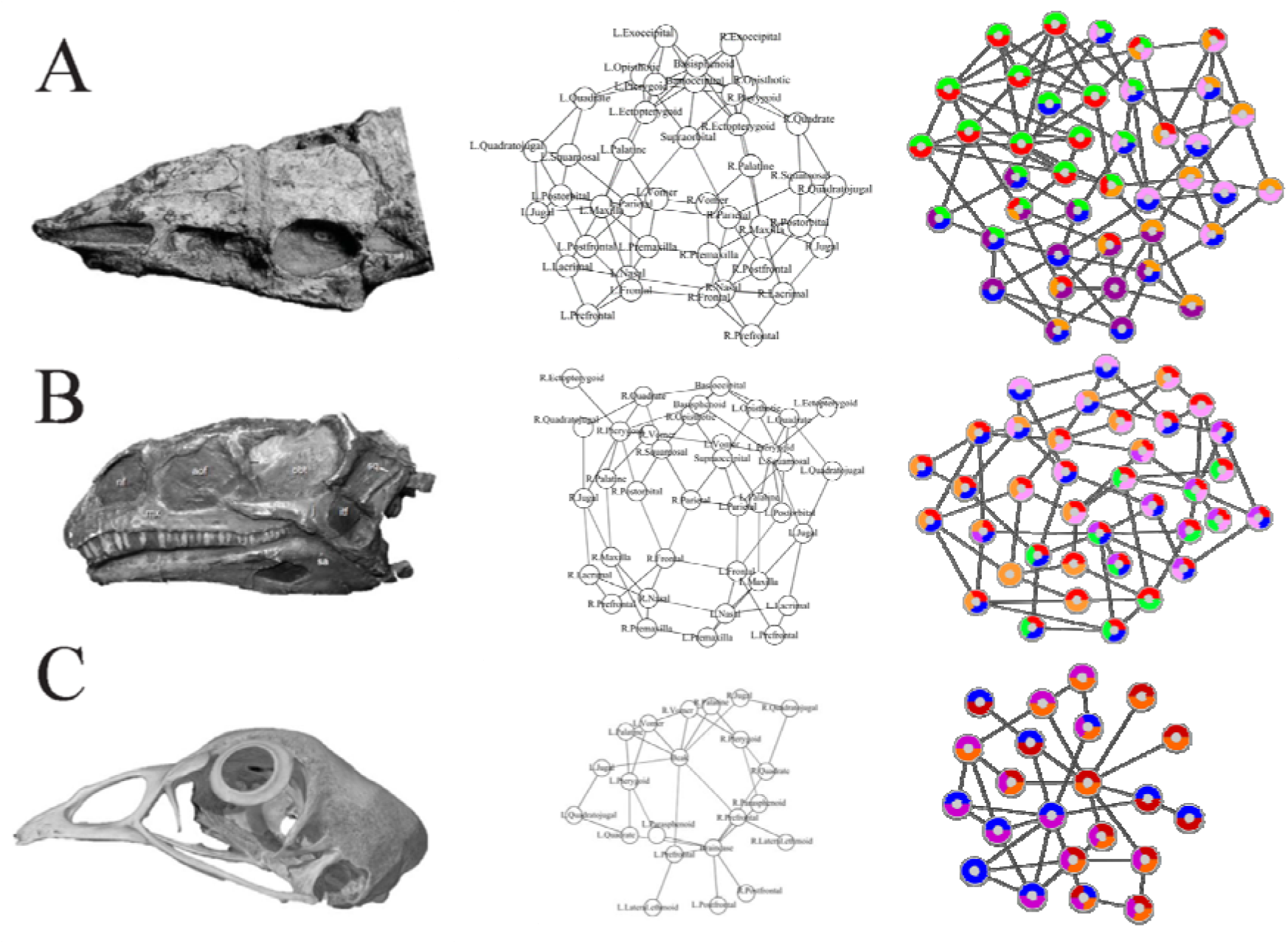
Anatomical network models. Example of the network models for three archosaurian skulls: (A) *Aetosaurus* from Schoch (2007)^63^; (B) *Plateosaurus* from Prieto-Marquez & Norell (2011)^107^; (C) *Gallus* from Digimorph. The pair-wise articulations among the bones of skulls (left) are formalized as network models (middle) and later analyzed, for example, to identify the skull anatomical node-based modules (right). See methods for details.

Networks and network modules and their respective complexity, integration, modularity, and anisomerism could be quantified by these network parameters: density of connections, clustering coefficient, path length, heterogeneity of connections, and parcellation^20, 23, 31, 32^. Here, complexity is defined as the relationship of bones in a skull and is associated with how abundant are the interactions that bones have with each other (i.e. density of connections), how interdependent or integrated the bones are (i.e. clustering coefficient), and proximity between nodes (i.e. path length). A more complex network would have higher density, higher clustering coefficient, and shorter path length. Anisomerism is defined as a deviation among anatomical parts^33^ and could be observed by the specialization of bones and measured by heterogeneity of connections, i.e. how each bone has a different number of connection^25^. Modularity is measured by parcellation, which is the number of modules and the consistency in the number of bones per module.

## MATERIAL AND METHODS

### Sampling

We sampled extinct and extant species, and for some forms included both adults and juveniles to account for ontogenetic trends within archosaurs. Namely, adults *Aetosaurus ferratus, Archaeopteryx lithographica, Citipati osmolskae, Coelophysis bauri, Compsognathus longipes, Dakosaurus andiniensis, Desmatosuchus haplocerus, Dibothrosuchus elaphros, Dilophosaurus wetherilli, Eoraptor lunensis, Ichthyornis dispar, Plateosaurus engelhardti, Psittacosaurus lujiatunensis*, *Riojasuchus tenuisceps, Sphenosuchus acutus, Velociraptor mongoliensis, Gallus gallus, Geospiza fortis* and *Nothura maculosa;* and juveniles *Gallus gallus, Geospiza fortis, Nothura maculosa* and *Alligator mississippiensis.* Within our sample set, eight species represent the transition from crurotarsan archosaur ancestor to modern crocodilians and 13 species represent the transition from non-avian theropods to modern birds as described previously^34–43^. Due to the sample size limitation for extinct taxa, reconstructed and type forms were used to represent each taxon and intraspecific variation could not be accounted for.

### Phylogenetic Context

We created a phylogenetic tree (Figure 2) based on the previous studies^34–37,39–44^. The tree was calibrated using the R package paleotree^45^ by the conservative “equal” method^46,47^; branching events were constrained using the minimum dates for known internal nodes based on fossil data from Benton and Donoghue^48^ (listed in Table S3) and the first and last occurrences of all 21 species from the Paleobiology Database using the paleobioDB package^49^ in R. Because there were two extinct *Nothura* species in the Paleobiology Database, the last occurrence for extant *Nothura* species was adjusted to 0 (Table S2).

**Figure 2.**
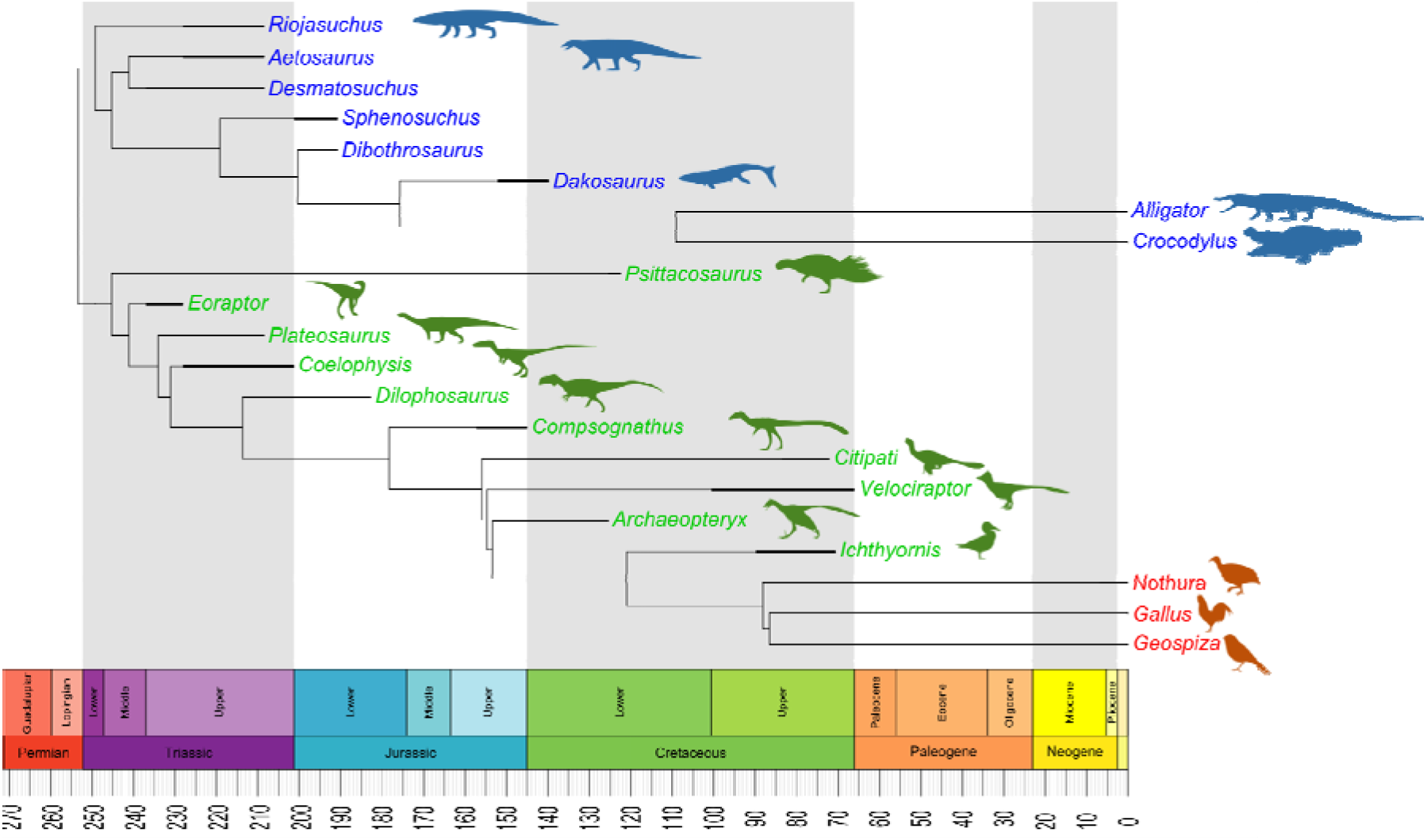
Phylogenetic framework. A phylogenetic tree was created based on the evolutionary relations among taxa as detailed in previous work^34–43^. Bifurcation times were calibrated based on fossil dates from Benton and Donoghue^48^ using the equal method in the paleotree package^45–47^. First and last occurrences were from Paleobiology Database (details listed in Table S2). Silhouettes were from Phylopic.org. See methods for details.

### Network Modelling

We built anatomical network models for each archosaur skull in our sample set based on detailed literature descriptions and CT scans of complete skulls (see Supplementary Information 1). Skull bones were represented as the nodes of the network model and their pair-wise articulations (e.g. sutures and synchondroses) were represented as links between pairs of nodes (Figure 1). Skull network models were formalized as binary adjacency matrices, in which a 1 codes for two bones articulating and a 0 codes for absence of articulation. Bones that were fused together without trace of a suture in the specimens examined were formalized as a single individual bone.

### Network Analysis

Following Esteve-Altava et al^28^, we quantified the following topological variables for each network model: the number of nodes (N), the number of links (K), the density of connections (D), the mean clustering coefficient (C), the mean path length (L), the heterogeneity of connections (H), the assortativity of connections (A), and the parcellation (P). The morphological interpretation of these topological variables has been detailed elsewhere^28^. A summary is provided here. N and K represent the direct count of the number of individual bones and articulations observed in the skull. D is the number of connections divided by the maximum number of possible connections (it ranges from 0 to 1); D is a proxy measure for morphological complexity. C is the average number of neighboring bones that connect to one another in a network (i.e., actual triangles of nodes compared to the maximum possible): a value close to 1 shows all neighboring bones connect to each other while a value close to 0 shows neighboring bones do not connect to each other; C is a proxy measure for anatomical integration derived from co-dependency between bones. L measures average number of links separating two nodes (it ranges from 1 to N-1); L is a proxy measure of anatomical integration derived from the effective proximity between bones. H measures how heterogeneous connections are in a network: skulls composed of bones with a different number of articulations have higher H values. If all bones had the same number of connections (i.e., H = 0), it means that all bones were connected in the same way and the skull had a regular shape. A measures whether nodes with the same number of connections connect to each other (it ranges from −1 to 1); H and A are a proxy measure for anisomerism or diversification of bones. P measures the number of modules and the uniformity in the number of bones they group (it ranges from 0 to 1); P is a proxy for the degree of modularity in the skull. Calculating P requires a given partition of the network into modules (see next below).

Network parameters were quantified in R^50^ using the igraph package^51^. Networks visualization was made using the visNetwork package^52^ and Cytoscape^53^.

### Principal Component Analysis

We performed a Principal Component Analysis (PCA) of the eight topological variables with a singular value decomposition of the centered and scaled measures. On the resulting PCs, we used a PERMANOVA (10,000 iterations) to test whether topological variables discriminate between: (1) Avialae and non-Avialae; (2) adults and juveniles; (3) extinct and extant; (4) Crurotarsi and Avemetatarsalia; (5) Neornithes and non-Neornithes; (6) early flight, can do soaring flight, can do flapping flight, gliding, and flightless (details in Table S5); (7) Crurotarsi, non-avian Dinosauria, and Aves; and (8) carnivorous, omnivorous, and herbivorous (dietary information in Supplementary Information 4). First, we performed the tests listed above for all archosaurs. Then, we repeated these tests for a sub-sample that included all archosaurs, except for all modern birds. Next, we repeated these tests for a sub-sample that included all archosaurs, except for adult birds.

### Modularity Analysis

To find the optimal partition into network modules we used a node-based informed modularity strategy^54^. This method starting with the local modularity around every individual node, using cluster_spinglass function in igraph^51^, then it returns the modular organization of the network by merging non-redundant modules and assessing their intersection statistically using combinatorial theory^55^.

## RESULTS

### Topological discrimination of skull bones

A Principal Component Analysis (PCA) of the eight topological variables measured in skull network models discriminates skulls with different anatomical organizations (Figs. S1-S3). When all sampled skulls are compared together, the first three principal components (PCs) explain 89.4% of the total variation of the sample. PC1 (57.5%) discriminates skulls by number of their bones (N), density of connections (D), and degree of modularity (P). PC2 (21.3%) discriminates skulls by their degree of integration (C) and anisomerism (H). Finally, PC3 (10.6%) discriminates skulls by whether bones with similar number of articulations connect with each other (A).

PERMANOVA tests confirm that different skull anatomies map onto different regions of the morphospace. Thus, we can discriminate: Avialae (Aves plus *Ichthyornis* and *Archaeopteryx)* versus non-Avialae (F_1,23_ = 4.124, *p* = 0.006699; Fig. 3B); Neornithes plus toothless archosaurs versus archosaurs with teeth (F_1,23_ = 6.99, *p* = 0.0005999; Fig. 3C); Aves (include all modern birds) versus Crurotarsi versus non-avian Dinosauria (F_2,22_ = 3.837, *p* = 0.000699; Fig. 3D); and extant and extinct species (F_1,23_ = 4.304, *p* = 0.0017; Fig. S1C). However, we find no statistically significant difference in morphospace occupation between crurotarsans and avemetatarsalians (F_1,23_ = 1.46, *p* = 0.2002, Fig. S1D).

**Figure 3.**
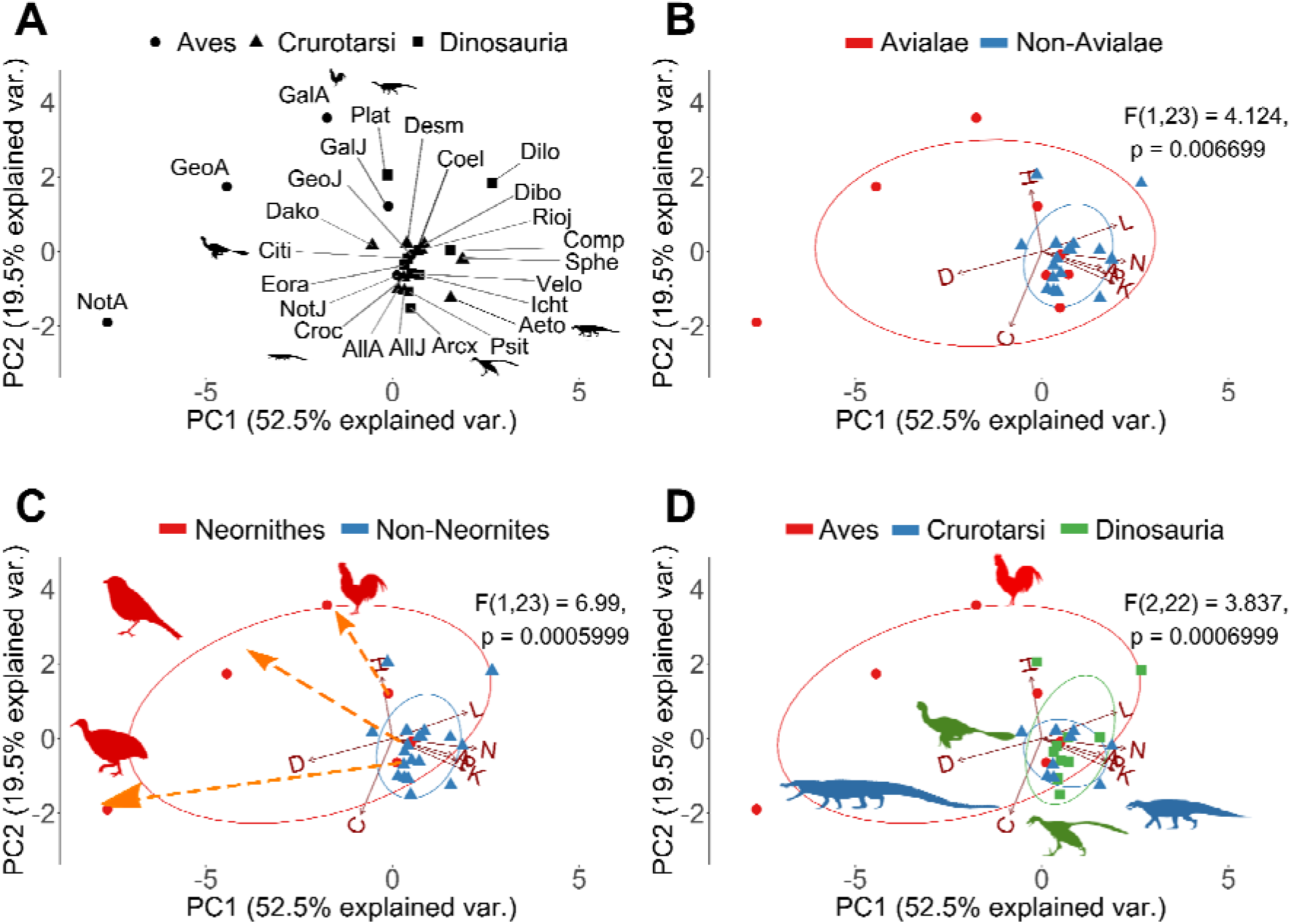
Principal components decomposition of topological variables. (A) Skull distribution for each taxon (see labels below). (B) Comparison of Avialae versus non-Avialae shows that non-Avialae occupy part of the Avialae morphospace. (C) Comparison of Neornithes versus non-Neornithes shows that non-Neornithes overlap with part of the Neornithes morphospace. Orange dotted arrows show the ontogenetic change in modern birds from juvenile stage to adult stage. (D) Comparison of Aves, Crurotarsi, and Dinosauria shows that they occupied different morphospace. Ellipses show a normal distribution confidence interval around groups for comparison. Labels: N, Number of nodes; K, Number of links; D, Density of Connection; C, Mean clustering coefficient; H, Heterogeneity of connection; L, Mean path length; A, Assortativity of connection; P, Parcellation. Aeto, *Aetosaurus;* AllA, adult *Alligator;* AllJ, juvenile *Alligator;* Arcx, *Archaeopteryx;* Citi, *Citipati;* Coel, *Coelophysis;* Comp, *Compsognathus;* Croc, *Crocodylus;* Dako, *Dakosaurus;* Desm, *Desmatosuchus;* Dibo, *Dibothrosuchus;* Dilo, *Dilophosaurus;* Eora, *Eoraptor;* GalA, adult *Gallus;* GalJ, juvenile *Gallus;* GeoA, adult *Geospiza;* GeoJ, juvenile *Geospiza;* Icht, *Ichthyornis;* NotA, adult *Nothura;* NotJ, juvenile *Nothura;* Plat, *Plateosaurus;* Psit, *Psittacosaurus;* Rioj, *Riojasuchus;* Sphe, *Sphenosuchus;* Velo, *Velociraptor.* Silhouettes were from Phylopic.org.

When all avians are excluded from the comparison, the first three PCs now explain 80.6% of the total variation (Figs. S4-6). PC1 (38.6%) discriminates skulls by the density of their inter-bone connections (D) and effective proximity (L). PC2 (22.6%) discriminates skulls by the number of bones and their articulations (N and K). Finally, PC3 (19.5%) now discriminates skulls by their anisomerism (H) and whether bones with the same number of connections connect to each (A). PERMANOVA tests could not discriminate between Crurotarsi and non-avian Dinosauria (F_1,17_ = 1.235, *p* = 0.3022; Fig. S4D), and between extant and extinct species (F_1,17_ = 2.274, *p* = 0.06399; Fig. S4C).

When only adult birds are excluded, the first three PCs explain 79.7% of the topological variation (Figs. S7-9). PC1 (35.8%), PC2 (24.5%), and PC3 (19.5%) discriminate skull similarly as when all birds are excluded (see above). PERMANOVA tests also could not discriminate between juvenile birds, crurotarsans, and non-avian dinosaurs (F_2,19_ = 1.682, *p* = 0.09649; Fig. S7D), and between extant and extinct species (F_1,20_ = 2.119, *p* = 0.06169; Fig. S7C).

Regardless of the sub-sample compared, we found no statistically significant difference in morphospace occupation between taxa stratified by flying ability and diet (Fig. S1E, see Supplementary Information 4 for details). This suggests that at least for the given sample set changes in cranial network-anatomy (i.e. how bones connect to each other) are independent of both dietary adaptations and the ability to fly.

### Number of network modules

The number of network modules identified in archosaur skulls ranged from one (i.e. fully integrated skull) in adult birds *Nothura maculosa* (the spotted tinamou) and *Geospiza fortis* (medium ground finch) to eight in the non-avian dinosaur *Citipati* (Table S10). The number of network modules within the studied taxa decreases during evolution of both major archosaurian clades: from 6 *(Riojasuchus)* to 4 *(Desmatosuchus,)* and from 6 *(Dibothrosuchus)* to 4 *(Dakosaurus* and all adult crocodilians) modules in Crurotarsi; from 6 *(Coelophysis)* to 4 *(Dilophosaurus* and *Compsognathus),* and from 8 *(Citipati)* to 4 *(Velociraptor, Archaeopteryx, Ichthyornis,* and juvenile modern birds) modules in theropod-juvenile bird transition (Fig. 4A and 4B, Table S10). We found no modular division of the skull in adult *Nothura* and *Geospiza.* This is most likely because these skulls are highly integrated due to the extensive cranial bone fusion in adults, which, in turn, results in a network with very few nodes. In general, skull networks are partitioned into overlapping modules.

**Figure 4.**
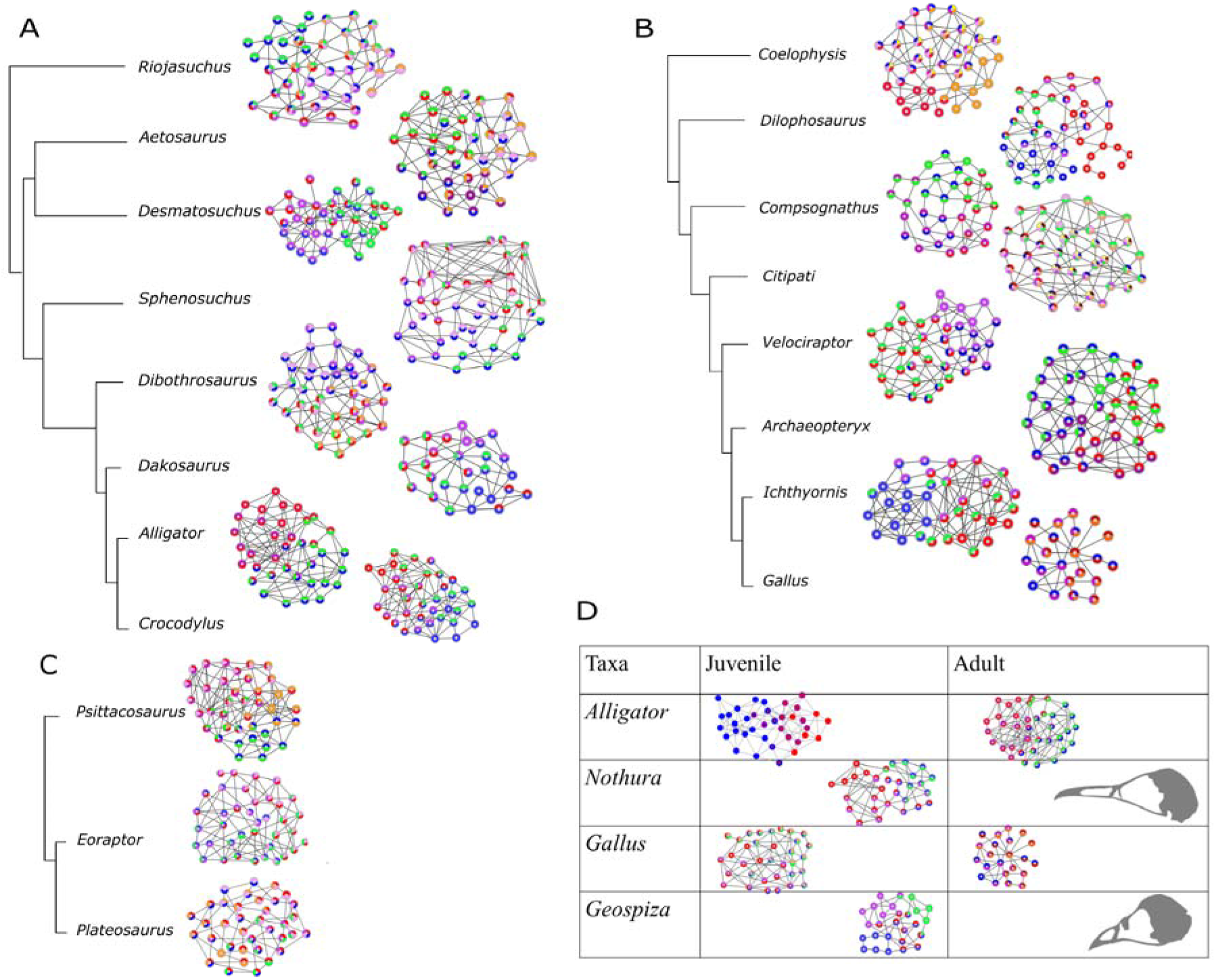
Visualizations of the module composition changes across phylogeny. The number of node-based modules ranged from 1 to 8. (C) shows the difference in module composition among the ornithischian *Psittacosaurus,* the basal saurischian *Eoraptor,* and the sauropodomorph *Plateosaurus.* (D) Comparisons of the adult and juvenile stages of extant species. Adult *Nothura* and *Geospiza* are shaded in grey as one module was identified because of the small number of nodes and links due to a highly fused skull. Nodes were colored based on their modules. Composition of each module is listed in Supplementary Table 4.

## DISCUSSION

### Occupation of morphospace and evolution of skull architecture

The two major groups of archosaurs (Crurotarsi and Avemetatarsalia) show an analogous trend towards a reduction in the number of skull bones (Table S8; Supplementary Information 3), in line with the Williston’s Law, which states that vertebrate skulls tend to become more specialized with fewer bones as a result of fusions of neighboring bones during evolution^25,56,57^.This reduction in the number of bones and articulations, together with an increase in density, is also observed within aetosaurs and sphenosuchians (Table S8). Likewise, we observed fusion of paired bones into new unpaired ones: for example, left and right frontals, parietals, and palatines are fused through their midline suture in the more derived taxa, such as the crocodilians (Table S6). Bone fusion in extant species produced skulls that are more densely connected than the skulls of extinct species (Fig. S1C). It was previously suggested that the more connected skulls would have more developmental and functional inter-dependences among bones, and, hence, they would be more evolutionarily constrained^22,23^. Similarly, avian cranium with its strongly correlated traits has lower evolutionary rates and bird skulls are less diverse overall^12^.

Bhullar et al. pointed out that avian kinesis relies on their loosely integrated skulls with less contact and, thus, skulls with highly overlapping bones would be akinetic^58^. This contradicts our observations here in that kinetic crown birds have more complex and integrated skulls than the akinetic crurotarsans and the partially kinetic *Riojasuchus^59^*. The reason could be that Bhullar et al. factored in how much connective tissue and number of contact points each bone has, but not the total number of connections possible from the number of bones in these taxa. The total number of articulations possible is the denominator used to calculate density. More recently, Werneburg and colleagues showed *Tyrannosaurus,* suspected to have kinesis, also has a higher density when compared to akinetic *Alligator* but lower density when compared to the more derived and clearly kinetic *Gallus* skull^29^.

When compared with modules identified by Felice et al.^60^, the node-based modules, such as the rostral and neurocranial modules (shown as blue and red modules in Fig. 4), are composed of elements essentially similar to those described as variational modules (more details in Supplementary Information 2). The supraoccipital and basioccipital bones were part of the same topology-defined (Supplementary Information 2, Fig. 4) and shape-defined module in most taxa, likely due to its functional importance in connecting the vertebral column with the skull^60^.

### Crurotarsi

The aetosaurs, *Aetosaurus* and *Desmatosuchus,* and the sphenosuchians, *Sphenosuchus* and *Dibothrosuchus,* show an increase in complexity within their lineages. The more derived aetosaur *Desmatosuchus* has a fused skull roof (parietal fused with supraoccipital, laterosphenoid, prootic and opisthotic) and toothless premaxilla that are absent in the less derived aetosaur *Aetosaurus*^61–63^. In contrast, basal and derived sphenosuchian are more topologically similar. Their main difference is that basipterygoid and epiotic are separate in *Sphenosuchus* but are fused with other bones in the more derived *Dibothrosuchus^64,65^*. When we compared aetosaurs and sphenosuchians, we found that sphenosuchians have a skull roof intermediately fused condition between *Aetosaurus* and *Dibothrosuchus:* interparietal sutures in both sphenosuchians are fused while supraoccipital, laterosphenoid, opisthotic, and prootic remain separate.

To understand cranial topology in Thalattosuchia, a clade with adaptations specialized for marine life, we included *Dakosaurus andiniensis.* These adaptations comprise nasal salt glands^66^, hypocercal tail, paddle-like forelimbs, ziphodont dentition, fusion of the inter-premaxillary suture, a fused vomer, and a short and high snout^67,68^. Despite these adaptations, *Dakosaurus* has a cranial complexity closer to that of extant crocodilians by similarly having inter-frontal and interparietal fusions^67,68^. In addition to the fused frontals and parietals, both *Crocodylus* and *Alligator* have a fused palate and a fused pterygoid bones.

In turn, crurotarsans first fuse the skull roof and skull base, followed by the fusion of the face (more details on Table S6). Interestingly, this resonates with the pattern of sutural fusion in alligator ontogeny, which cranial (i.e. frontoparietal) has the highest degree of suture closure followed by skull base (i.e. basioccipital-exoccipital) and then the face (i.e. internasal)^69^ suggesting that the same mechanism may control topological changes in both ontogeny and evolution.

### Avemetatarsalia

Avemetatarsalian transition is marked with a faster ontogenetic bone growth in more derived taxa, indicated by higher degree of vascularization, growth marks, and vascular canal arrangement (reviewed by Bailleul^70^), more pneumatized skulls (reviewed by Gold^71^), and an increase in complexity reminiscent of what is observed in crurotarsans. The basal ornithischian *Psittaosaurus lujiatunensis* and basal saurischian *Eoraptor lunensis* are relatively close to each other on the morphospace (Fig. 3), with the *Psittacosaurus* skull showing slightly more density because of fused palatines, a trait which is also observed in extant crocodilians and some birds, and its extra rostral bone as observed in other ceratopsians^72^.

The basal sauropodomorph *Plateosaurus engelhardti* has the lowest clustering coefficient (i.e. lower integration) of archosaurs, suggesting that skulls of sauropodmorphs are less integrated than those of saurischians^31^, accompanied by poorly connected bones (as seen in the network in Fig. 4C). Poorly connected bones, for example epipterygoid, and some connections, such as the ectopterygoid-jugal articulation, are later lost in neosauropods^43,73^.

Within theropods, the ceratosaurian *Coelophysis* is more derived and has a slightly more complex and specialized skull than the ceratosaurian *Dilophosaurus*^42^. Their positions on the morphospace suggest that ceratosaurians occupy a region characterized by a higher mean path length (L), when compared to other archosaurs (Fig. 3). *Compsognathus* is close to *Riojasuchus* on the morphospace with a similar mean path length (Figs 3 and S4, Table S8), its facial bones are also unfused, and it has a similar composition for its facial modules (see facial modules in *Compsognathus* and nasal modules in *Riojasuchus* on Table S4 and Figure S10). These observations suggest an ancestral facial topology (see Table S6 and S8 for more details) is concomitant to the magnitude of shape change reported for compsognathids^34^. *Compsognathus* possesses an independent postorbital that is absent from *Ichthyornis* to modern birds. It also has an independent prefrontal that is absent in most Oviraptorsauria and Paraves^74^, including *Citipati, Velociraptor,* and from *Ichthyornis* to modern birds. Despite its ancestral features, the back of the skull and the skull base of *Compsognathus* are fused, similarly to other Paravians and modern birds.

The oviraptorid *Citipati* has a skull topology that occupies a morphospace within non-avian theropods, despite its unique vertically-oriented premaxilla and short beak^34,75^. *Citipati* has an independent epipterygoid that is also present in some non-avian theropods and ancestral archosaurs, such as *Plateosaurus erlenbergiensis,* but which is absent in extant archosaurs^75–78^. *Citipati* also has fused skull roof (with fused interparietals), skull base, and face, marked with fused internasal and the avian-like inter-premaxillary sutures.

Like other dromaeosaurids, *Velociraptor’s* eyes are positioned lateral to the rostrum. Its prefrontal bone is either absent or fused with the lacrimal while it remains separate in other dromaeosaurids^79–81^. We observed a loss of the prefrontals from *Citipati* to modern birds, but not in more ancestral archosaurs or crurotarsans. Bones forming the *Velociraptor* basicranium, such as basioccipital, and basisphenoid are fused with other members of the basicranium (listed in Table S6). Despite having a similar number of bones and articulations to *Citipati,* the cranial bones in *Velociraptor* are more integrated with each other and are more likely to connect to bones with a different number of articulations (i.e. more disparity) (Table S8). Like *Compsognathus* and other primitive non-avian dinosaurs, *Velociraptor* has an ancestral facial topology with separate premaxilla, maxilla, and nasal bones.

### *Archaeopteryx* and *Ichthyornis* as intermediates between non-avian theropods and modern birds

The skull of *Archaeopteryx* occupied a region of the morphospace closer to non-avian dinosaurs and crurotarsans than to juvenile birds (Fig. 3). The distance of *Archaeopteryx* from crown birds and its proximity in the morphospace to *Velociraptor* and *Citipati* along the PC1 axis (Fig. 3) may reflect the evolving relationship between cranial topology and endocranial volume. In fact, *Archaeopteryx* has an endocranial volume which is intermediate between the ancestral non-avian dinosaurs and crown birds^82,83^ and it is within the plesiomorphic range of other non-avian Paraves^84^. This makes *Archaoepteryx* closer to dromaeosaurid *Velociraptor* than to oviraptor *Citipati,* for both its skull anatomy and its endocranial volume^84^. Modifications related to the smaller endocranial volume in *Archaeopteryx* include the unfused bones in the braincase, the independent reappearance of a separate prefrontal after the loss in Paraves^74^, a separate left and right premaxilla as observed in crocodilian snouts and ancestral dinosaurs, and the presence of separate postorbitals, which might restrict the fitting for a larger brain^34^.

Compared to *Archaeopteryx, Ichthyornis* is phylogenetically closer to modern birds and occupies a region of the morphospace near the juvenile birds and extant crocodilians when adult birds are included in the analysis (Fig. 3), but closer to extant crocodilians when all birds or when adult birds are removed (Figs. S4-9). The proximity between *Ichthyornis* and juvenile birds may be explained by the similar modular division (as observed in Figs. 4B and 4D; Table S4, Fig. S10), presence of anatomical features characteristic of modern birds, such as the loss of the postorbital bones, the fusion of the left and right premaxilla to form the beak, a bicondylar quadrate that form a joint with the braincase, and the arrangement of the rostrum, jugal, and quadratojugal required for a functional cranial kinetic system^58,85–88^. The proximity between *Ichthyornis* and extant crocodilians in terms of complexity (Figs. S4-9, Table S8) may be explained by the fused frontal and fused parietal, and separate maxilla, nasal, prootic and laterosphenoid (Table S6).

### Paleognath and neognath birds

Juvenile birds have a skull roof with relatively less fused bones with the interfrontal, interparietal, and frontoparietal sutures open, and a more fused skull base. Postorbital is already fused in all juvenile birds (i.e. after hatching). Collectively, juvenile neognaths show a skull anatomy with a fused cranial base, relatively less fused roof, and unfused face that resembles the anatomy of ancestral non-avian theropods. Unlike what is observed in non-avian theropods, frontal, parietal, nasal, premaxilla, and maxilla eventually fuse with the rest of the skull in adult modern birds. However, in the palatal region not all the sutures are completely closed: the caudal ends of the vomers remained unfused in adult *Nothura,* which is a characteristic common in Tinamidae^89^. A similar pattern of suture closure has been described in another paleognath, the emu, in which the sutures of the base of the skull close first and then the cranial and facial sutures close while palatal sutures remain open^69^. The only difference is that in *Nothura,* where closure of major cranial sutures (frontoparietal, interfrontal, and interparietal) happens after the facial sutures closure. In summary, when compared with neognaths, the skull of the paleognath *Nothura* is more homogeneous and complex in both juvenile and adult stages. As the skull grows, its bones fuse and both its complexity and heterogeneity increase.

Within the neognaths, the skull of *Geospiza fortis* is more complex and more homogenous than *Gallus gallus* in both juvenile and adult stages: bones in *Geospiza* skull are more likely to connect with bones with the same number of connections than *Gallus.* These two trajectories illustrate how the connectivity of each bone diversifies and becomes more specialized within a skull as sutures fuse together, as predicted by the Williston’s law.

As in crurotarsans, major transitions in Avemetatarsalia are associated with the fusion first of the skull base, then the skull roof, and, finally, with the face (more details on Table S6). This is more similar to the temporal pattern of sutural closure during ontogeny in the emu (skull base first, skull roof second, facial third) than to the one observed in the alligator (cranial first, skull base second, facial third)^69^, thus suggesting that the same mechanism for ontogeny may have been coopted in Avemetatarsalia evolution.

### Ontogenetic differences in topology between birds and crocodilians

Our comparisons on network anatomy found that juvenile birds occupy a region of the morphospace that is closer to the less derived archosaurs and crurotarsans than to that occupied by adult modern birds (Fig. S1B). Juvenile birds have a degree of anisomerism of skull bones and skull anatomical complexity closer to that in crurotarsans and non-avian dinosaurs, while their pattern of integration overlaps with that of adult birds, crurotarsans, and non-avian dinosaurs. These similarities in complexity and heterogeneity may be explained by the comparably higher number and symmetrical spatial arrangements of circumorbital ossification centres in early embryonic stages^74^. For example, both crown avians and *A. mississippiensis* have two ossification centres that fuse into one for lacrimals^74,90^. Meanwhile, ossification centres that form the prefrontal and postorbital, fuse in prenatal birds but remain separate in adult non-avian dinosaurs^74,90,91^. These ossification centres later develop into different, but overlapping, number of bones and their arrangement in juvenile birds (27 – 34 bones) and adult non-avian theropods (32 – 44 bones) with discrepancies explained by the heterochronic fusion of the ossification centres (Table S8).

Following postnatal fusions and growth, modern bird skulls become more heterogeneous and their bones more connected and topologically closer to each other (Figs. 3C and 5; Table S8). This makes avian skull bones more diverse and functionally integrated. Simultaneously, skull topology in birds diversifies with ontogeny within their lineage, as shown by the ontogenetic trajectories of *Gallus, Nothura,* and *Geospiza* (Figs. 3C and 5). Thus, bones (1) develop from ossification centres shared among crurotarsans and avemetatarsalians, (2) interact as modules with heterogeneity and complexity similar to basal members at juvenile stage, and (3) then fuse and diversify to produce skulls of adult birds.

**Figure 5.**
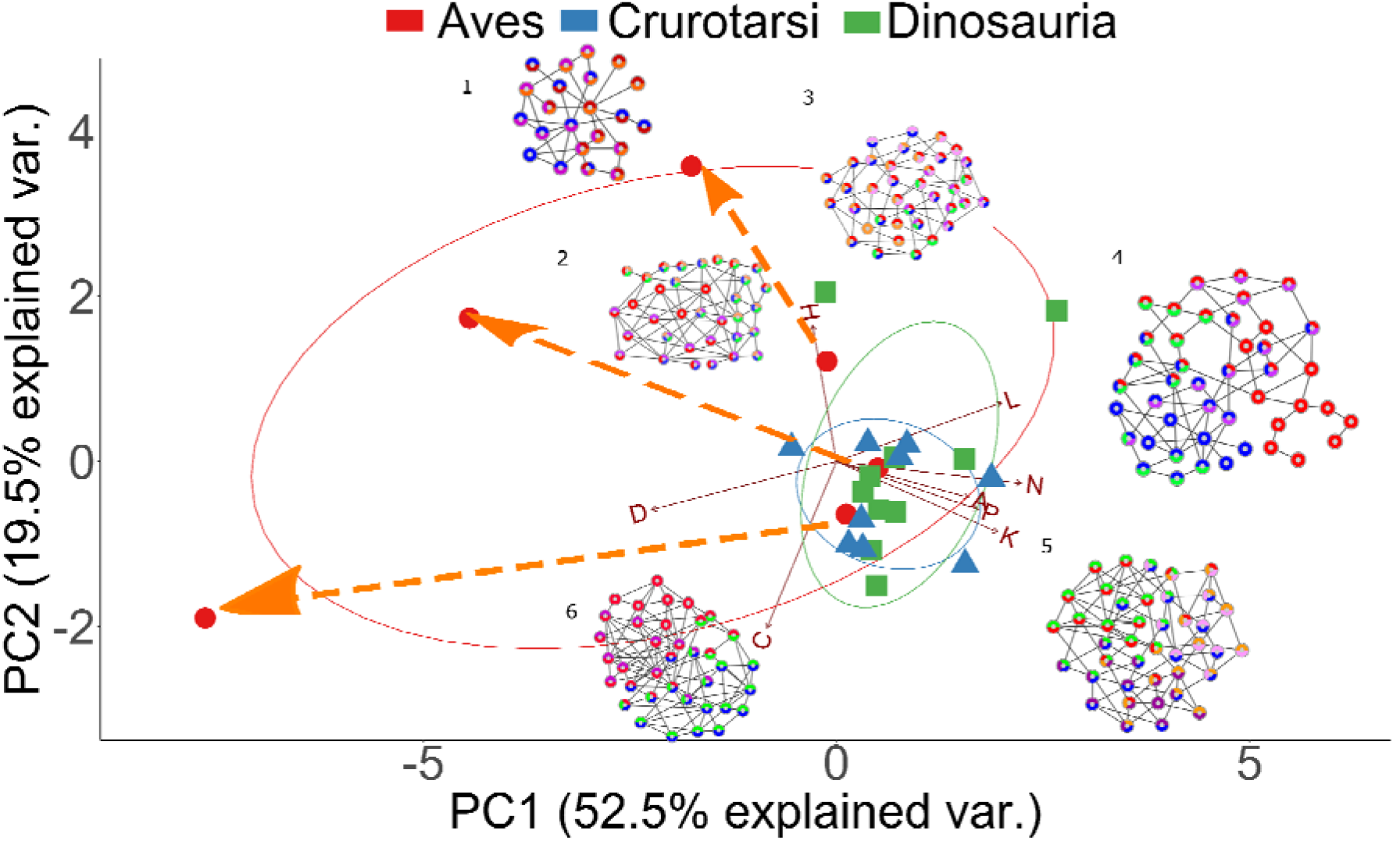
Overview of the evolution of archosaurian skull topology: Modern birds and few non-avian dinosaurs have more heterogeneous connections than crurotarsans; extant taxa have fewer bones and articulations than the extinct ones; bones in juvenile modern birds fuse and produce a more densely connected adult skull. Modules and networks of the following taxa are shown: (1) *Gallus,* (2) juvenile *Gallus,* (3) *Plateosaurus,* (4) *Dilophosaurus,* (5) *Aetosaurus,* (6) adult *Alligator.* Morphospace of Aves is significantly different from Crurotarsi and Dinosauria when adult birds are included. Orange arrows show the ontogenetic changes from juvenile to adult stages in neornithes. Taxa on the left side of the biplot have higher density and fewer bones, such as *Gallus* and *Alligator,* than taxa on the right, such as *Aetosaurus* and *Dilophosaurus*.

The skulls of birds, crocodilians, and dinosaurs develop from ossification centres with comparable spatial locations in the embryonic head^74^. When both evolutionary and ontogenetic cranial shape variation was compared among crocodilians, Morris and colleagues showed that at mid-to late embryonic stages, cranial shapes originated from a conserved region of skull shape morphospace^92^. They suggested that crocodilian skull morphogenesis at early and late embryonic stages are controlled by signaling molecules that are important in other amniotes as well, such as *Bmp4, calmodulin, Sonic hedgehog (Shh);* and *Indian hedgehog*^92–99^. Then, from late prenatal stages onward, snout of crocodilians narrows^100^ and elongates following different ontogenetic trajectories to give the full spectrum of crocodilian cranial diversity^92^.

Another major transformation in archosaurian evolution is the origin of skulls of early and modern birds from the snouted theropods. This transition involved two significant heterochronic shifts^34,101^. First, avians evolved highly paedomorphic skull shapes compared to their ancestors by developmental truncation^34^. This was followed, by a peramorphic shift where primitively paired premaxillary bones fused and the resulting beak bone elongated to occupy much of the new avian face^101^. By comparison, the skull of *Alligator* undergoes extensive morphological change and closing of the interfrontal and interparietal sutures during embryogenesis is followed by the prolonged postnatal and maturation periods, with the lack of suture closure and even widening of some sutures^102,103^. Bailleul and colleagues suggested that mechanisms that inhibit suture closure, rather than bone resorption, cause the alligator sutures to remain open during ontogeny^103^.

Nevertheless, juvenile and adult alligators share the same cranial topology featuring similar module compositions and both occupy a region of morphospace close to *Crocodylus* (Figs. 4D and S10; Table S4 and S8). Such topological arrangement suggests that conserved molecular, cellular, and developmental genetic processes underlie skull composition and topology observed across crocodilians. Likewise, oviraptorid dinosaurs, as represented by *Citipati,* display their own unique skull shape and ontogenetic transformation^34^, while retaining a topology conserved with other theropods. Combined, this evidence suggests that developmental mechanisms controlling skull composition and interaction among skull elements are conserved among theropods.

The process of osteogenesis underlies the shape and topology of the bony skull. In chicken embryo, inhibition of FGF and WNT signaling pathways prevented fusion of the suture that separates the left and right premaxilla, disconnected the premaxilla-palatine articulation and changed their shapes giving the distal face a primitive snout-like appearance^101^. The site of bone fusion in experimental unfused, snout-like chicken premaxillae showed reduced expression of skeletal markers *Runx2, Osteopontin,* and the osteogenic marker *Col I*^101^, implying localized molecular mechanisms regulating suture closure and shape of individual cranial bones. Thus, changes in gene expression during craniofacial patterning in avians^95,96,98,104–106^, non-avian dinosaurs, and crocodilians^92,101^ contribute to the clade-specific differences in skull anatomical organization resulting from the similar patterns of bone fusion of bones.

Finally, we observe some network modules where some bones within the same modules in juveniles will later fuse in adult birds, but not in *A. mississippiensis* (Supplementary Information 5; Figs. 4E and S10, Table S4). For example, in *Nothura,* premaxilla, nasal, parasphenoid, pterygoid, vomer, and maxilla grouped in the same juvenile module will later fuse during formation of the upper beak in the adult. In *A. mississippiensis,* premaxilla, maxilla, nasal, lacrimal, prefrontal, jugal, frontal, and ectopterygoid are also in the same juvenile module, but remain separate structures in adult. These findings suggest that bones within the same module may be more likely to fuse together in ontogeny but doing so is a lineage-specific feature.

Comparisons of juveniles and adults for extant birds and the alligator revealed ontogenetic changes linked to the evolution of the skull organization in archosaurs. Whereas the anatomical organization of the skull of juvenile alligators resembles that of adults, the anatomy of juvenile modern birds is closer to that of non-avian dinosaurs than to that of adult avians of the same species in terms of morphological complexity and anisomerism, probably due to the spatial arrangements of ossification centres at embryonic stages^74,90,91^. More specifically, the differences in skull organization between crown birds and non-avian dinosaurs could be explained by postnatal fusion of bones.

## CONCLUSION

A network-based comparison of the cranial anatomy of archosaurs shows that differences within and among archosaurian clades are associated with an increase of anatomical complexity, a reduction in number of bones (as predicted by the Williston’s Law), and an increase of anisomerism marked by bone fusion, for both crurotarsans and avemetatarsalians. Our findings indicate that the anatomical organization of the skull is controlled by developmental mechanisms that diversified across and within each lineage: heterotopic changes in craniofacial patterning genes, heterochronic prenatal fusion of ossification centres^74,90,91^, and lineage-specific postnatal fusion of sutures. Some of these mechanisms have been shown to be conserved in other tetrapods. For example, heterotopy of craniofacial patterning genes also took place between chick and mice embryos^95,96,106^. Hu and Marcucio showed that mouse frontonasal ectodermal zone could alter the development of the avian frontonasal process, suggesting a conserved mechanism for frontonasal development in vertebrates^96^. Our findings illustrate how a comparative analysis of the anatomical organization of the skull can reveal both common and disparate patterns and processes determining skull evolution in vertebrates.

## ACKNOWLEDGEMENT

We thank Jake Horton for coding the adult and juvenile matrices for *Alligator mississippiensis* and *Crocodylus moreletii,* Patrick Campbell of Natural History Museum London for providing reptile specimens, Alfie Gleeson and Digimorph for CT scans of crocodiles, and staff from Natural History Museum library for literature search. BE-A has received financial support through the Postdoctoral Junior Leader Fellowship Programme from “la Caixa” Banking Foundation (LCF/BQ/LI18/11630002) and also thanks the Unidad de Excelencia María de Maeztu funded by the AEI (CEX2018-000792-M). HWL’s Master Thesis that inspired this project was funded by Imperial College London and Natural History Museum, London.

## AUTHOR CONTRIBUTION

HWL, BE-A, AA designed the study.

HWL coded network models.

HWL and BE-A wrote the R scripts and performed the analyses.

All authors discussed the results and wrote the manuscript.

## Data Availability

Data and R code are available at https://figshare.com/s/80714fb9a06e886cd412.

## SUPPLEMENTARY MATERIALS

Table S1. Variance distribution across principal components

Table S2. First and last occurrence dates used to calibrate phylogenetic tree

Table S3. Internal nodes used for the phylogenetic tree

Table S4. Composition of modules for each taxon

Table S5. Categories of archosaurs based on capabilities of flight

Table S6. List of major fusion of bones with other bones in archosaurs

Table S7. Variation explained by each parameter

Table S8. Topological network parameters measured for each taxon

Table S9. Network parameters categorized by diet

Table S10. Number of modules.

Fig. S1. First two PC of topological parameters for all taxa.

Fig. S2. Second and third PC of topological parameters for all taxa.

Fig. S3. First and third PC of topological parameters for all taxa.

Fig. S4. First two PC of topological parameters for all taxa excluding avians.

Fig. S5. Second and third PC of topological parameters for all taxa excluding avians.

Fig. S6. First and third PC of topological parameters for all taxa excluding avians.

Fig. S7. First two PC of topological parameters for all taxa excluding adult avians.

Fig. S8. Second and third PC of topological parameters for all taxa excluding adult avians.

Fig. S9. First and third PC of topological parameters for all taxa excluding adult avians.

Fig. S10. Node-based modules of archosaurs based on details listed on Table S4.

Supplementary Information 1 References and notes about the specimens used.

Supplementary Information 2 Comparison between network-modules and variational modules in archosaurs.

Supplementary Information 3 Comparison of network parameters among Aves, Crurotarsi, and non-avian Dinosauria.

Supplementary Information 4 Comparison based on diet.

Supplementary Information 5 Comparison of juvenile avian modules with adult avian bones.

Supplementary Information 6 Supplementary Reference

