## Supplementary Materials for "Evolutionary and ontogenetic changes of the anatomical organization and modularity in the skull of archosaurs"

**Short title: Evolution of network anatomy in archosaurian skulls**

Hiu Wai Lee[1,2], Borja Esteve-Altava[3*], Arkhat Abzhanov[1,2*]

### Supplementary Materials

**Table S1: Variance distribution across principal components for all taxa (A), when modern birds were excluded (B), and when adult birds were excluded (C).**

| **A** | PC1 | PC2 | PC3 | PC4 | PC5 | PC6 | PC7 | PC8 |
|---|---|---|---|---|---|---|---|---|
| Standard deviation | 2.145 | 1.304 | 0.923 | 0.614 | 0.507 | 0.400 | 0.217 | 0.061 |
| Proportion of Variance | 0.575 | 0.213 | 0.106 | 0.047 | 0.032 | 0.020 | 0.006 | 0.000 |
| Cumulative Proportion | 0.575 | 0.788 | 0.894 | 0.941 | 0.974 | 0.994 | 1.000 | 1.000 |

| **B** | PC1 | PC2 | PC3 | PC4 | PC5 | PC6 | PC7 | PC8 |
|---|---|---|---|---|---|---|---|---|
| Standard deviation | 1.757 | 1.343 | 1.249 | 0.978 | 0.588 | 0.436 | 0.237 | 0.021 |
| Proportion of Variance | 0.386 | 0.226 | 0.195 | 0.120 | 0.043 | 0.024 | 0.007 | 0.000 |
| Cumulative Proportion | 0.386 | 0.611 | 0.806 | 0.926 | 0.969 | 0.993 | 1.000 | 1.000 |

| **C** | PC1 | PC2 | PC3 | PC4 | PC5 | PC6 | PC7 | PC8 |
|---|---|---|---|---|---|---|---|---|
| Standard deviation | 1.691 | 1.400 | 1.248 | 0.996 | 0.612 | 0.440 | 0.249 | 0.022 |
| Proportion of Variance | 0.358 | 0.245 | 0.195 | 0.124 | 0.047 | 0.024 | 0.008 | 0.000 |
| Cumulative Proportion | 0.358 | 0.602 | 0.797 | 0.921 | 0.968 | 0.992 | 1.000 | 1.000 |

**Table S2: First and last occurrence dates (million years ago) obtained from Paleobiology Database and used to calibrate phylogenetic tree.** Species-level occurrence dates were obtained except for *Nothura* and *Archaeopteryx.* For *Nothura* and *Archaeopteryx,* genus-level occurrence dates were used. Because *Nothura* was an extant species, a value of 0 was used for its last occurrence in the analysis. The data were downloaded from the Paleobiology Database on 1 December, 2019, using the following species names. For *Nothura* and *Archaeopteryx*, the data were downloaded using the genus names.

| Taxa Name | First occurrence | Last occurrence |
|---|---|---|
| *Riojasuchus tenuisceps* | 228 | 208.5 |
| *Aetosaurus ferratus* | 228 | 208.5 |
| *Desmatosuchus haplocerus* | 237 | 208.5 |
| *Sphenosuchus acutus* | 201.3 | 190.8 |
| *Dibothrosuchus elaphros* | 199.3 | 190.8 |

| | | |
|---|---|---|
| *Dakosaurus andiniensis* | 152.1 | 139.8 |
| *Alligator mississippiensis* | 66 | 0 |
| *Crocodylus moreletii* | 0.126 | 0 |
| *Psittacosaurus lujiatunensis* | 125.45 | 122.46 |
| *Eoraptor lunensis* | 237 | 228 |
| *Plateosaurus engelhardti* | 228 | 208.5 |
| *Coelophysis bauri* | 228 | 201.3 |
| *Dilophosaurus wetherilli* | 199.3 | 182.7 |
| *Compsognathus longipes* | 157.3 | 145 |
| *Citipati osmolskae* | 83.6 | 72.1 |
| *Velociraptor mongoliensis* | 100.5 | 66 |
| *Archaeopteryx* | 152.1 | 125.45 |
| *Ichthyornis dispar* | 89.8 | 70.6 |
| *Nothura* | 6.8 | 0.012 |
| *Gallus gallus* | 0.126 | 0 |
| *Geospiza fortis* | 0.0117 | 0 |

**Table S3: Internal nodes used to calibrate the phylogenetic tree and were based on fossil dates from Benton and Donoghue[1].**

| Internal nodes | Date (Million years ago) |
|---|---|
| Ornithodira-Crurotarsi-Avemetatarsalia | 250.4 |
| Galloanserae-Neoaves/Neognathae | 86.5 |
| Palaeognathae-Neognathae/Neornithes | 86.5 |

**Table S4: Composition of modules and their corresponding *p* values for each taxon, listed accordingly to phylogenetic tree (Fig. 2).** Color of modules used correspond to color of modules in Fig. 4. Red modules include the left supraoccipital or supraorbital. Blue modules include the left premaxilla.

| | Module | *p* value | Bones grouped in each module |
|---|---|---|---|
| *Riojasuchus* | Quadrate | 9.55E-10 | Basioccipital, Basisphenoid, L.Ectopterygoid, L.Jugal, L.Opisthotic, L.Palatine, L.Parietal, L.Postfrontal, L.Postorbital, L.Pterygoid, L.Quadrate, L.Quadratojugal, L.Squamosal, L.Vomer, R.Ectopterygoid, R.Jugal, R.Opisthotic, R.Palatine, R.Parietal, R.Postfrontal, R.Postorbital, R.Pterygoid, R.Quadrate, R.Quadratojugal, R.Squamosal, R.Vomer, Supraoccipital |
| | Snout | 1.85E-10 | Basioccipital, Basisphenoid, L.Ectopterygoid, L.Jugal, L.Lacrimal, L.Maxilla, L.Nasal, L.Palatine, L.Postorbital, L.Prefrontal, L.Premaxilla, L.Pterygoid, L.Quadrate, L.Quadratojugal, L.Squamosal, L.Vomer, R.Ectopterygoid, R.Jugal, R.Lacrimal, R.Maxilla, R.Nasal, R.Palatine, R.Postorbital, R.Prefrontal, R.Premaxilla, R.Pterygoid, R.Quadrate, R.Quadratojugal, R.Squamosal, R.Vomer |

| | | | |
|---|---|---|---|
| | R. Parietal | 1.43E-06 | Basioccipital, L.Opisthotic, L.Parietal, R.Frontal, R.Jugal, R.Lacrimal, R.Laterosphenoid, R.Opisthotic, R.Parietal, R.Postfrontal, R.Postorbital, R.Prefrontal, R.Prootic, R.Quadrate, R.Quadratojugal, R.Squamosal, Supraoccipital |
| | L. Parietal | 1.43E-06 | Basioccipital, L.Frontal, L.Jugal, L.Lacrimal, L.Laterosphenoid, L.Opisthotic, L.Parietal, L.Postfrontal, L.Postorbital, L.Prefrontal, L.Prootic, L.Quadrate, L.Quadratojugal, L.Squamosal, R.Opisthotic, R.Parietal, Supraoccipital |
| | R. Nasal | 2.39E-08 | L.Jugal, L.Lacrimal, L.Maxilla, L.Nasal, L.Prefrontal, L.Premaxilla, R.Frontal, R.Jugal, R.Lacrimal, R.Laterosphenoid, R.Maxilla, R.Nasal, R.Opisthotic, R.Parietal, R.Postfrontal, R.Postorbital, R.Prefrontal, R.Premaxilla, R.Prootic, R.Quadrate, R.Quadratojugal, R.Squamosal |
| | L Nasal | 2.39E-08 | L.Frontal, L.Jugal, L.Lacrimal, L.Laterosphenoid, L.Maxilla, L.Nasal, L.Opisthotic, L.Parietal, L.Postfrontal, L.Postorbital, L.Prefrontal, L.Premaxilla, L.Prootic, L.Quadrate, L.Quadratojugal, L.Squamosal, R.Jugal, R.Lacrimal, R.Maxilla, R.Nasal, R.Prefrontal, R.Premaxilla |
| *Aetosaurus* | Module | *p* value | Bones grouped in each module |
| | Palatine | 6.30E-08 | Basioccipital, Basisphenoid, L.Ectopterygoid, L.Exoccipital, L.Maxilla, L.Opisthotic, L.Palatine, L.Premaxilla, L.Pterygoid, L.Quadrate, L.Vomer, R.Ectopterygoid, R.Exoccipital, R.Maxilla, R.Opisthotic, R.Palatine, R.Premaxilla, R.Pterygoid, R.Quadrate, R.Vomer, Supraorbital |
| | Quadrate | 1.24E-06 | Basioccipital, Basisphenoid, L.Ectopterygoid, L.Exoccipital, L.Opisthotic, L.Pterygoid, L.Quadrate, L.Quadratojugal, L.Squamosal, R.Ectopterygoid, R.Exoccipital, R.Opisthotic, R.Pterygoid, R.Quadrate, R.Quadratojugal, R.Squamosal, Supraorbital |
| | Parietal | 7.04E-06 | L.Frontal, L.Jugal, L.Parietal, L.Postfrontal, L.Postorbital, L.Quadrate, L.Quadratojugal, L.Squamosal, R.Frontal, R.Jugal, R.Parietal, R.Postfrontal, R.Postorbital, R.Quadrate, R.Quadratojugal, R.Squamosal, Supraorbital |
| | R.Maxilla | 2.11E-06 | R.Frontal, R.Jugal, R.Lacrimal, R.Maxilla, R.Nasal, R.Palatine, R.Parietal, R.Postfrontal, R.Postorbital, R.Prefrontal, R.Premaxilla, R.Quadrate, R.Quadratojugal, R.Squamosal, R.Vomer |
| | L.Maxilla | 2.11E-06 | L.Frontal, L.Jugal, L.Lacrimal, L.Maxilla, L.Nasal, L.Palatine, L.Parietal, L.Postfrontal, L.Postorbital, L.Prefrontal, L.Premaxilla, L.Quadrate, L.Quadratojugal, L.Squamosal, L.Vomer |
| | Premaxilla | 3.62E-07 | L.Frontal, L.Jugal, L.Lacrimal, L.Maxilla, L.Nasal, L.Palatine, L.Prefrontal, L.Premaxilla, L.Vomer, R.Frontal, R.Jugal, R.Lacrimal, R.Maxilla, R.Nasal, R.Palatine, R.Prefrontal, R.Premaxilla, R.Vomer |
| *De sm* | Module | *p* value | Bones grouped in each module |

| | | | |
|---|---|---|---|
| | L.parietal | 4.52E-08 | Basioccipital, Basisphenoid, L.Ectopterygoid, L.Frontal, L.Jugal, L.Lacrimal, L.Maxilla, L.Nasal, L.Palatine, L.Parietal, L.Postfrontal, L.Postorbital, L.Prefrontal, L.Premaxilla, L.Presphenoid, L.Pterygoid, L.Quadrate, L.Quadratojugal, L.Squamosal, L.Vomer |
| | R.parietal | 4.52E-08 | Basioccipital, Basisphenoid, R.Ectopterygoid, R.Frontal, R.Jugal, R.Lacrimal, R.Maxilla, R.Nasal, R.Palatine, R.Parietal, R.Postfrontal, R.Postorbital, R.Prefrontal, R.Premaxilla, R.Presphenoid, R.Pterygoid, R.Quadrate, R.Quadratojugal, R.Squamosal, R.Vomer |
| | Basicranium | 0.000372 | Basioccipital, Basisphenoid, L.Palatine, L.Parietal, L.Presphenoid, L.Pterygoid, L.Quadrate, L.Quadratojugal, L.Squamosal, R.Palatine, R.Parietal, R.Presphenoid, R.Pterygoid, R.Quadrate, R.Quadratojugal, R.Squamosal |
| | Vomer | 5.73E-07 | L.Ectopterygoid, L.Jugal, L.Lacrimal, L.Maxilla, L.Nasal, L.Palatine, L.Prefrontal, L.Premaxilla, L.Pterygoid, L.Vomer, R.Ectopterygoid, R.Jugal, R.Lacrimal, R.Maxilla, R.Nasal, R.Palatine, R.Prefrontal, R.Premaxilla, R.Pterygoid, R.Vomer |
| *Sphenosuchus* | Module | *p* value | Bones grouped in each module |
| | L.prefrontal | 9.69E-07 | L.Ectopterygoid, L.Epiotic, L.Frontal, L.Jugal, L.Lacrimal, L.Laterosphenoid, L.Maxilla, L.Nasal, L.Paroccipital, L.Postorbital, L.Prefrontal, L.Premaxilla, L.Prootic, L.Quadrate, L.Quadratojugal, L.Squamosal, Parietal, R.Epiotic, Supraoccipital |
| | R.prefrontal | 9.69E-07 | L.Epiotic, Parietal, R.Ectopterygoid, R.Epiotic, R.Frontal, R.Jugal, R.Lacrimal, R.Laterosphenoid, R.Maxilla, R.Nasal, R.Paroccipital, R.Postorbital, R.Prefrontal, R.Premaxilla, R.Prootic, R.Quadrate, R.Quadratojugal, R.Squamosal, Supraoccipital |
| | Basicranium | 8.65E-10 | Basioccipital, Basisphenoid, L.Basipterygoid, L.Epiotic, L.Frontal, L.Laterosphenoid, L.Paroccipital, L.Postorbital, L.Prootic, L.Quadrate, L.Quadratojugal, L.Squamosal, Parietal, R.Basipterygoid, R.Epiotic, R.Frontal, R.Laterosphenoid, R.Paroccipital, R.Postorbital, R.Prootic, R.Quadrate, R.Quadratojugal, R.Squamosal, Supraoccipital |
| | Pterygoid | 5.28E-09 | Basioccipital, Basisphenoid, L.Basipterygoid, L.Ectopterygoid, L.Epiotic, L.Epipterygoid, L.Palatine, L.Paroccipital, L.Prootic, L.Pterygoid, L.Quadrate, L.Quadratojugal, L.Squamosal, L.Vomer, R.Basipterygoid, R.Ectopterygoid, R.Epiotic, R.Epipterygoid, R.Palatine, R.Paroccipital, R.Prootic, R.Pterygoid, R.Quadrate, R.Quadratojugal, R.Squamosal, R.Vomer, Supraoccipital |
| | Snout | 3.29E-08 | L.Ectopterygoid, L.Epipterygoid, L.Jugal, L.Lacrimal, L.Maxilla, L.Nasal, L.Palatine, L.Prefrontal, L.Premaxilla, L.Pterygoid, L.Quadratojugal, L.Vomer, R.Ectopterygoid, R.Epipterygoid, R.Jugal, |

| | | | |
|---|---|---|---|
| | | | R.Lacrimal, R.Maxilla, R.Nasal, R.Palatine, R.Prefrontal, R.Premaxilla, R.Pterygoid, R.Quadratojugal, R.Vomer |
| *Dibothrosuchus* | Module | *p* value | Bones grouped in each module |
| | L.Postorbital | 9.11E-05 | Basioccipital, Basisphenoid, L.Ectopterygoid, L.Frontal, L.Jugal, L.Laterosphenoid, L.Paroccipital, L.Postorbital, L.Prootic, L.Pterygoid, L.Quadrate, L.Quadratojugal, L.Squamosal, Parietal, R.Paroccipital, Supraoccipital |
| | R.Postorbital | 9.11E-05 | Basioccipital, Basisphenoid, L.Paroccipital, Parietal, R.Ectopterygoid, R.Frontal, R.Jugal, R.Laterosphenoid, R.Paroccipital, R.Postorbital, R.Prootic, R.Pterygoid, R.Quadrate, R.Quadratojugal, R.Squamosal, Supraoccipital |
| | Basicranium | 1.06E-07 | Basioccipital, Basisphenoid, L.Frontal, L.Laterosphenoid, L.Paroccipital, L.Postorbital, L.Prootic, L.Quadrate, L.Quadratojugal, L.Squamosal, Parietal, R.Frontal, R.Laterosphenoid, R.Paroccipital, R.Postorbital, R.Prootic, R.Quadrate, R.Quadratojugal, R.Squamosal, Supraoccipital |
| | R.Nasal | 9.31E-07 | L.Premaxilla, R.Ectopterygoid, R.Frontal, R.Jugal, R.Lacrimal, R.Laterosphenoid, R.Maxilla, R.Nasal, R.Palatine, R.Postorbital, R.Prefrontal, R.Premaxilla, R.Prootic, R.Pterygoid, R.Quadrate, R.Quadratojugal, R.Squamosal, R.Vomer |
| | L.Nasal | 9.31E-07 | L.Ectopterygoid, L.Frontal, L.Jugal, L.Lacrimal, L.Laterosphenoid, L.Maxilla, L.Nasal, L.Palatine, L.Postorbital, L.Prefrontal, L.Premaxilla, L.Prootic, L.Pterygoid, L.Quadrate, L.Quadratojugal, L.Squamosal, L.Vomer, R.Premaxilla |
| | Snout | 3.42E-08 | L.Ectopterygoid, L.Jugal, L.Lacrimal, L.Maxilla, L.Nasal, L.Palatine, L.Prefrontal, L.Premaxilla, L.Pterygoid, L.Quadratojugal, L.Vomer, R.Ectopterygoid, R.Jugal, R.Lacrimal, R.Maxilla, R.Nasal, R.Palatine, R.Prefrontal, R.Premaxilla, R.Pterygoid, R.Quadratojugal, R.Vomer |
| *Dakosaurus* | Module | *p* value | Bones grouped in each module |
| | Exoccipital | 6.24E-06 | Basioccipital, Basisphenoid,Frontoparietal, L.Exoccipital, L.Jugal, L.Postorbital, L.Quadrate, L.Quadratojugal, L.Squamosal, R.Exoccipital, R.Jugal, R.Postorbital, R.Quadrate, R.Quadratojugal, R.Squamosal, Supraorbital |
| | Frontoparietal | 1.62E-05 | Frontoparietal, L.Exoccipital, L.Jugal, L.Lacrimal, L.Nasal, L.Postorbital, L.Prefrontal, L.Quadrate, L.Quadratojugal, L.Squamosal, R.Nasal, R.Prefrontal, Supraorbital |
| | Basisphenoid | 1.95E-05 | Basioccipital, Basisphenoid, L.Exoccipital, L.Jugal, L.Palatine, L.Postorbital, L.Quadrate, L.Quadratojugal, L.Squamosal, Pterygoid, R.Exoccipital, R.Palatine, Supraorbital |

| | | | |
|---|---|---|---|
| | Nasal | 4.06E-07 | Basisphenoid,Frontoparietal, L.Jugal, L.Lacrimal, L.Maxilla, L.Nasal, L.Palatine, L.Postorbital, L.Prefrontal, L.Quadratojugal, Premaxilla, Pterygoid, R.Jugal, R.Lacrimal, R.Maxilla, R.Nasal, R.Palatine, R.Postorbital, R.Prefrontal, R.Quadratojugal |
| Adult *Alligator* | Module | *p* value | Bones grouped in each module |
| | Pterygoid | 0.001381 | Basisphenoid, L.Laterosphenoid, L.Prootic, L.Quadrate, L.Vomer, Palatine, Parietal, Pterygoid, R.Laterosphenoid, R.Prootic, R.Quadrate, R.Vomer |
| | Postorbit al | 3.37E-09 | Basioccipital, Basisphenoid,Frontal, L.Ectopterygoid, L.Jugal, L.Laterosphenoid, L.Otoccipital, L.Postorbital, L.Prootic, L.Quadrate, L.Quadratojugal, L.Squamosal, Parietal, Pterygoid, R.Ectopterygoid, R.Jugal, R.Laterosphenoid, R.Otoccipital, R.Postorbital, R.Prootic, R.Quadrate, R.Quadratojugal, R.Squamosal, Supraocciptal |
| | Frontal | 3.50E-07 | Frontal, L.Ectopterygoid, L.Jugal, L.Lacrimal, L.Maxilla, L.Nasal, L.Postorbital, L.Prefrontal, L.Premaxilla, L.Quadratojugal, L.Vomer, Palatine, R.Ectopterygoid, R.Jugal, R.Lacrimal, R.Maxilla, R.Nasal, R.Prefrontal, R.Premaxilla, R.Vomer |
| | Jugal | 1.39E-06 | Frontal, L.Lacrimal, L.Maxilla, L.Nasal, L.Prefrontal, L.Premaxilla, L.Vomer, Palatine, R.Ectopterygoid, R.Jugal, R.Lacrimal, R.Maxilla, R.Nasal, R.Postorbital, R.Prefrontal, R.Premaxilla, R.Quadratojugal, R.Vomer |
| Juvenile *Alligator* | Module | *p* value | Bones grouped in each module |
| | Pterygoid | 0.00147 | Basisphenoid, L.Laterosphenoid, L.Prootic, L.Quadrate, L.Vomer, Palatine, Parietal, Pterygoid, R.Laterosphenoid, R.Prootic, R.Quadrate, R.Vomer |
| | Snout | 4.81E-08 | Frontal, L.Ectopterygoid, L.Jugal, L.Lacrimal, L.Maxilla, L.Nasal, L.Postorbital, L.Prefrontal, L.Premaxilla, L.Quadratojugal, L.Vomer, Palatine, R.Ectopterygoid, R.Jugal, R.Lacrimal, R.Maxilla, R.Nasal, R.Postorbital, R.Prefrontal, R.Premaxilla, R.Quadratojugal, R.Vomer |
| | Neurocra nium | 1.22E-07 | Basioccipital, Basisphenoid, L.Laterosphenoid, L.Otoccipital, L.Postorbital, L.Prootic, L.Quadrate, L.Quadratojugal, L.Squamosal, Parietal, Pterygoid, R.Laterosphenoid, R.Otoccipital, R.Postorbital, R.Prootic, R.Quadrate, R.Quadratojugal, R.Squamosal, Supraocciptal |
| Cr oc | Module | *p* value | Bones grouped in each module |

| | | | |
|---|---|---|---|
| | Pterygoid | 0.001374 | Basisphenoid, L.Laterosphenoid, L.Prootic, L.Quadrate, L.Quadratojugal, L.Vomer, Palatine, Pterygoid, R.Laterosphenoid, R.Prootic, R.Quadrate, R.Quadratojugal, R.Vomer |
| | Frontal | 0.000341 | Frontal, L.Lacrimal, L.Laterosphenoid, L.Nasal, L.Postorbital, L.Prefrontal, L.Squamosal, Parietal, R.Lacrimal, R.Laterosphenoid, R.Nasal, R.Postorbital, R.Prefrontal, R.Squamosal, Supraocciptal |
| | Postorbital | 4.60E-09 | Basioccipital, Basisphenoid, L.Ectopterygoid, L.Jugal, L.Laterosphenoid, L.Otoccipital, L.Postorbital, L.Prootic, L.Quadrate, L.Quadratojugal, L.Squamosal, Parietal, Pterygoid, R.Ectopterygoid, R.Jugal, R.Laterosphenoid, R.Otoccipital, R.Postorbital, R.Prootic, R.Quadrate, R.Quadratojugal, R.Squamosal, Supraocciptal |
| | Vomer | 3.84E-08 | Frontal, L.Ectopterygoid, L.Jugal, L.Lacrimal, L.Maxilla, L.Nasal, L.Postorbital, L.Prefrontal, L.Premaxilla, L.Quadratojugal, L.Vomer, Palatine, R.Ectopterygoid, R.Jugal, R.Lacrimal, R.Maxilla, R.Nasal, R.Postorbital, R.Prefrontal, R.Premaxilla, R.Quadratojugal, R.Vomer |
| *Psittacosaurus* | Module | *p* value | Bones grouped in each module |
| | Nasal | 0.0005094 | L.Frontal, L.Lacrimal, L.Nasal, L.Prefrontal, L.Premaxilla, Maxilla, R.Frontal, R.Lacrimal, R.Nasal, R.Prefrontal, R.Premaxilla, Rostral |
| | Maxilla | 2.74E-06 | L.Ectopterygoid, L.Jugal, L.Lacrimal, L.Pterygoid, L.Quadrate, L.Quadratojugal, Maxilla, Palatine, Parasphenoid, R.Ectopterygoid, R.Jugal, R.Lacrimal, R.Pterygoid, R.Quadrate, R.Quadratojugal, Vomer |
| | Frontal | 3.97E-05 | L.Frontal, L.Lacrimal, L.Laterosphenoid, L.Nasal, L.Postorbital, L.Prefrontal, L.Premaxilla, Parasphenoid, Parietal, R.Frontal, R.Laterosphenoid, R.Nasal, R.Postorbital, R.Prefrontal, R.Premaxilla, Rostral |
| | Basicranium | 2.44E-10 | Basioccipital, Basisphenoid, L.Ectopterygoid, L.Frontal, L.Jugal, L.Laterosphenoid, L.Paroccipital.process, L.Postorbital, L.Prootic, L.Pterygoid, L.Quadrate, L.Quadratojugal, L.Squamosal, Parasphenoid, Parietal, R.Ectopterygoid, R.Frontal, R.Jugal, R.Laterosphenoid, R.Paroccipital.process, R.Postorbital, R.Prootic, R.Pterygoid, R.Quadrate, R.Quadratojugal, R.Squamosal, Supraoccipital |
| | R.QJ | 9.63E-06 | Basioccipital, Basisphenoid, L.Paroccipital.process, L.Prootic, L.Squamosal, Parietal, R.Frontal, R.Jugal, R.Laterosphenoid, R.Paroccipital.process, R.Postorbital, R.Prootic, R.Quadrate, R.Quadratojugal, R.Squamosal, Supraoccipital |

| | Module | p value | Bones grouped in each module |
|---|---|---|---|
| | L.QJ | 9.63E-06 | Basioccipital, Basisphenoid, L.Frontal, L.Jugal, L.Laterosphenoid, L.Paroccipital.process, L.Postorbital, L.Prootic, L.Quadrate, L.Quadratojugal, L.Squamosal, Parietal, R.Paroccipital.process, R.Prootic, R.Squamosal, Supraoccipital |
| *Eoraptor* | Module | *p* value | Bones grouped in each module |
| | Vomer | 4.33E-08 | Basisphenoid, L.Ectopterygoid, L.Jugal, L.Lacrimal, L.Maxilla, L.Nasal, L.Palatine, L.Prefrontal, L.Premaxilla, L.Pterygoid, R.Ectopterygoid, R.Jugal, R.Lacrimal, R.Maxilla, R.Nasal, R.Palatine, R.Prefrontal, R.Premaxilla, R.Pterygoid, Vomer |
| | Pterygoid | 2.81E-08 | Basioccipital, Basisphenoid, L.Ectopterygoid, L.Jugal, L.Laterosphenoid, L.Parietal, L.Paroccipital.process, L.Postorbital, L.Pterygoid, L.Quadrate, L.Quadratojugal, L.Squamosal, R.Ectopterygoid, R.Jugal, R.Laterosphenoid, R.Parietal, R.Paroccipital.process, R.Postorbital, R.Pterygoid, R.Quadrate, R.Quadratojugal, R.Squamosal, Supraoccipital |
| | R.jugal | 3.74E-08 | L.Maxilla, L.Nasal, L.Premaxilla, R.Ectopterygoid, R.Frontal, R.Jugal, R.Lacrimal, R.Laterosphenoid, R.Maxilla, R.Nasal, R.Palatine, R.Parietal, R.Paroccipital.process, R.Postorbital, R.Prefrontal, R.Premaxilla, R.Pterygoid, R.Quadrate, R.Quadratojugal, R.Squamosal, Supraoccipital, Vomer |
| | L.jugal | 3.74E-08 | L.Ectopterygoid, L.Frontal, L.Jugal, L.Lacrimal, L.Laterosphenoid, L.Maxilla, L.Nasal, L.Palatine, L.Parietal, L.Paroccipital.process, L.Postorbital, L.Prefrontal, L.Premaxilla, L.Pterygoid, L.Quadrate, L.Quadratojugal, L.Squamosal, R.Maxilla, R.Nasal, R.Premaxilla, Supraoccipital, Vomer |
| | Frontal | 3.56E-08 | Basioccipital, L.Frontal, L.Lacrimal, L.Laterosphenoid, L.Nasal, L.Parietal, L.Paroccipital.process, L.Postorbital, L.Prefrontal, L.Quadrate, L.Quadratojugal, L.Squamosal, R.Frontal, R.Lacrimal, R.Laterosphenoid, R.Nasal, R.Parietal, R.Paroccipital.process, R.Postorbital, R.Prefrontal, R.Quadrate, R.Quadratojugal, R.Squamosal, Supraoccipital |
| *Plateosaurus* | Module | *p* value | Bones grouped in each module |
| | Squamosal | 8.42E-13 | Basioccipital, Basisphenoid, L.Ectopterygoid, L.Frontal, L.Jugal, L.Lacrimal, L.Maxilla, L.Opisthotic, L.Palatine, L.Parietal, L.Postorbital, L.Prefrontal, L.Pterygoid, L.Quadrate, L.Quadratojugal, L.Squamosal, L.Vomer, R.Ectopterygoid, R.Frontal, R.Jugal, R.Lacrimal, R.Maxilla, R.Opisthotic, R.Palatine, R.Parietal, R.Postorbital, R.Prefrontal, R.Pterygoid, R.Quadrate, R.Quadratojugal, R.Squamosal, R.Vomer, Supraoccipital |

| | | | |
|---|---|---|---|
| | Frontal | 7.81E-07 | L.Frontal, L.Lacrimal, L.Nasal, L.Opisthotic, L.Parietal, L.Postorbital, L.Prefrontal, L.Squamosal, R.Frontal, R.Lacrimal, R.Nasal, R.Opisthotic, R.Parietal, R.Postorbital, R.Prefrontal, R.Squamosal, Supraoccipital |
| | vomer | 0.01467 | Basisphenoid, L.Ectopterygoid, L.Palatine, L.Postorbital, L.Pterygoid, L.Vomer, R.Ectopterygoid, R.Palatine, R.Postorbital, R.Pterygoid, R.Vomer |
| | basisphenoid | 0.006904 | Basioccipital, Basisphenoid, L.Ectopterygoid, L.Jugal, L.Pterygoid, L.Quadrate, L.Quadratojugal, R.Ectopterygoid, R.Jugal, R.Pterygoid, R.Quadrate, R.Quadratojugal |
| | R.Snout | 3.30E-08 | L.Maxilla, L.Nasal, L.Palatine, L.Premaxilla, L.Vomer, R.Ectopterygoid, R.Frontal, R.Jugal, R.Lacrimal, R.Maxilla, R.Nasal, R.Palatine, R.Parietal, R.Postorbital, R.Prefrontal, R.Premaxilla, R.Pterygoid, R.Quadrate, R.Quadratojugal, R.Squamosal, R.Vomer |
| | L.Snout | 3.30E-08 | L.Ectopterygoid, L.Frontal, L.Jugal, L.Lacrimal, L.Maxilla, L.Nasal, L.Palatine, L.Parietal, L.Postorbital, L.Prefrontal, L.Premaxilla, L.Pterygoid, L.Quadrate, L.Quadratojugal, L.Squamosal, L.Vomer, R.Maxilla, R.Nasal, R.Palatine, R.Premaxilla, R.Vomer |
| *Coelophysis* | Module | *p* value | Bones grouped in each module |
| | vomer | 0.0002621 | Basisphenoid, L.Ectopterygoid, L.Jugal, L.Lacrimal, L.Maxilla, L.Palatine, L.Pterygoid, R.Ectopterygoid, R.Jugal, R.Lacrimal, R.Maxilla, R.Palatine, R.Pterygoid, Vomer |
| | premaxilla | 2.01E-06 | L.Ectopterygoid, L.Frontal, L.Jugal, L.Lacrimal, L.Maxilla, L.Nasal, L.Postorbital, L.Prefrontal, L.Premaxilla, R.Ectopterygoid, R.Frontal, R.Jugal, R.Lacrimal, R.Maxilla, R.Nasal, R.Postorbital, R.Prefrontal, R.Premaxilla |
| | L.ect | 3.76E-06 | Basisphenoid, L.Ectopterygoid, L.Palatine, L.Pterygoid, R.Ectopterygoid, R.Frontal, R.Jugal, R.Lacrimal, R.Maxilla, R.Nasal, R.Palatine, R.Postorbital, R.Prefrontal, R.Premaxilla, R.Pterygoid, Vomer |
| | R.ect | 3.76E-06 | Basisphenoid, L.Ectopterygoid, L.Frontal, L.Jugal, L.Lacrimal, L.Maxilla, L.Nasal, L.Palatine, L.Postorbital, L.Prefrontal, L.Premaxilla, L.Pterygoid, R.Ectopterygoid, R.Palatine, R.Pterygoid, Vomer |
| | R.Neurocranium | 3.48E-08 | Basioccipital, Basisphenoid, R.Ectopterygoid, R.Frontal, R.Jugal, R.Lacrimal, R.Laterosphenoid, R.Maxilla, R.Nasal, R.Palatine, R.Parietal, R.Paroccipital.process, R.Postorbital, R.Prefrontal, R.Premaxilla, R.Prootic, R.Pterygoid, R.Quadrate, R.Quadratojugal, R.Squamosal, Vomer |

| | | | |
|---|---|---|---|
| | L.Neurocranium | 3.48E-08 | Basioccipital, Basisphenoid, L.Ectopterygoid, L.Frontal, L.Jugal, L.Lacrimal, L.Laterosphenoid, L.Maxilla, L.Nasal, L.Palatine, L.Parietal, L.Paroccipital.process, L.Postorbital, L.Prefrontal, L.Premaxilla, L.Prootic, L.Pterygoid, L.Quadrate, L.Quadratojugal, L.Squamosal, Vomer |
| *Dilophosaurus* | Module | *p* value | Bones grouped in each module |
| | Neurocranium | 1.05E-09 | L.Ectopterygoid, L.Frontal, L.Jugal, L.Lacrimal, L.Maxilla, L.Nasal, L.Palatine, L.Postorbital, L.Prefrontal, L.Premaxilla, L.Pterygoid, L.Quadrate, L.Quadratojugal, L.Squamosal, R.Ectopterygoid, R.Frontal, R.Jugal, R.Lacrimal, R.Maxilla, R.Nasal, R.Palatine, R.Postorbital, R.Prefrontal, R.Premaxilla, R.Pterygoid, R.Quadrate, R.Quadratojugal, R.Squamosal, Vomer |
| | Snout | 1.64E-11 | Basioccipital, Basisphenoid, L.Epiotic, L.Exooccipital, L.Jugal, L.Laterosphenoid, L.Opisthotic, L.Parietal, L.Postorbital, L.Prootic, L.Quadrate, L.Quadratojugal, L.Squamosal, R.Epiotic, R.Exooccipital, R.Jugal, R.Laterosphenoid, R.Opisthotic, R.Parietal, R.Postorbital, R.Prootic, R.Quadrate, R.Quadratojugal, R.Squamosal, Supraoccipital |
| | R.Lacrimal | 0.0001292 | Basisphenoid, R.Frontal, R.Jugal, R.Lacrimal, R.Laterosphenoid, R.Nasal, R.Opisthotic, R.Parietal, R.Postorbital, R.Prefrontal, R.Prootic, R.Quadrate, R.Quadratojugal, R.Squamosal |
| | L.Lacrimal | 0.0001292 | Basisphenoid, L.Frontal, L.Jugal, L.Lacrimal, L.Laterosphenoid, L.Nasal, L.Opisthotic, L.Parietal, L.Postorbital, L.Prefrontal, L.Prootic, L.Quadrate, L.Quadratojugal, L.Squamosal |
| *Compsognathus* | Module | *p* value | Bones grouped in each module |
| | L.face | 1.30E-07 | L.Ectopterygoid, L.Frontal, L.Jugal, L.Lacrimal, L.Maxilla, L.Nasal, L.Palatine, L.Parietal, L.Postorbital, L.Prefrontal, L.Premaxilla, L.Pterygoid, L.Quadrate, L.Quadratojugal, L.Squamosal, L.Vomer, R.Palatine, R.Vomer |
| | Premaxilla | 0.0003067 | L.Ectopterygoid, L.Jugal, L.Lacrimal, L.Maxilla, L.Nasal, L.Premaxilla, R.Ectopterygoid, R.Jugal, R.Lacrimal, R.Maxilla, R.Nasal, R.Premaxilla |
| | R.face | 1.30E-07 | L.Ectopterygoid, L.Jugal, L.Lacrimal, L.Maxilla, L.Nasal, L.Premaxilla, R.Ectopterygoid, R.Jugal, R.Lacrimal, R.Maxilla, R.Nasal, R.Premaxilla |
| | Neurocranium | 0.0001686 | L.Frontal, L.Parietal, L.Postorbital, L.Prefrontal, L.Quadrate, L.Squamosal, R.Frontal, R.Parietal, R.Postorbital, R.Prefrontal, R.Quadrate, R.Squamosal |
| *Citipati* | Module | *p* value | Bones grouped in each module |
| | Supraoccipital | 0.0001169 | L.Exoccipital, L.Postfrontal, L.Quadrate, L.Quadratojugal, L.Squamosal, Parietal, R.Exoccipital, R.Postfrontal, R.Quadrate, R.Quadratojugal, R.Squamosal, Supraoccipital |

| | | | |
|---|---|---|---|
| | Parietal | 1.17E-06 | Basisphenoid, L.Basipterygoid, L.Exoccipital, L.Frontal, L.Laterosphenoid, L.Orbitosphenoid, L.Postfrontal, L.Postorbital, L.Squamosal, Parietal, R.Basipterygoid, R.Exoccipital, R.Frontal, R.Laterosphenoid, R.Orbitosphenoid, R.Postfrontal, R.Postorbital, R.Squamosal, Supraoccipital |
| | Nasal | 1.52E-05 | L.Ectopterygoid, L.Frontal, L.Jugal, L.Lacrimal, L.Maxilla, L.Postfrontal, L.Postorbital, L.Quadratojugal, L.Squamosal, Nasal, Parietal, Premaxilla, R.Frontal, R.Lacrimal, R.Postfrontal |
| | Orbitosphenoid | 0.0003628 | Basisphenoid, L.Basipterygoid, L.Frontal, L.Lacrimal, L.Laterosphenoid, L.Orbitosphenoid, L.Postorbital, Nasal, Premaxilla, R.Basipterygoid, R.Frontal, R.Lacrimal, R.Laterosphenoid, R.Orbitosphenoid, R.Postorbital |
| | R.Frontal | 4.68E-07 | Basisphenoid, L.Basipterygoid, L.Exoccipital, L.Frontal, L.Orbitosphenoid, L.Postfrontal, Nasal, Parietal, R.Basipterygoid, R.Epipterygoid, R.Exoccipital, R.Frontal, R.Jugal, R.Lacrimal, R.Laterosphenoid, R.Orbitosphenoid, R.Postfrontal, R.Postorbital, R.Pterygoid, R.Quadrate, R.Quadratojugal, R.Squamosal, Supraoccipital |
| | L.Frontal | 4.68E-07 | Basisphenoid, L.Basipterygoid, L.Epipterygoid, L.Exoccipital, L.Frontal, L.Jugal, L.Lacrimal, L.Laterosphenoid, L.Orbitosphenoid, L.Postfrontal, L.Postorbital, L.Pterygoid, L.Quadrate, L.Quadratojugal, L.Squamosal, Nasal, Parietal, R.Basipterygoid, R.Exoccipital, R.Frontal, R.Orbitosphenoid, R.Postfrontal, Supraoccipital |
| | R.jugal | 1.39E-10 | Basisphenoid, L.Basipterygoid, L.Ectopterygoid, L.Lacrimal, L.Maxilla, L.Palatine, L.Pterygoid, Nasal, Premaxilla, R.Basipterygoid, R.Ectopterygoid, R.Epipterygoid, R.Exoccipital, R.Frontal, R.Jugal, R.Lacrimal, R.Laterosphenoid, R.Maxilla, R.Orbitosphenoid, R.Palatine, R.Postfrontal, R.Postorbital, R.Pterygoid, R.Quadrate, R.Quadratojugal, R.Squamosal, Vomer |
| | L.jugal | 1.39E-10 | Basisphenoid, L.Basipterygoid, L.Ectopterygoid, L.Epipterygoid, L.Exoccipital, L.Frontal, L.Jugal, L.Lacrimal, L.Laterosphenoid, L.Maxilla, L.Orbitosphenoid, L.Palatine, L.Postfrontal, L.Postorbital, L.Pterygoid, L.Quadrate, L.Quadratojugal, L.Squamosal, Nasal, Premaxilla, R.Basipterygoid, R.Ectopterygoid, R.Lacrimal, R.Maxilla, R.Palatine, R.Pterygoid, Vomer |
| *Velociraptor* | Module | *p* value | Bones grouped in each module |
| | Nasal | 1.31E-08 | L.Ectopterygoid, L.Frontal, L.Jugal, L.Lacrimal, L.Maxilla, L.Nasal, L.Palatine, L.Postorbital, L.Premaxilla, L.Pterygoid, L.Vomer, R.Ectopterygoid, R.Frontal, R.Jugal, R.Lacrimal, R.Maxilla, R.Nasal, R.Palatine, R.Postorbital, R.Premaxilla, R.Pterygoid, R.Vomer |

| | | | |
|---|---|---|---|
| | Squamosal | 1.91E-08 | Basioccipital, Basisphenoid, L.Exoccipital, L.Frontal, L.Laterosphenoid, L.Parietal, L.Postorbital, L.Prootic, L.Quadrate, L.Quadratojugal, L.Squamosal, R.Exoccipital, R.Frontal, R.Laterosphenoid, R.Parietal, R.Postorbital, R.Prootic, R.Quadrate, R.Quadratojugal, R.Squamosal, Supraoccipital |
| | Premaxilla | 6.08E-06 | L.Ectopterygoid, L.Frontal, L.Jugal, L.Lacrimal, L.Laterosphenoid, L.Maxilla, L.Nasal, L.Palatine, L.Parietal, L.Postorbital, L.Premaxilla, L.Prootic, L.Pterygoid, L.Vomer, R.Nasal, R.Premaxilla |
| | Neurocranium | 2.01E-09 | Basioccipital, Basisphenoid, L.Exoccipital, L.Frontal, L.Lacrimal, L.Laterosphenoid, L.Nasal, L.Parietal, L.Postorbital, L.Prootic, L.Pterygoid, L.Quadrate, L.Quadratojugal, L.Squamosal, R.Exoccipital, R.Frontal, R.Lacrimal, R.Laterosphenoid, R.Nasal, R.Parietal, R.Postorbital, R.Prootic, R.Pterygoid, R.Quadrate, R.Quadratojugal, R.Squamosal, Supraoccipital |
| *Archaeopteryx* | Module | *p* value | Bones grouped in each module |
| | Pterygoid | 5.87E-09 | L.Ectopterygoid, L.Jugal, L.Lacrimal, L.Maxilla, L.Nasal, L.Palatine, L.Prefrontal, L.Premaxilla, L.Pterygoid, L.Quadrate, L.Quadratojugal, R.Ectopterygoid, R.Jugal, R.Lacrimal, R.Maxilla, R.Nasal, R.Palatine, R.Prefrontal, R.Premaxilla, R.Pterygoid, R.Quadrate, R.Quadratojugal, Vomer |
| | Neurocranium | 2.30E-07 | Basioccipital, Basisphenoid, L.Laterosphenoid, L.Parietal, L.Paroccipital.Process, L.Postorbital, L.Prootic, L.Quadrate, L.Quadratojugal, L.Squamosal, L.Supraoccipital, R.Laterosphenoid, R.Parietal, R.Paroccipital.Process, R.Postorbital, R.Prootic, R.Quadrate, R.Quadratojugal, R.Squamosal, R.Supraoccipital |
| | R.prefrontal | 1.15E-10 | Basioccipital, Basisphenoid, L.Maxilla, L.Nasal, L.Palatine, L.Premaxilla, L.Pterygoid, R.Ectopterygoid, R.Frontal, R.Jugal, R.Lacrimal, R.Laterosphenoid, R.Maxilla, R.Nasal, R.Palatine, R.Parietal, R.Paroccipital.Process, R.Postorbital, R.Prefrontal, R.Premaxilla, R.Prootic, R.Pterygoid, R.Quadrate, R.Quadratojugal, R.Squamosal, R.Supraoccipital, Vomer |
| | L.Prefrontal | 1.15E-10 | Basioccipital, Basisphenoid, L.Ectopterygoid, L.Frontal, L.Jugal, L.Lacrimal, L.Laterosphenoid, L.Maxilla, L.Nasal, L.Palatine, L.Parietal, L.Paroccipital.Process, L.Postorbital, L.Prefrontal, L.Premaxilla, L.Prootic, L.Pterygoid, L.Quadrate, L.Quadratojugal, L.Squamosal, L.Supraoccipital, R.Maxilla, R.Nasal, R.Palatine, R.Premaxilla, R.Pterygoid, Vomer |
| *Ichthyornis* | Module.number | *p* value | Bones grouped in each module |
| | L.Pterygoid | 0.009571 | Basioccipital, Basisphenoid, L.Exoccipital, L.Jugal, L.Laterosphenoid, L.Opisthotic, L.Palatine, L.Prootic, L.Pterygoid, L.Quadrate, L.Quadratojugal, L.Squamosal |

| | | | |
|---|---|---|---|
| | R.Pterygoid | 0.009571 | Basioccipital, Basisphenoid, R.Exoccipital, R.Jugal, R.Laterosphenoid, R.Opisthotic, R.Palatine, R.Prootic, R.Pterygoid, R.Quadrate, R.Quadratojugal, R.Squamosal |
| | Neurocranium | 2.76E-08 | Basioccipital, Basisphenoid, L.Epiotic, L.Exoccipital, L.Laterosphenoid, L.Opisthotic, L.Prootic, L.Pterygoid, L.Quadrate, L.Squamosal, Parietal, R.Epiotic, R.Exoccipital, R.Laterosphenoid, R.Opisthotic, R.Prootic, R.Pterygoid, R.Quadrate, R.Squamosal, Supraoccipital |
| | Maxilla | 5.75E-06 | Frontal, L.Jugal, L.Lacrimal, L.Maxilla, L.Nasal, L.Palatine, L.Quadratojugal, Mesethmoid, Premaxilla, R.Jugal, R.Lacrimal, R.Maxilla, R.Nasal, R.Palatine, R.Quadratojugal, Vomer |
| Adult [illegible] | Module | *p* value | Bones grouped in each module |
| | 1 | | Braincase, R.Jugal.Bar, L.Jugal.Bar, R.Quadrate, L.Quadrate,Upper.Beak |
| Adult *Gallus* | Module | *p* value | Bones grouped in each module |
| | Pterygopalatine | 5.29E-06 | Beak, L.Jugal, L.Palatine, L.Parasphenoid, L.Pterygoid, L.Quadrate, L.Quadratojugal, L.Vomer, R.Jugal, R.Palatine, R.Parasphenoid, R.Pterygoid, R.Quadrate, R.Quadratojugal, R.Vomer |
| | Braincase | 0.0005155 | Braincase, L.Jugal, L.Lateral.ethmoid, L.Parasphenoid, L.Postfrontal, L.Prefrontal, L.Pterygoid, L.Quadrate, L.Quadratojugal, R.Lateral.ethmoid, R.Parasphenoid, R.Postfrontal, R.Prefrontal |
| | Jugal | 0.0007588 | Braincase, L.Jugal, L.Parasphenoid, L.Postfrontal, L.Pterygoid, L.Quadrate, L.Quadratojugal, R.Jugal, R.Parasphenoid, R.Postfrontal, R.Pterygoid, R.Quadrate, R.Quadratojugal |
| | Beak | 0.0007588 | Beak, L.Jugal, L.Lateral.ethmoid, L.Palatine, L.Prefrontal, L.Vomer, R.Jugal, R.Lateral.ethmoid, R.Palatine, R.Prefrontal, R.Vomer |
| Adult *Geospiz* | Module | *p* value | Bones grouped in each module |
| | 1 | 2.71E-05 | L.Jugal.Bar, R.Jugal.Bar, R.Pterygoid, R.Quadrate,Upper.Beak, L.Pterygoid, L.Quadrate, Braincase, Palate, Parasphenoid |
| Juvenile *Nothura* | Module | *p* value | Bones grouped in each module |
| | Mesethmoid | 5.46E-06 | L.Exoccipital, L.Frontal, L.Jugal, L.Lacrimal, L.Laterosphenoid, L.Maxilla, L.Parietal, L.Quadrate, L.Quadratojugal, L.Squamosal, Mesethmoid, Nasal, Premaxilla, R.Frontal, R.Lacrimal |

| | | | |
|---|---|---|---|
| | Paraspenoid | 0.0002755 | Basioccipital, L.Maxilla, L.Pterygoid, L.Vomer, Nasal, Parasphenoid, Premaxilla, R.Maxilla, R.Pterygoid, R.Vomer |
| | Maxilla | 4.83E-06 | L.Jugal, L.Lacrimal, L.Maxilla, L.Pterygoid, L.Quadratojugal, L.Vomer, Mesethmoid, Nasal, Parasphenoid, Premaxilla, R.Jugal, R.Lacrimal, R.Maxilla, R.Pterygoid, R.Quadratojugal, R.Vomer |
| | Quadratojugal | 5.87E-09 | Basioccipital, L.Exoccipital, L.Frontal, L.Jugal, L.Lacrimal, L.Laterosphenoid, L.Parietal, L.Quadrate, L.Quadratojugal, L.Squamosal, Mesethmoid, R.Exoccipital, R.Frontal, R.Jugal, R.Lacrimal, R.Laterosphenoid, R.Parietal, R.Quadrate, R.Quadratojugal, R.Squamosal, Supraoccipital |
| Juvenile *Gallus* | Module | *p* value | Bones grouped in each module |
| | Pterygopalatine | 3.71E-06 | L.Jugal, L.Maxilla, L.Nasal, L.Prefrontal, L.Prootic, L.Pterygopalatine, L.Quadrate, L.Quadratojugal, Mesethmoid, Premaxilla, R.Jugal, R.Maxilla, R.Nasal, R.Prefrontal, R.Prootic, R.Pterygopalatine, R.Quadrate, R.Quadratojugal, Vomer |
| | Nasal | 5.83E-06 | L.Frontal, L.Jugal, L.Maxilla, L.Nasal, L.Postfrontal, L.Prefrontal, L.Pterygopalatine, L.Quadratojugal, Mesethmoid,, R.Frontal, R.Jugal, R.Maxilla, R.Nasal, R.Postfrontal, R.Prefrontal, R.Pterygopalatine, R.Quadratojugal, Vomer |
| | Jugal | 1.85E-05 | L.Jugal, L.Maxilla, L.Nasal, L.Prefrontal, L.Prootic, L.Pterygopalatine, L.Quadrate, L.Quadratojugal, Premaxilla, R.Jugal, R.Maxilla, R.Nasal, R.Prefrontal, R.Prootic, R.Pterygopalatine, R.Quadrate, R.Quadratojugal, Vomer |
| | Frontal | 3.84E-05 | L.Frontal, L.Nasal, L.Orbitosphenoid, L.Parietal, L.Postfrontal, L.Prefrontal, L.Squamosal, Mesethmoid, R.Frontal, R.Nasal, R.Orbitosphenoid, R.Parietal, R.Postfrontal, R.Prefrontal, R.Squamosal, Supraoccipital |
| | Basicranial | 1.38E-07 | Basioccipital, Basisphenoid, L.Exoccipital, L.Frontal, L.Jugal, L.Orbitosphenoid, L.Parietal, L.Postfrontal, L.Prootic, L.Quadrate, L.Quadratojugal, L.Squamosal, R.Exoccipital, R.Frontal, R.Jugal, R.Orbitosphenoid, R.Parietal, R.Postfrontal, R.Prootic, R.Quadrate, R.Quadratojugal, R.Squamosal, Supraoccipital |
| Juvenile *Geospiza* | Module | *p* value | Bones grouped in each module |
| | Braincase | 0.003688 | Basioccipital, L.Exoccipital, L.Parietal, L.Quadrate, L.Quadratojugal, R.Exoccipital, R.Parietal, R.Quadrate, R.Quadratojugal, Supraoccipital |
| | Squamosal | 0.0001178 | L.Frontal, L.Orbitosphenoid, L.Parietal, L.Squamosal, Mesethmoid, R.Frontal, R.Orbitosphenoid, R.Parietal, R.Squamosal, Supraoccipital |

| | | | |
|---|---|---|---|
| | Quadratojugal | 1.64E-05 | Basioccipital, Basisphenoid, L.Exoccipital, L.Palatine, L.Pterygoid, L.Quadrate, L.Quadratojugal, R.Exoccipital, R.Palatine, R.Pterygoid, R.Quadrate, R.Quadratojugal, Supraoccipital, Vomer |
| | Maxilla | 0.008401 | L.Maxilla, L.Quadratojugal, Mesethmoid, Nasal, Premaxilla, R.Maxilla, R.Quadratojugal |

**Table S5: Categories of archosaurs based on capabilities of flight.** *Archaeopteryx* was suggested to be capable to glide but incapable of flapping flight because it lived on arid locations full of low shrubs, had forelimb feathers similar to flightless birds and had shoulder socket different from flying birds[2]. *Nothura* and *Gallus* cannot fly for a long distance but can have burst-off flapping starts as shown by their convergently similar sternum, wing, and pectoral girdle[3]. *Ichthyornis* can fly proficiently as shown by its large humeral proximal end, and modern deltopectoral crest and sternum[3]. Other archosaurs used in this analysis cannot fly. Avialae that can do soaring flight appeared to be more affected by whether bones with the same number of articulations connect to each other (A), the number of articulations, and parcellation ($F_{3, 21} = 2.205$, $p = 0.08539$; Fig. S1).

| Taxa | Flight Category |
|---|---|
| *Gallus, Nothura* | Flapping flight |
| *Geospiza, Ichthyornis* | Soaring flight |
| *Archaeopteryx* | May be able to glide. |
| *Aetosaurus, Alligator, Citipati, Coelophysis, Compsognathus, Crocodylus, Dakosaurus, Desmatosuchus, Dibothrosuchus, Dilophosaurus, Eoraptor, Plateosaurus, Psittacosaurus, Riojasuchus, Sphenosuchus, Velociraptor* | Flightless |

**Table S6: List of major fusion of bones with other bones in archosaurs**. Bones that fused with other bones are shaded in grey. LR marked the midline fusion of the left and right bones. E marked the fusion of the bone to other bones. * marked the specimen differences in *Archaeopteryx* for prefrontals: the one reconstructed by Elzanowski[4] and Rauhut[5] did not have any prefrontals while the Eichstätt (5th) specimen was later confirmed by Smith-Paredes et al.[6] to have prefrontals. Unfused bones (shaded in light orange) were observed in archosaurs: epiotic in *Dilosphosaurus* and *Ichthyornis* and epipterygoid in *Citipati*.  For *Velociraptor*, the suture between exoccipital and supraoccipital was not clear, meaning these two bones could be fused together. BO: basioccipital, Bra: braincase, BS: basisphenoid, Epio: epiotic, Epip: epipterygoid, Exo: exoccipital, F: frontal, Lac: lacrimal, LE: lateral ethmoid, Las: laterosphenoid, M: maxilla, N: nasal, Opi: opisthotic, P: parietal, Pal: palatine, Par: parasphenoid, PO: postorbital, PreF: prefrontal, PreM: premaxilla, Pro: prootic, PostF: postfrontal, Pter: pterygoid, Sup: supraoccipital, V: vomer.

| | Pal | | V | | Sup | | BO | BS | PO | Pro | PostF | PreF |
|---|---|---|---|---|---|---|---|---|---|---|---|---|
| | E | LR | E | LR | E | LR | E | E | E | E | E | E |
| *Riojasuchus* | | | | | | | | | | | | |
| *Aetosaurus* | | | | | | | | | | | | |
| *Desmatosuchus* | | | | | Opi, P, Las, Pro | | | | | | | |
| *Sphenosuchus* | | | | | | | | | | | | |
| *Dibothrosaurus* | | | | | | | | | | | | |

| | | | | | | | | | |
|---|---|---|---|---|---|---|---|---|---|
| *Dakosaurus* | | | | | | | | | |
| *Alligator* (Adult) | | | | | | | | | |
| *Alligator* (Juv) | | | | | | | | | |
| *Crocodylus* | | | | | | | | | |
| *Psittacosaurus* | | | | | | | | | |
| *Eoraptor* | | | | | | | | | |
| *Plateosaurus* | | | | | | | | | |
| *Coelophysis* | | | | | | | | | |
| *Dilophosaurus* | | | | | | | | | |
| *Compsognathus* | | | | | | | | | |
| *Citipati* | | | | | | | | | |
| *Velociraptor* | | | Exo | Opi | Par | | | | Lac |
| *Archaeopteryx* | | | | | | | | | * |
| *Ichthyornis* | | | | | | | | | |
| Adults | | | | | | | | | |
| *Nothura* | | | | | | | | | |
| *Geospiza* | | | | | | | | | |
| *Gallus* | V | | | | | | | | |
| Juveniles | | | | | | | | | |
| *Nothura* | | | | | | | | | |
| *Geospiza* | | | | | | | | | |
| *Gallus* | V | | | | Par | | | | LE |

| | Exo | Opi | Jugal | Las | N | M | PreM | F | P | New Bones |
|---|---|---|---|---|---|---|---|---|---|---|
| | E | E | E | E | LR | LR | LR | LR | LR | |
| *Riojasuchus* | Opi | Exo | | | | | | | | |
| *Aetosaurus* | | | | | | | | | | |
| *Desmatosuchus* | | | | | | | | | | |
| *Sphenosuchus* | Opi | Exo | | | | | | | | |
| *Dibothrosuchus* | Opi | Exo | | | | | | | | |
| *Dakosaurus* | Opi | Exo | | | | | | | | |
| *Alligator* (Adult) | | | | | | | | | | |
| *Alligator* (Juv) | | | | | | | | | | |
| *Crocodylus* | | | | | | | | | | |
| *Psittacosaurus* | Opi | Exo | | | | | | | | Rostral |
| *Eoraptor* | Opi | Exo | | | | | | | | |
| *Plateosaurus* | Opi | Exo | | | | | | | | |
| *Coelophysis* | Opi | Exo | | | | | | | | |
| *Dilophosaurus* | | | | | | | | | | Epio |
| *Compsognathus* | | | | | | | | | | |
| *Citipati* | | | | | | | | | | Epip |
| *Velociraptor* | Sup | BO | | | | | | | | |
| *Archaeopteryx* | Opi | Exo | | | | | | | | |
| *Ichthyornis* | | | | | | | | | | Epio |
| Adults | | | | | | | | | | |
| *Nothura* | Bra | Bra | | | | | | | | |
| *Geospiza* | Bra | Bra | | | | | | | | |
| *Gallus* | Bra | Bra | | | | | | | | |

| |
|---|
| Juveniles |
| *Nothura* |
| *Geospiza* |
| *Gallus* |

The presence of separate opisthotic and exoccipital bones were observed in *Aetosaurus*, *Dilophosaurus*, and *Ichthyornis*; the presence of separate exoccipital bones were observed in *Citipati* and juvenile *Nothura.* Because they articulated with each other, basisphenoid and basioccipital fused to each other in *Compsognathus*, *Citipati*, and adult avians. There were no separate basisphenoid bones in juvenile avians. In *Velociraptor*, basisphenoid and basioccipital fused to other bones. No separate supraoccipital was observed in *Coelophysis*, *Compsognathus, Desmatosuchus,* modern adult birds, and juvenile *Gallus*. Supraoccipital separated into left and right bones were observed in *Archaeopteryx*.

The absence of a separate jugal bone only appeared in adult *Nothura* and the derived avian *Geospiza* (adult and juvenile). The absence of a separate postorbital only appeared from *Ichthyornis* to modern birds. Separate prootics were absent in *Aetosaurus*, *Dakosaurus*, *Desmatosuchus, Eoraptor*, *Plateosaurus*, *Compsognathus*, *Citipati*, and modern birds (except for juvenile *Gallus*). Similar trend was observed for the absence of separate laterosphenoids, except they were present in *Eoraptor, Citipati*, and juvenile *Nothura*. Separate postfrontals were present in from *Riojasuchus* to *Desmatosuchus*, *Citipati*, *Gallus*, and juvenile *Geospiza*. Separate prefrontals were absent from *Citipati* to *Nothura* and could not be separated from lateral ethmoid in juvenile *Gallus*. The juvenile prefrontal was fused to lateral ethmoid but was later separated may have been caused by the independent growth of the prefrontal ossification center[6].

Left and right palatine fused to each other in adult birds, juvenile *Nothura*, juvenile *Gallus*, extant crurotarsans, and *Psittacosaurus*. Palatine fused with either pterygoid (*Gallus*) or maxilla (*Nothura*) in modern birds. Separate vomers were not found in *Dakosaurus*. Left and right vomers fused in *Psittacosaurus*, *Eoraptor*, *Coelophysis*, *Dilophosaurus*, *Citipati*, and *Archaeopteryx*.

Right and left frontal fused to each other in *Ichthyornis*, adult avians, and from *Dakosaurus* to extant crurotarsans. Right and left parietal bones fused to each other in *Ichthyornis*, adult avians, from *Dakosaurus* to extant crurotarsans, *Psittacosaurus*, and *Citipati*. These indicated left and right fusion of palatine, frontal, and parietal are more common in extant species.

**Table S7: Variation explained by each parameter for (A) all taxa, (B) when modern birds were excluded, and (C) when adult birds were excluded.** Parameters that explained the most (in absolute values) are in bold.

| **A** | PC1 | PC2 | PC3 | PC4 | PC5 | PC6 | PC7 | PC8 |
|---|---|---|---|---|---|---|---|---|
| N | **0.44** | -0.11 | 0.26 | -0.04 | 0.01 | 0.30 | -0.25 | **-0.76** |
| K | 0.37 | -0.31 | 0.44 | -0.28 | -0.27 | 0.24 | -0.19 | **0.58** |
| D | **-0.44** | -0.20 | -0.09 | 0.02 | 0.26 | 0.10 | **-0.82** | 0.05 |
| C | -0.20 | **-0.61** | 0.33 | 0.00 | 0.57 | -0.02 | 0.39 | -0.04 |
| L | 0.39 | 0.21 | -0.25 | 0.36 | **0.53** | 0.50 | 0.03 | 0.29 |
| H | -0.10 | **0.62** | 0.44 | -0.49 | 0.41 | -0.06 | -0.04 | 0.02 |
| A | 0.31 | -0.22 | **-0.58** | **-0.66** | 0.22 | -0.20 | -0.02 | -0.02 |
| P | **0.42** | 0.00 | 0.18 | 0.35 | 0.20 | **-0.74** | -0.26 | 0.08 |

| **B** | PC1 | PC2 | PC3 | PC4 | PC5 | PC6 | PC7 | PC8 |
|---|---|---|---|---|---|---|---|---|
| N | 0.15 | **0.71** | 0.12 | -0.01 | -0.24 | -0.05 | -0.08 | **-0.62** |
| K | -0.35 | **0.54** | 0.19 | 0.00 | -0.25 | 0.18 | 0.29 | **0.61** |
| D | **-0.50** | -0.20 | 0.03 | 0.03 | 0.03 | 0.09 | **0.70** | -0.46 |
| C | -0.42 | 0.26 | 0.09 | -0.26 | **0.72** | -0.34 | -0.21 | 0.00 |
| L | **0.50** | 0.12 | -0.02 | 0.03 | 0.15 | **-0.60** | **0.57** | 0.16 |
| H | 0.05 | 0.28 | **-0.75** | 0.36 | 0.33 | 0.32 | 0.09 | 0.02 |
| A | 0.15 | 0.00 | **0.59** | **0.68** | 0.35 | 0.21 | -0.01 | -0.02 |
| P | 0.39 | 0.04 | 0.17 | **-0.58** | 0.32 | **0.57** | 0.22 | -0.01 |

| **C** | PC1 | PC2 | PC3 | PC4 | PC5 | PC6 | PC7 | PC8 |
|---|---|---|---|---|---|---|---|---|
| N | 0.12 | **-0.65** | 0.27 | -0.11 | -0.04 | 0.10 | -0.02 | **0.68** |
| K | -0.30 | **-0.56** | 0.22 | -0.11 | -0.30 | 0.17 | 0.15 | **-0.63** |
| D | **-0.54** | 0.22 | -0.09 | 0.05 | -0.30 | 0.06 | **0.67** | 0.34 |
| C | -0.50 | -0.15 | 0.14 | -0.09 | **0.58** | **-0.60** | -0.02 | 0.00 |
| L | **0.55** | -0.04 | 0.18 | -0.08 | 0.32 | -0.11 | 0.72 | -0.16 |
| H | 0.05 | -0.09 | 0.26 | **0.93** | -0.11 | -0.21 | 0.01 | -0.02 |
| A | 0.11 | 0.31 | **0.60** | -0.31 | -0.48 | -0.44 | -0.10 | 0.02 |
| P | -0.19 | 0.29 | **0.62** | 0.04 | 0.37 | 0.59 | -0.06 | 0.01 |

**Table S8. Topological network parameters measured for each taxon, categorized based on the phylogenetic tree in Fig. 2.**

Aves are separated into adult and juveniles before categorized based on the phylogenetic position. Median scores (bold) and ranges (in parenthesis) of network parameters in Crurotarsi, Dinosauria (excluding modern birds), Saurischia (excluding modern birds), Theropoda (excluding modern birds), Aves, adult Aves, and juvenile Aves.

| | Number of Nodes (N) | Number of Links (K) | Density of Connections (D) | Mean Clustering | Mean Path | Heterogeneity of | Assortativity of | Parcellation (P) |
|---|---|---|---|---|---|---|---|---|

| | | | | Coefficient (C) | Length (L) | Connections (H) | Connections (A) | |
|---|---|---|---|---|---|---|---|---|
| Crurotarsi | **38 (28 - 46)** | **97 (56 - 114)** | **0.127 (0.095 - 0.159)** | **0.379 (0.266 - 0.45)** | **2.704 (2.429 - 3.278)** | **0.398 (0.284 - 0.471)** | **-0.138 (-0.217 - 0.185)** | **0.683 (0.422 - 0.799)** |
| *Riojasuchus* | 43 | 86 | 0.095 | 0.398 | 3.278 | 0.382 | -0.217 | 0.422 |
| *Aetosaurus* | 41 | 96 | 0.117 | 0.356 | 2.924 | 0.284 | 0.049 | 0.799 |
| *Desmatosuchus* | 38 | 89 | 0.127 | 0.379 | 2.704 | 0.471 | -0.215 | 0.738 |
| *Sphenosuchus* | 46 | 114 | 0.110 | 0.373 | 2.945 | 0.460 | 0.185 | 0.675 |
| *Dibothrosuchus* | 40 | 97 | 0.124 | 0.266 | 2.712 | 0.406 | -0.140 | 0.713 |
| *Dakosaurus* | 28 | 56 | 0.148 | 0.328 | 2.638 | 0.379 | -0.138 | 0.540 |
| *Alligator* (Adult) | 37 | 106 | 0.159 | 0.451 | 2.429 | 0.398 | -0.138 | 0.671 |
| *Alligator* (Juvenile) | 37 | 104 | 0.156 | 0.418 | 2.470 | 0.371 | -0.188 | 0.794 |
| *Crocodylus* | 37 | 103 | 0.155 | 0.415 | 2.441 | 0.418 | -0.084 | 0.683 |
| Dinosauria | **39 (32 - 44)** | **88 (61 - 112)** | **0.124(0.076 - 0.159)** | **0.332 (0.085 - 0.422)** | **2.778 (2.509 - 3.981)** | **0.355 (0.272 - 0.439)** | **-0.119 (-0.36 - 0.282)** | **0.602 (0.096 - 0.841)** |
| *Psittacosaurus* | 39 | 108 | 0.146 | 0.422 | 2.509 | 0.363 | -0.128 | 0.648 |
| Saurischia | **39 (32 - 44)** | **86 (61 - 112)** | **0.123 (0.076 - 0.159)** | **0.325 (0.085 - 0.419)** | **2.827 (2.53 - 3.981)** | **0.348 (0.272 - 0.439)** | **-0.109 (-0.36 - 0.282)** | **0.557 (0.096 - 0.841)** |
| *Eoraptor* | 38 | 90 | 0.128 | 0.325 | 2.727 | 0.332 | -0.109 | 0.405 |
| *Plateosaurus* | 37 | 66 | 0.099 | 0.085 | 3.128 | 0.371 | -0.360 | 0.096 |
| Theropoda | **39 (32-44)** | **86 (61-112)** | **0.123 (0.076 - 0.159)** | **0.338 (0.132 - 0.419)** | **2.827 (2.53 – 3.981)** | **0.348 (0.272 - 0.439)** | **-0.092 (-0.209 - 0.282)** | **0.687 (0.268 - 0.841)** |
| *Coelophysis* | 39 | 81 | 0.109 | 0.338 | 2.938 | 0.389 | -0.209 | 0.713 |

| | | | | | | | | |
|---|---|---|---|---|---|---|---|---|
| *Dilophosaurus* | 44 | 72 | 0.076 | 0.132 | 3.981 | 0.439 | 0.128 | 0.687 |
| *Compsognathus* | 32 | 61 | 0.123 | 0.198 | 3.012 | 0.329 | 0.282 | 0.841 |
| *Citipati* | 40 | 97 | 0.124 | 0.273 | 2.729 | 0.315 | -0.092 | 0.268 |
| *Velociraptor* | 39 | 86 | 0.116 | 0.394 | 2.827 | 0.348 | -0.138 | 0.557 |
| *Archaeopteryx* | 41 | 112 | 0.137 | 0.419 | 2.673 | 0.272 | -0.156 | 0.468 |
| *Ichthyornis* | 36 | 100 | 0.159 | 0.387 | 2.530 | 0.429 | 0.008 | 0.841 |
| Aves | **24.5 (6 - 34)** | **40 (9 - 72)** | **0.143 (0.128 - 0.6)** | **0.291 (0.126 - 0.733)** | **2.618 (1.4 - 2.877)** | **0.392 (0.296 - 0.634)** | **-0.192 (- 0.455 - 0.093)** | **0.462 (0 - 0.886)** |
| Adults | **10 (6-22)** | **15 (9 - 33)** | **0.333 (0.142 - 0.6)** | **0.157 (0.126 - 0.733)** | **1.844 (1.4 - 2.519)** | **0.416 (0.365 - 0.634)** | **-0.397 (- 0.454 - (- 0.222))** | **0 (0 - 0.478)** |
| *Nothura* | 6 | 9 | 0.600 | 0.733 | 1.400 | 0.365 | -0.455 | 0.000 |
| *Gallus* | 22 | 33 | 0.143 | 0.126 | 2.519 | 0.634 | -0.397 | 0.478 |
| *Geospiza* | 10 | 15 | 0.333 | 0.157 | 1.844 | 0.416 | -0.222 | 0.000 |
| Juvenile | **30 (27 - 34)** | **62 (47 - 64)** | **0.134 (0.128 - 0.143)** | **0.319 (0.263 - 0.348)** | **2.818 (2.717 - 2.877)** | **0.368 (0.296 - 0.48)** | **-0.039 (- 0.161 - 0.093)** | **0.604 (0.446 - 0.886)** |
| *Nothura* | 30 | 62 | 0.143 | 0.348 | 2.818 | 0.296 | -0.039 | 0.604 |
| *Gallus* | 34 | 72 | 0.128 | 0.263 | 2.717 | 0.480 | -0.161 | 0.446 |
| *Geospiza* | 27 | 47 | 0.134 | 0.319 | 2.877 | 0.368 | 0.093 | 0.886 |

**Table S9. Network parameters categorized by diet.** Median and ranges (in parenthesis) of network parameters of carnivores, herbivores, omnivores, omnivores when Aves are excluded, and piscivores. Herbivores include *Psittacosaurus* and *Plateosaurus*. Piscivore is represented by *Ichthyornis dispar*. Aves, *Aetosaurus*, *Eoraptor*, and *Desmatosuchus* are omnivores.

| | Number of Nodes (N) | Number of Links (K) | Density of Connections (D) | Mean Clustering Coefficient (C) | Mean Path Length (L) | Heterogeneity of Connections (H) | Assortativity of Connections (A) | Parcellation (P) |
|---|---|---|---|---|---|---|---|---|
| Carnivore (n=10) | 39 (28-46) | 85 (56-114) | 0.124 (0.076-0.159) | 0.373 (-0.132-0.451) | 2.729 (2.429-4.981) | 0.382 (0.272-0.46) | -0.138 (-0.217-0.282) | 0.683 (0.268-0.841) |
| Herbivore (n=2) | 37- 39 | 66- 108 | 0.099- 0.146 | 0.085-0.422 | 2.509-3.128 | 0.363- 0.371 | -0.36-(-0.128) | 0.096-0.648 |
| Omnivore (n=9) | 30(6-41) | 62 (9-96) | 0.134 (0.117-0.6) | 0.325 (0.126-0.733) | 2.717 (1.4-2.924) | 0.368 (0.284-0.634) | -0.161 (-0.455-0.093) | 0.478 (0-0.886) |
| exclude Aves (n=3) | 38 (38-41) | 90 (89 – 96) | 0.127(0.117-0.128) | 0.356 (0.325-0.379) | 2.727 (2.704-2.924) | 0.332 (0.284-0.471) | -0.109 (-0.215 -0.049) | 0.738 (0.405-0.799) |
| Piscivore (n=1) | 36 | 100 | 0.159 | 0.387 | 2.530 | 0.429 | 0.008 | 0.841 |

**Table S12. The number of modules generated from each network using Node-based Informed Modularity Strategy (NIMS).** The numbers in bold are the medians and ranges of modules calculated from each class. Juvenile Aves have a median of four modules, which is between the median number of modules in adult Aves and Crurotarsi. Dinosauria have the highest median number of modules.

| | Number of Modules |
|---|---|
| Crurotarsi | **4 (3-6)** |
| *Riojasuchus* | 6 |
| *Aetosaurus* | 6 |
| *Desmatosuchus* | 4 |
| *Sphenosuchus* | 5 |
| *Dibothrosuchus* | 6 |
| *Dakosaurus* | 4 |
| *Alligator* (Adult) | 4 |
| *Alligator* (Juvenile) | 3 |
| *Crocodylus* | 4 |
| Dinosauria | **4.5 (4-8)** |
| *Psittacosaurus* | 6 |
| Saurischia | **4 (4-8)** |
| *Eoraptor* | 5 |
| *Plateosaurus* | 6 |
| Theropoda | **4 (4-8)** |
| *Coelophysis* | 6 |
| *Dilophosaurus* | 4 |
| *Compsognathus* | 4 |

| | |
|---|---|
| *Citipati* | 8 |
| *Velociraptor* | 4 |
| *Archaeopteryx* | 4 |
| *Ichthyornis* | 4 |
| Aves | **4 (1-5)** |
| Adult | **1 (1-4)** |
| *Nothura* | 1 |
| *Gallus* | 4 |
| *Geospiza* | 1 |
| Juvenile | **4 (4-5)** |
| *Nothura* | 4 |
| *Gallus* | 5 |
| *Geospiza* | 4 |

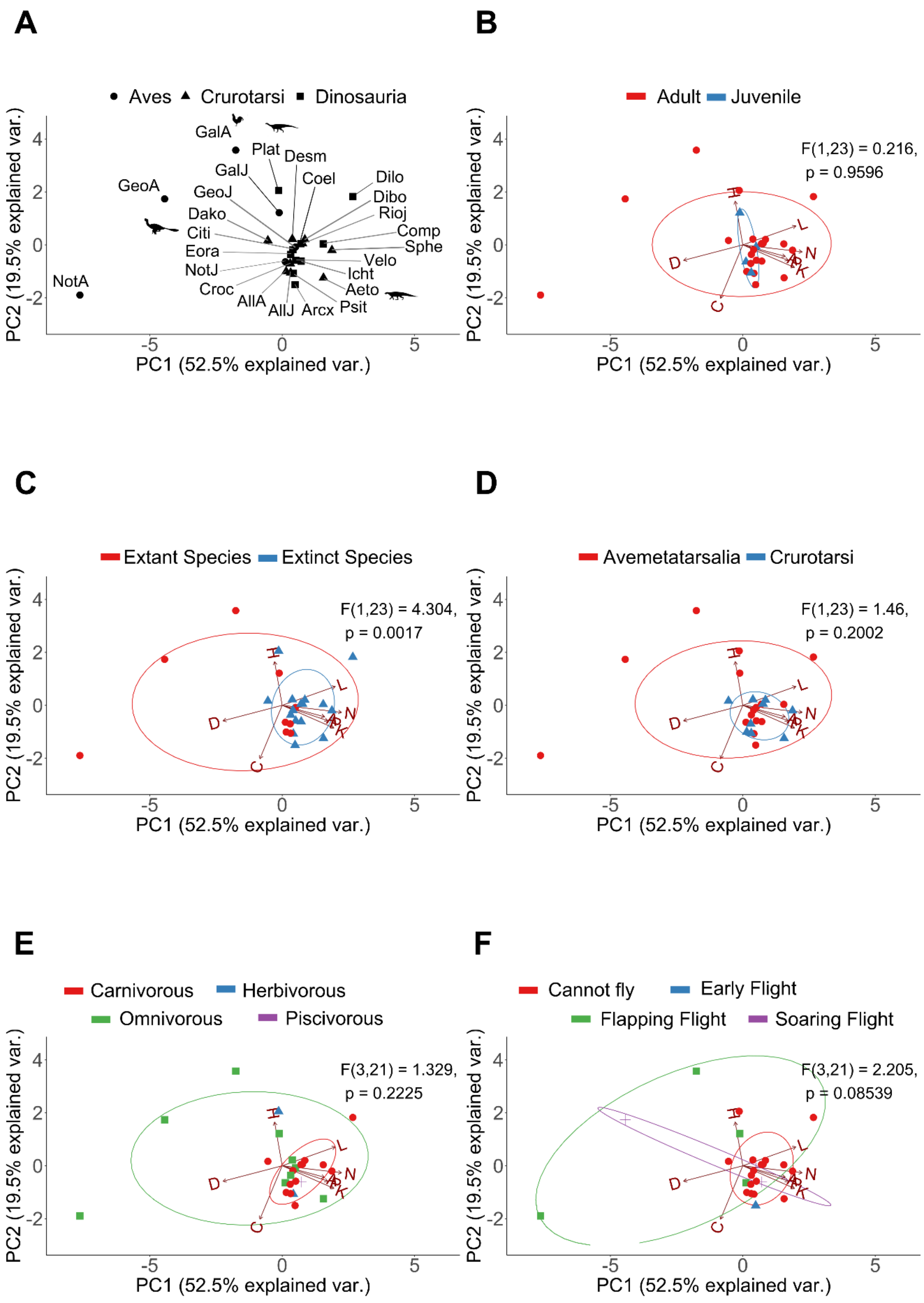
A
B
C
D
E
F
Aves
Crurotarsi
Dinosauria
GalA
Plat
Desm
GalJ
Coel
Dilo
GeoA
GeoJ
Dibo
Rioj
Dako
Comp
Citi
Sphe
Eora
Velo
NotJ
Icht
NotA
Croc
Aeto
AllA
AllJ
Arcx
Psit
Adult
Juvenile
F(1,23) = 0.216,
p = 0.9596
Extant Species
Extinct Species
F(1,23) = 4.304,
p = 0.0017
Avemetatarsalia
Crurotarsi
F(1,23) = 1.46,
p = 0.2002
Carnivorous
Herbivorous
Omnivorous
Piscivorous
F(3,21) = 1.329,
p = 0.2225
Cannot fly
Early Flight
Flapping Flight
Soaring Flight
F(3,21) = 2.205,
p = 0.08539
H
L
N
K
D
C
PC1 (52.5% explained var.)
PC2 (19.5% explained var.)

**Figure S1. First two PC of topological parameters for all taxa**. (A) Skull distribution for each taxon (see labels below). (B) Comparison of juveniles versus adults shows that juveniles lie in the morphospace of adults. (C) Comparison of extant taxa versus extinct taxa shows that extant taxa occupy a different morphospace from extinct taxa. (D) Comparison of Crurotasi versus Avemetatarsalia shows that Crurotarsi overlap with the Avemetatarsalia morphospace. (E) Taxa with different diets overlap with each other. (F) Taxa with different ability to fly overlap with each other. Ellipses show a normal distribution confidence interval around groups for comparison. Labels: N, number of nodes; K: number of links; D, density of connection; C: mean clustering coefficient; H: heterogeneity of connection; L: mean shortest path length; A: assortativity; P, parcellation. Aeto, *Aetosaurus;* AllA, adult *Alligator*; AllJ, juvenile *Alligator*; Arcx, *Archaeopteryx*; Citi, *Citipati*; Coel, *Coelophysis*; Comp, *Compsognathus*; Croc, *Crocodylus*; Dako, *Dakosaurus*; Desm, *Desmatosuchus*; Dibo, *Dibothrosuchus*; Dilo, *Dilophosaurus*; Eora, *Eoraptor*; GalA, adult *Gallus*; GalJ, juvenile *Gallus*; GeoA, adult *Geospiza*; GeoJ, juvenile *Geospiza*; Icht, *Ichthyornis*; NotA, adult *Nothura*; NotJ, juvenile *Nothura*; Plat, *Plateosaurus*; Psit, *Psittacosaurus*; Rioj, *Riojasuchus*; Sphe, *Sphenosuchus*; Velo, *Velociraptor.*

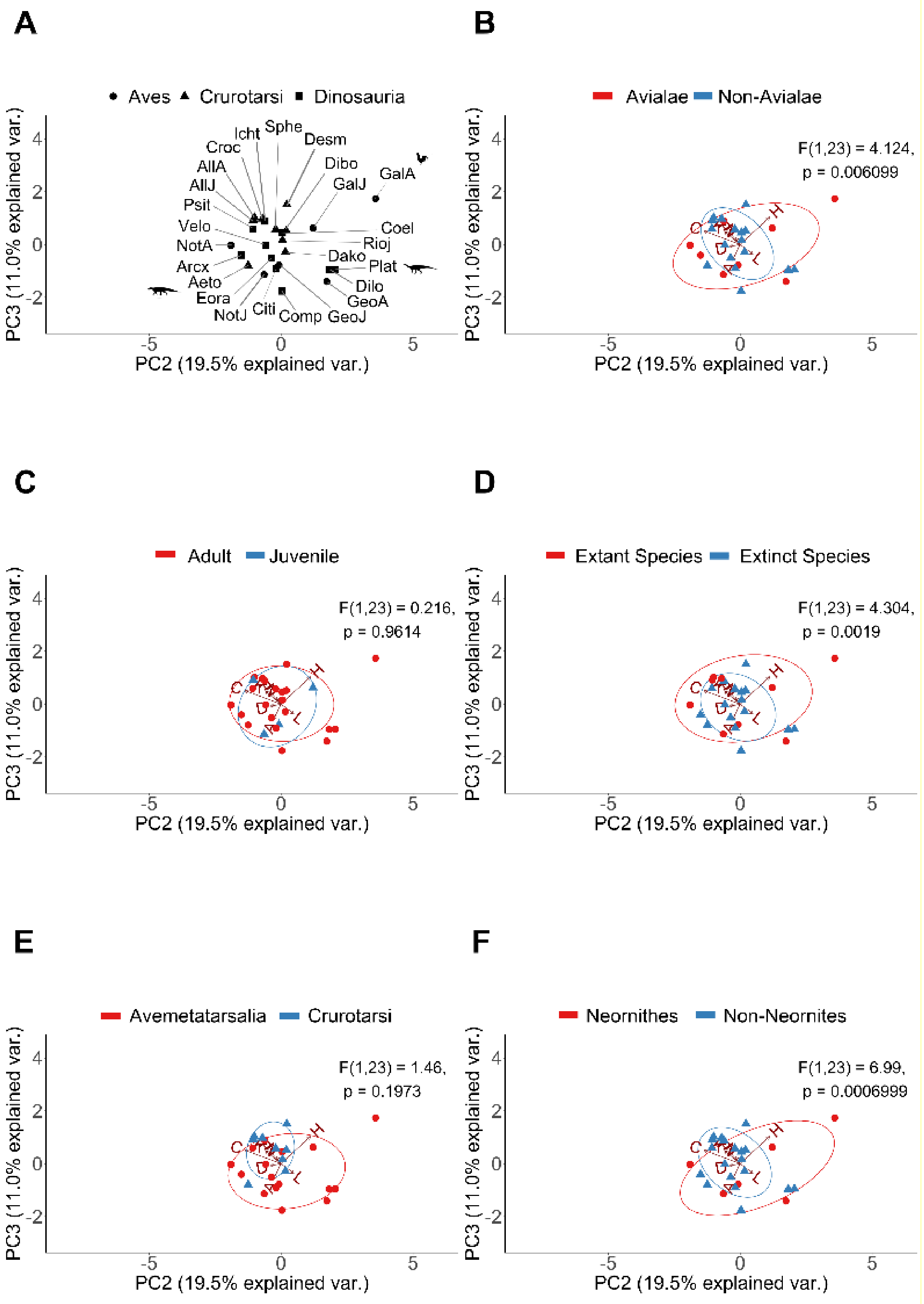
A
Aves Crurotarsi Dinosauria
Icht Sphe Desm Croc AllA Dibo AllJ GalJ GalA Psit Velo Coel Rioj NotA Dako Arcx Plat Aeto Dilo Eora GeoA NotJ Citi Comp GeoJ
PC3 (11.0% explained var.)
PC2 (19.5% explained var.)
B
Avialae Non-Avialae
F(1,23) = 4.124,
p = 0.006099
C
Adult Juvenile
F(1,23) = 0.216,
p = 0.9614
D
Extant Species Extinct Species
F(1,23) = 4.304,
p = 0.0019
E
Avemetatarsalia Crurotarsi
F(1,23) = 1.46,
p = 0.1973
F
Neornithes Non-Neornites
F(1,23) = 6.99,
p = 0.0006999

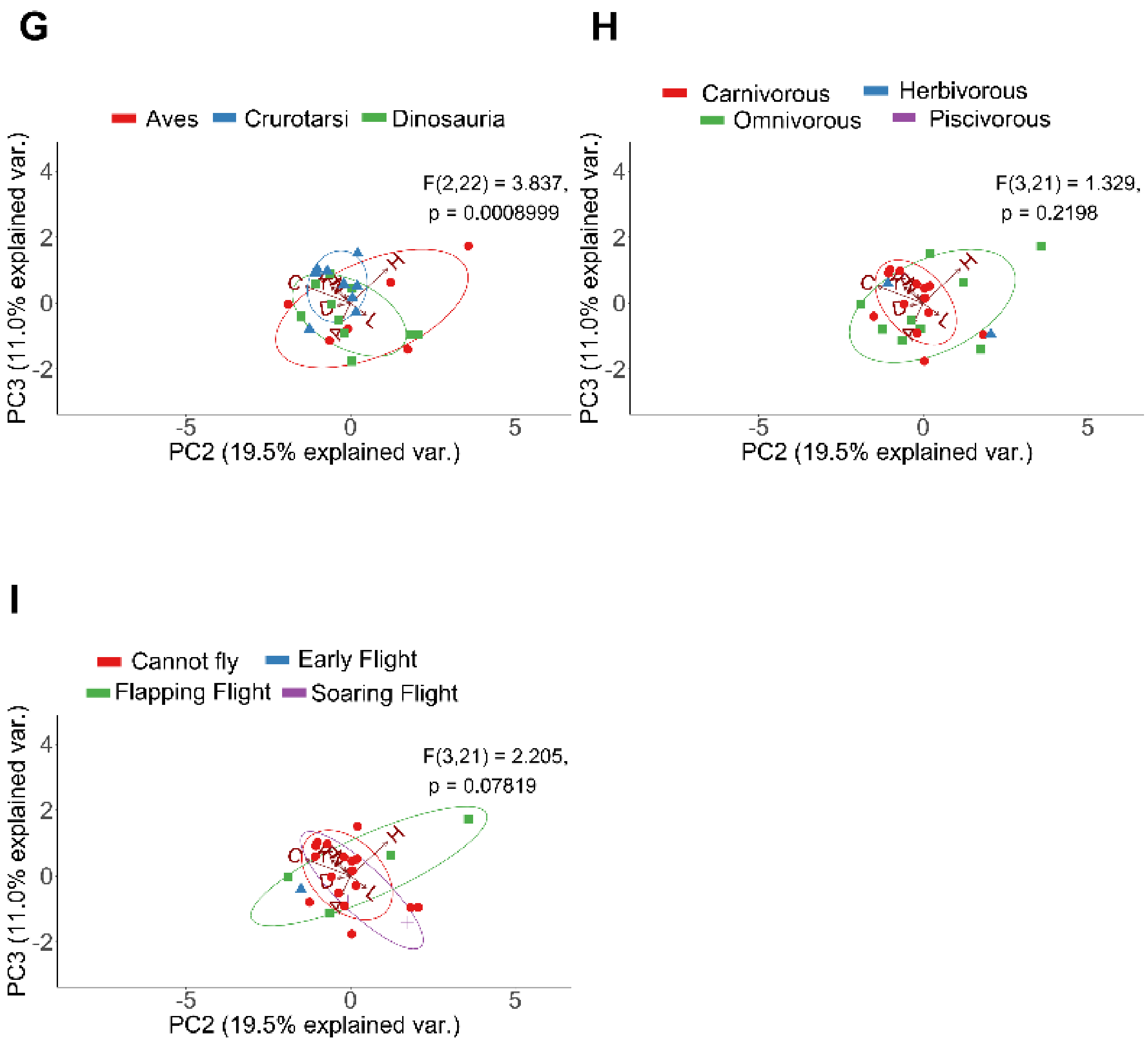

**Figure S2. Second and third PC of topological parameters for all taxa**. (A) Skull distribution for each taxon (see labels below). (B) Comparison of Avialae versus non-Avialae shows that Avialae occupy a different morphospace from non-Avialae. (C) Comparison of juveniles versus adults shows that juvenile overlap with the morphospace of adults. (D) Comparison of extant taxa versus extinct taxa shows that extant taxa occupy a different morphospace from extinct taxa. (E) Comparison of Crurotasi versus Avemetatarsalia shows that Crurotarsi overlap with the Avemetatarsalia morphospace. (F) Neornithes occupy a different morphospace from non-Neornithes. (G) Aves, Dinosauria, and Crurotarsi occupy a different morphospace from each other. (H) Taxa with different dietary requirements overlap with each other. (I) Taxa with different ability to fly overlap with each other. Ellipses show a normal distribution confidence interval around groups for comparison. Labels: N, number of nodes; K: number of links; D, density of connection; C: mean clustering coefficient; H: heterogeneity of connection; L: mean shortest path length; A: assortativity; P, parcellation. Aeto, *Aetosaurus;* AllA, adult *Alligator*; AllJ, juvenile *Alligator*; Arcx, *Archaeopteryx*; Citi, *Citipati*; Coel, *Coelophysis*; Comp, *Compsognathus*; Croc, *Crocodylus*; Dako, *Dakosaurus*; Desm, *Desmatosuchus*; Dibo, *Dibothrosuchus*; Dilo, *Dilophosaurus*; Eora, *Eoraptor*; GalA, adult *Gallus*; GalJ, juvenile *Gallus*; GeoA, adult *Geospiza*; GeoJ, juvenile *Geospiza*; Icht, *Ichthyornis*; NotA, adult *Nothura*; NotJ, juvenile *Nothura*; Plat, *Plateosaurus*; Psit, *Psittacosaurus*; Rioj, *Riojasuchus*; Sphe, *Sphenosuchus*; Velo, *Velociraptor.*

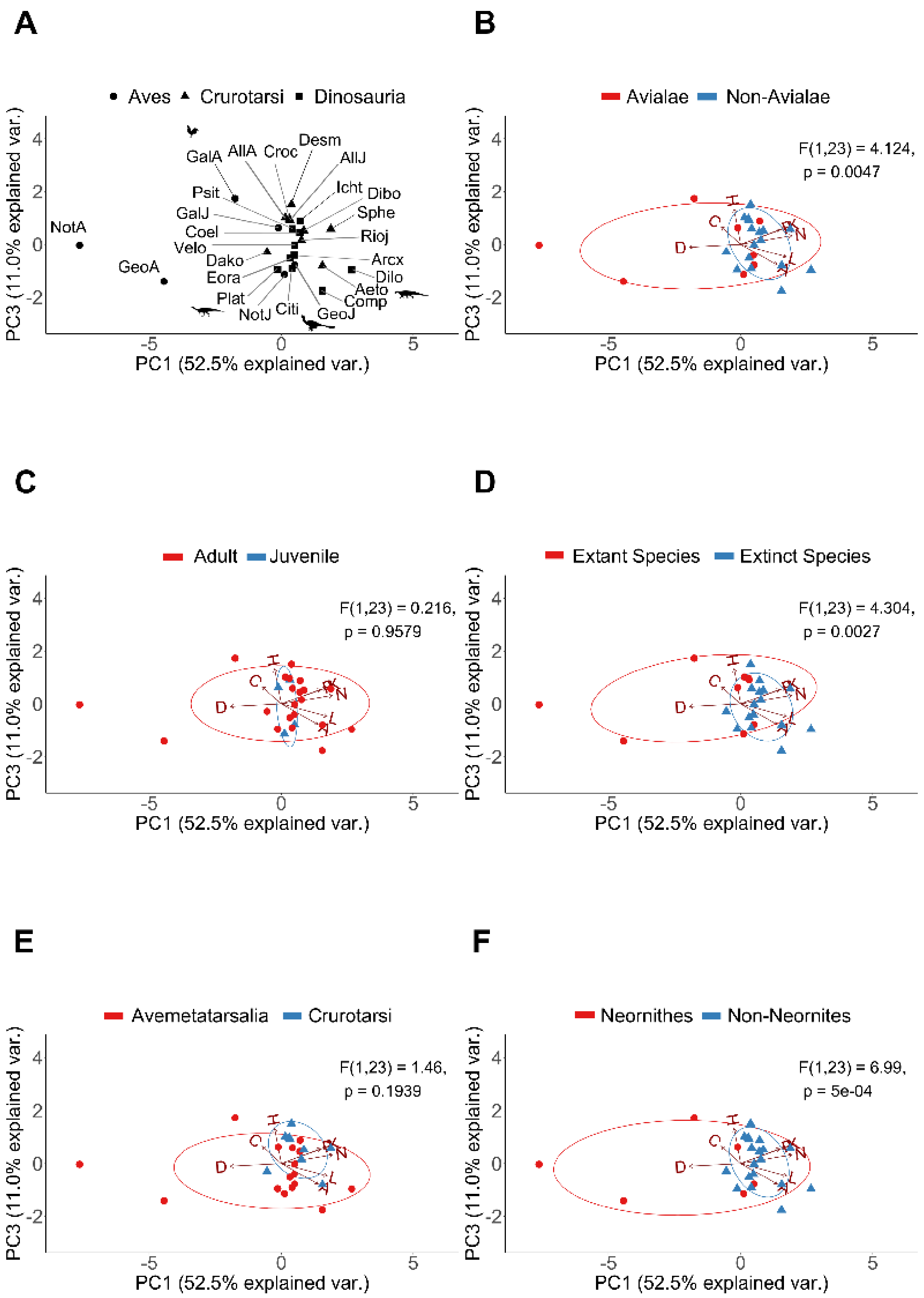
A
Aves Crurotarsi Dinosauria
PC3 (11.0% explained var.)
PC1 (52.5% explained var.)
NotA GeoA GalA AllA Croc Desm AllJ Psit Icht Dibo GalJ Sphe Coel Rioj Velo Dako Arcx Eora Dilo Aeto Plat Comp NotJ Citi GeoJ
B
Avialae Non-Avialae
F(1,23) = 4.124,
p = 0.0047
C
Adult Juvenile
F(1,23) = 0.216,
p = 0.9579
D
Extant Species Extinct Species
F(1,23) = 4.304,
p = 0.0027
E
Avemetatarsalia Crurotarsi
F(1,23) = 1.46,
p = 0.1939
F
Neornithes Non-Neornites
F(1,23) = 6.99,
p = 5e-04

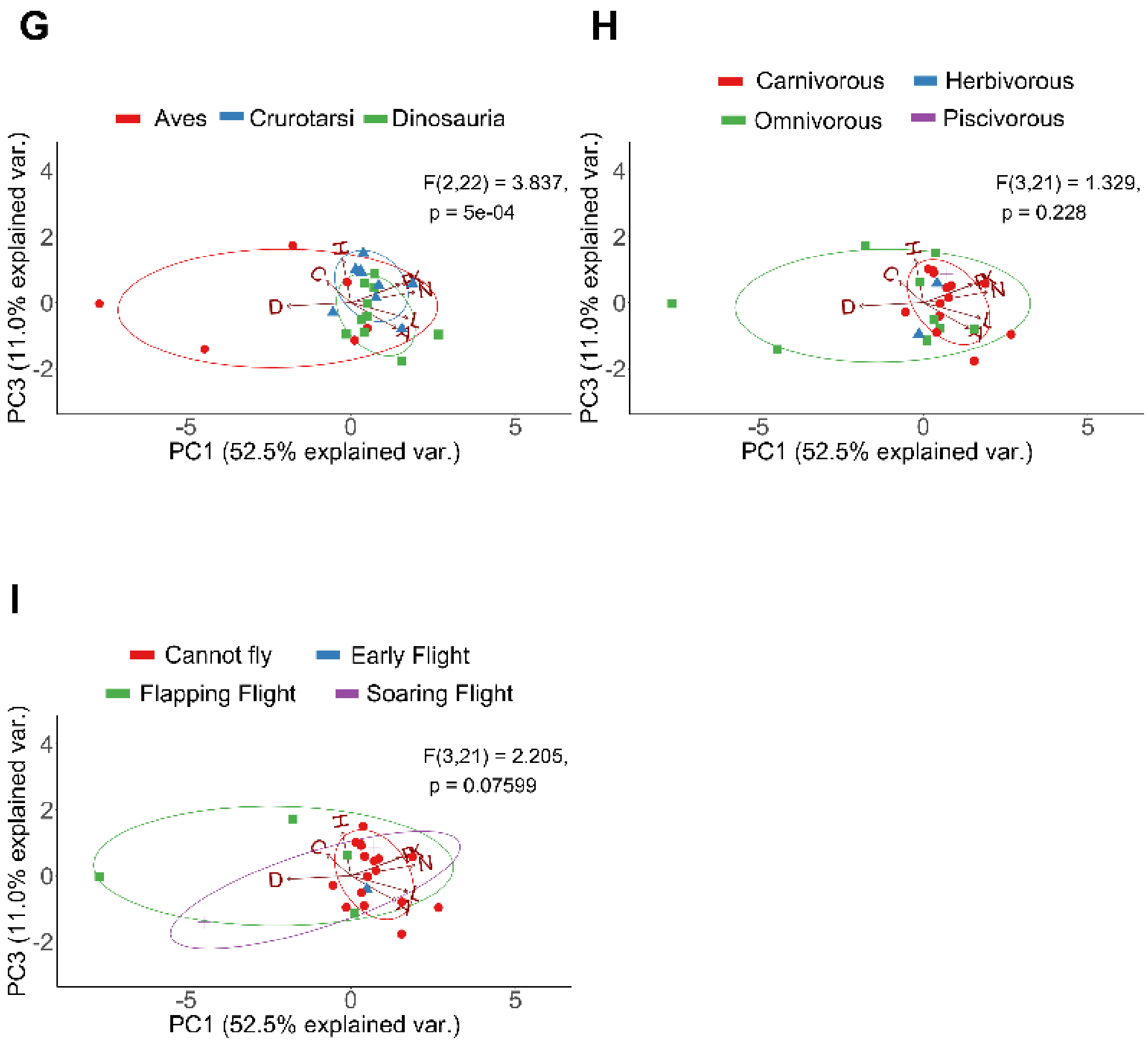

**Figure S3. First and third PC of topological parameters for all taxa**. (A) Skull distribution for each taxon (see labels below). (B) Comparison of Avialae versus non-Avialae shows that Avialae occupy a different morphospace from non-Avialae. (C) Comparison of juveniles versus adults shows that juvenile overlap with the morphospace of adults. (D) Comparison of extant taxa versus extinct taxa shows that extant taxa occupy a different morphospace from extinct taxa. (E) Comparison of Crurotasi versus Avemetatarsalia shows that Crurotarsi overlap with the Avemetatarsalia morphospace. (F) Neornithes occupy a different morphospace from non-Neornithes. (G) Aves, Dinosauria, and Crurotarsi occupy a different morphospace from each other. (H) Taxa with different dietary requirements overlap with each other. (I) Taxa with different ability to fly overlap with each other. Ellipses show a normal distribution confidence interval around groups for comparison. Labels: N, number of nodes; K: number of links; D, density of connection; C: mean clustering coefficient; H: heterogeneity of connection; L: mean shortest path length; A: assortativity; P, parcellation. Aeto, *Aetosaurus;* AllA, adult *Alligator*; AllJ, juvenile *Alligator*; Arcx, *Archaeopteryx*; Citi, *Citipati*; Coel, *Coelophysis*; Comp, *Compsognathus*; Croc, *Crocodylus*; Dako, *Dakosaurus*; Desm, *Desmatosuchus*;

Dibo, *Dibothrosuchus*; Dilo, *Dilophosaurus*; Eora, *Eoraptor*; GalA, adult *Gallus*; GalJ, juvenile *Gallus*; GeoA, adult *Geospiza*; GeoJ, juvenile *Geospiza*; Icht, *Ichthyornis*; NotA, adult *Nothura*; NotJ, juvenile *Nothura*; Plat, *Plateosaurus*; Psit, *Psittacosaurus*; Rioj, *Riojasuchus*; Sphe, *Sphenosuchus*; Velo, *Velociraptor*.

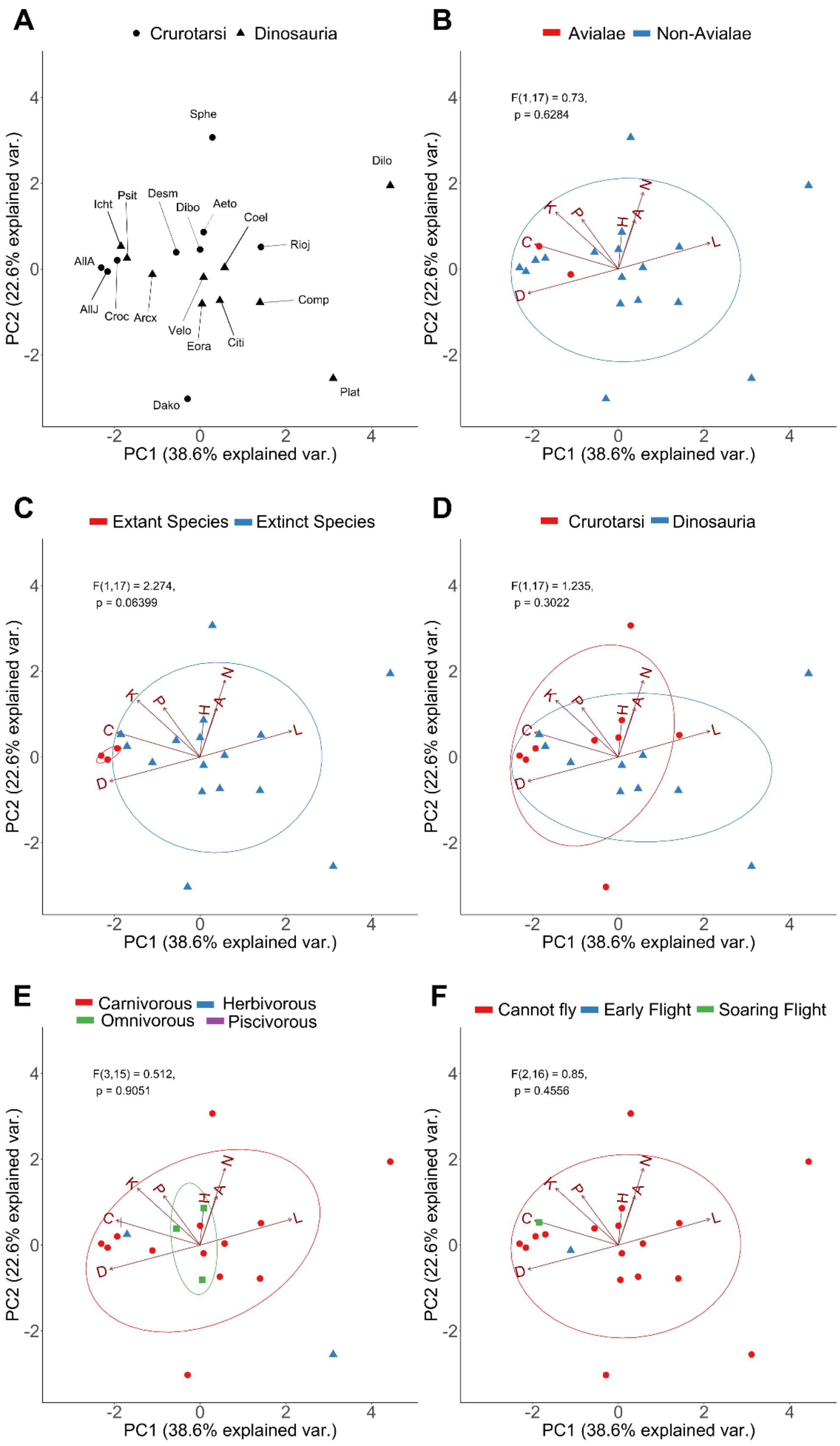
A
Crurotarsi
Dinosauria
Sphe
Dilo
Icht
Psit
Desm
Dibo
Aeto
Coel
Rioj
AllA
AllJ
Croc
Arcx
Velo
Eora
Citi
Comp
Plat
Dako
B
Avialae
Non-Avialae
F(1,17) = 0.73,
p = 0.6284
C
Extant Species
Extinct Species
F(1,17) = 2.274,
p = 0.06399
D
Crurotarsi
Dinosauria
F(1,17) = 1.235,
p = 0.3022
E
Carnivorous
Herbivorous
Omnivorous
Piscivorous
F(3,15) = 0.512,
p = 0.9051
F
Cannot fly
Early Flight
Soaring Flight
F(2,16) = 0.85,
p = 0.4556
PC1 (38.6% explained var.)
PC2 (22.6% explained var.)
K
P
H
A
N
C
L
D

**Figure S4. First two PC of topological parameters for all taxa excluding avians**. (A) Skull distribution for each taxon (see labels below). (B) Avialae lies in the morphospace of non-Avialae taxa. (C) Extant crurotarsans overlap with extinct taxa. (D) Crurotarsi overlap with Dinosauria. (E) Comparison of dietary requirements shows herbivore, omnivore, and piscivore overlap with carnivores. (F) Archosaurs with either soaring flight or early flight overlap with non-flying taxa. Ellipses show a normal distribution confidence interval around groups for comparison. Labels: N, number of nodes; K: number of links; D, density of connection; C: mean clustering coefficient; H: heterogeneity of connection; L: mean shortest path length; A: assortativity; P, parcellation. Aeto, *Aetosaurus;* AllA, adult *Alligator*; AllJ, juvenile *Alligator*; Arcx, *Archaeopteryx*; Citi, *Citipati*; Coel, *Coelophysis*; Comp, *Compsognathus*; Croc, *Crocodylus*; Dako, *Dakosaurus*; Desm, *Desmatosuchus*; Dibo, *Dibothrosuchus*; Dilo, *Dilophosaurus*; Eora, *Eoraptor*; Icht, *Ichthyornis*; Plat, *Plateosaurus*; Psit, *Psittacosaurus*; Rioj, *Riojasuchus*; Sphe, *Sphenosuchus*; Velo, *Velociraptor*.

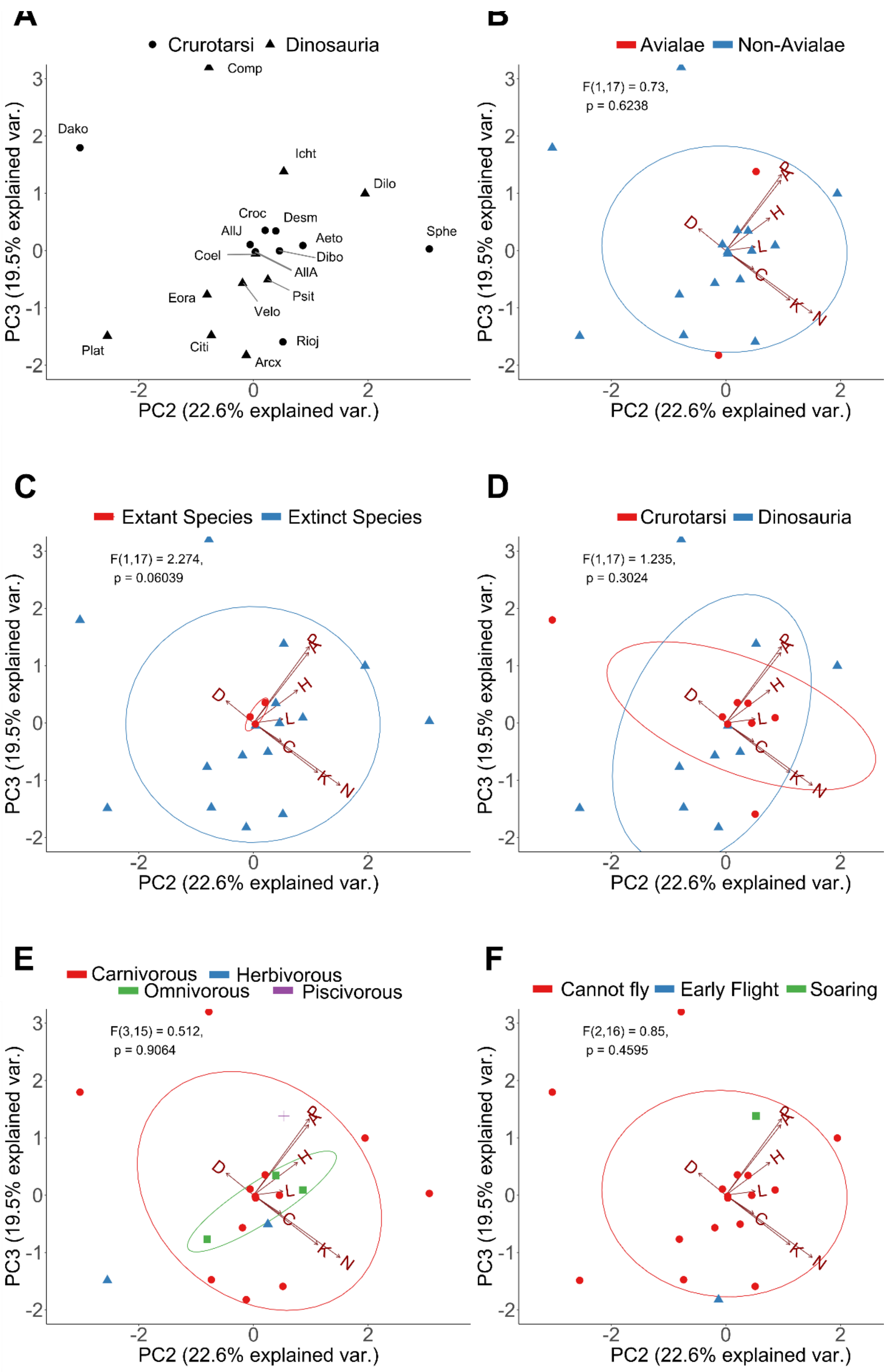
A
Crurotarsi
Dinosauria
Comp
Dako
Icht
Dilo
Croc
Desm
AllJ
Aeto
Sphe
Coel
Dibo
AllA
Psit
Eora
Velo
Rioj
Plat
Citi
Arcx
PC3 (19.5% explained var.)
PC2 (22.6% explained var.)
B
Avialae
Non-Avialae
F(1,17) = 0.73,
p = 0.6238
C
Extant Species
Extinct Species
F(1,17) = 2.274,
p = 0.06039
D
Crurotarsi
Dinosauria
F(1,17) = 1.235,
p = 0.3024
E
Carnivorous
Herbivorous
Omnivorous
Piscivorous
F(3,15) = 0.512,
p = 0.9064
F
Cannot fly
Early Flight
Soaring
F(2,16) = 0.85,
p = 0.4595
R
H
D
L
C
K
N

**Figure S5. Second and third PC of topological parameters for all taxa excluding avians**. (A) Skull distribution for each taxon (see labels below). (B) Avialae lies in the morphospace of non-Avialae taxa. (C) Extant crurotarsans overlap with extinct taxa. (D) Crurotarsi overlap with Dinosauria. (E) Herbivore and piscivore overlap with carnivores but not omnivore. (F) Taxon with soaring flight overlap with non-flying taxa but not the taxon with early flight. Ellipses show a normal distribution confidence interval around groups for comparison. Labels: N, number of nodes; K: number of links; D, density of connection; C: mean clustering coefficient; H: heterogeneity of connection; L: mean shortest path length; A: assortativity; P, parcellation. Aeto, *Aetosaurus;* AllA, adult *Alligator*; AllJ, juvenile *Alligator*; Arcx, *Archaeopteryx*; Citi, *Citipati*; Coel, *Coelophysis*; Comp, *Compsognathus*; Croc, *Crocodylus*; Dako, *Dakosaurus*; Desm, *Desmatosuchus*; Dibo, *Dibothrosuchus*; Dilo, *Dilophosaurus*; Eora, *Eoraptor*; Icht, *Ichthyornis*; Plat, *Plateosaurus*; Psit, *Psittacosaurus*; Rioj, *Riojasuchus*; Sphe, *Sphenosuchus*; Velo, *Velociraptor*.

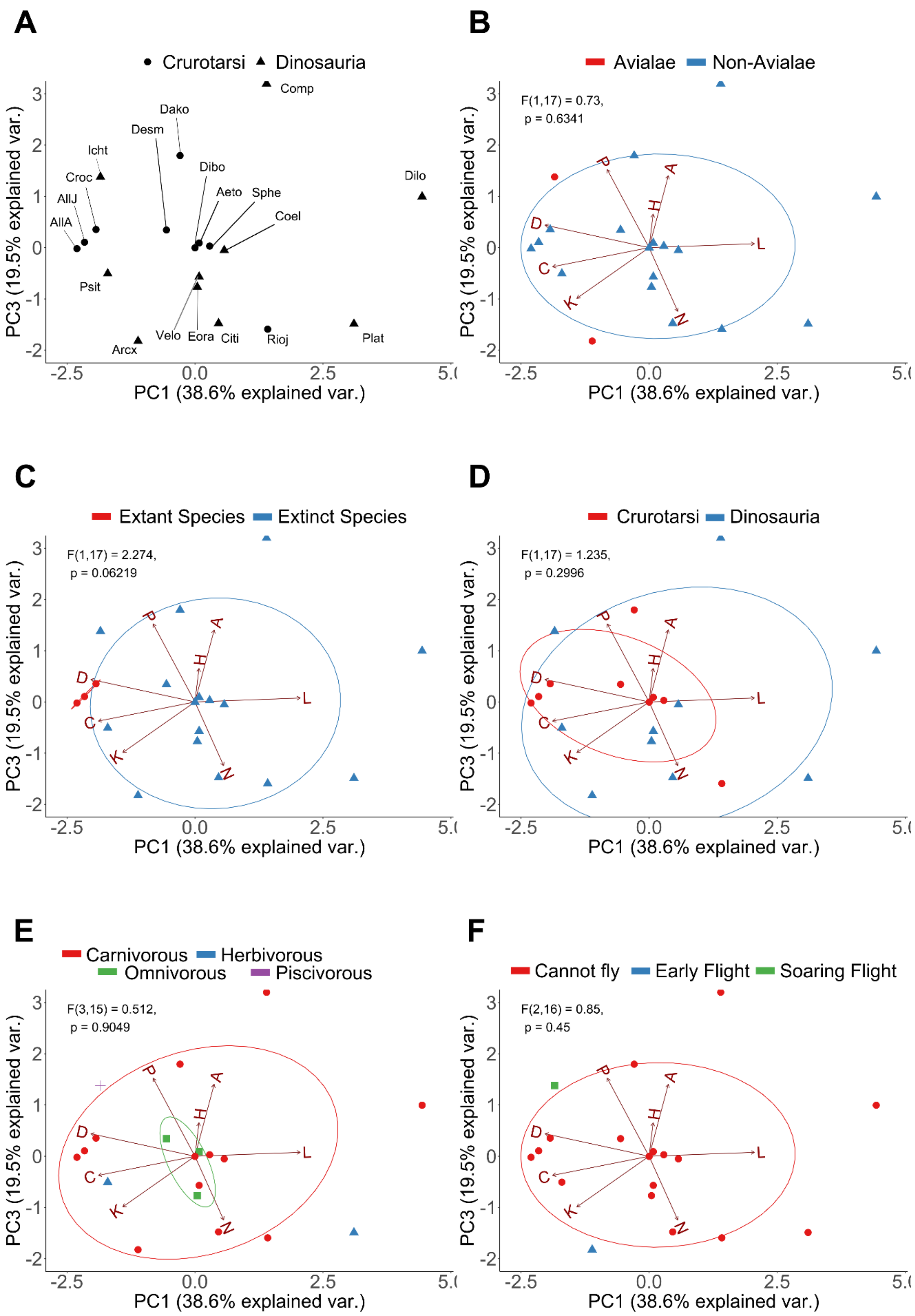
A
Crurotarsi Dinosauria
Comp
Dako
Desm
Icht
Croc
AllJ
AllA
Dibo
Aeto
Sphe
Coel
Dilo
Psit
Velo
Eora
Citi
Rioj
Plat
Arcx
B
Avialae Non-Avialae
F(1,17) = 0.73,
p = 0.6341
C
Extant Species Extinct Species
F(1,17) = 2.274,
p = 0.06219
D
Crurotarsi Dinosauria
F(1,17) = 1.235,
p = 0.2996
E
Carnivorous Herbivorous
Omnivorous Piscivorous
F(3,15) = 0.512,
p = 0.9049
F
Cannot fly Early Flight Soaring Flight
F(2,16) = 0.85,
p = 0.45
PC1 (38.6% explained var.)
PC3 (19.5% explained var.)

**Figure S6. First and third PC of topological parameters for all taxa excluding avians**. (A) Skull distribution for each taxon (see labels below). (B) Avialae lies in the morphospace of non-Avialae taxa. (C) Extant crurotarsans overlap extinct taxa. (D) Crurotarsi overlap with Dinosauria. (E) Herbivore and piscivore overlap with carnivores but not omnivore. (F) Taxon with soaring flight overlap with non-flying taxa but not the taxon with early flight. Ellipses show a normal distribution confidence interval around groups for comparison. Labels: N, number of nodes; K: number of links; D, density of connection; C: mean clustering coefficient; H: heterogeneity of connection; L: mean shortest path length; A: assortativity; P, parcellation. Aeto, *Aetosaurus;* AllA, adult *Alligator*; AllJ, juvenile *Alligator*; Arcx, *Archaeopteryx*; Citi, *Citipati*; Coel, *Coelophysis*; Comp, *Compsognathus*; Croc, *Crocodylus*; Dako, *Dakosaurus*; Desm, *Desmatosuchus*; Dibo, *Dibothrosuchus*; Dilo, *Dilophosaurus*; Eora, *Eoraptor*; Icht, *Ichthyornis*; Plat, *Plateosaurus*; Psit, *Psittacosaurus*; Rioj, *Riojasuchus*; Sphe, *Sphenosuchus*; Velo, *Velociraptor*

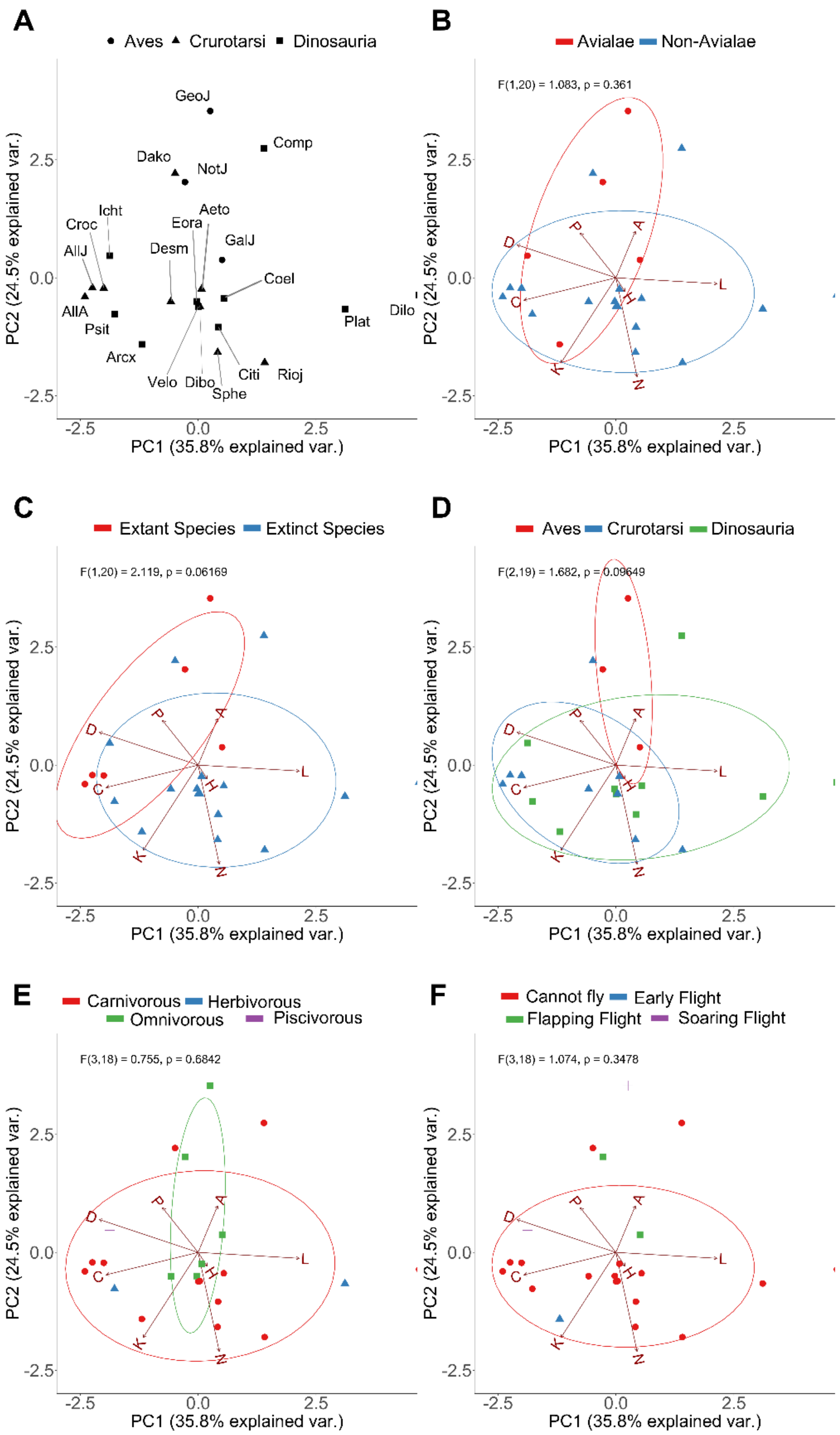
A
Aves
Crurotarsi
Dinosauria
GeoJ
Comp
Dako
NotJ
Icht
Croc
AllJ
Aeto
Eora
GalJ
Desm
Coel
Dilo
AllA
Psit
Plat
Arcx
Velo
Dibo
Citi
Rioj
Sphe
PC2 (24.5% explained var.)
PC1 (35.8% explained var.)
B
Avialae
Non-Avialae
F(1,20) = 1.083, p = 0.361
C
Extant Species
Extinct Species
F(1,20) = 2.119, p = 0.06169
D
Aves
Crurotarsi
Dinosauria
F(2,19) = 1.682, p = 0.09649
E
Carnivorous
Herbivorous
Omnivorous
Piscivorous
F(3,18) = 0.755, p = 0.6842
F
Cannot fly
Early Flight
Flapping Flight
Soaring Flight
F(3,18) = 1.074, p = 0.3478

**Figure S7. First two PC of topological parameters for all taxa excluding adult avians**. (A) Skull distribution for each taxon (see labels below). (B) PCA shows an overlap in morphospace occupation between Avialae and non-Avialae species. (C) Extant taxa overlap with extinct taxa. (D) Juvenile avians overlap with crurotarsans and non-avian dinosaurs. (E) Taxa with different dietary requirement overlap with each other. (F) Taxa with different abilities to flight overlap with each other. Ellipses show a normal distribution confidence interval around groups for comparison. Labels: N, number of nodes; K: number of links; D, density of connection; C: mean clustering coefficient; H: heterogeneity of connection; L: mean shortest path length; A: assortativity; P, parcellation. Aeto, *Aetosaurus;* AllA, adult *Alligator*; AllJ, juvenile *Alligator*; Arcx, *Archaeopteryx*; Citi, *Citipati*; Coel, *Coelophysis*; Comp, *Compsognathus*; Croc, *Crocodylus*; Dako, *Dakosaurus*; Desm, *Desmatosuchus*; Dibo, *Dibothrosuchus*; Dilo, *Dilophosaurus*; Eora, *Eoraptor*; GalJ, juvenile *Gallus*; GeoJ, juvenile *Geospiza*; Icht, *Ichthyornis*; NotJ, juvenile *Nothura*; Plat, *Plateosaurus*; Psit, *Psittacosaurus*; Rioj, *Riojasuchus*; Sphe, *Sphenosuchus*; Velo, *Velociraptor*.

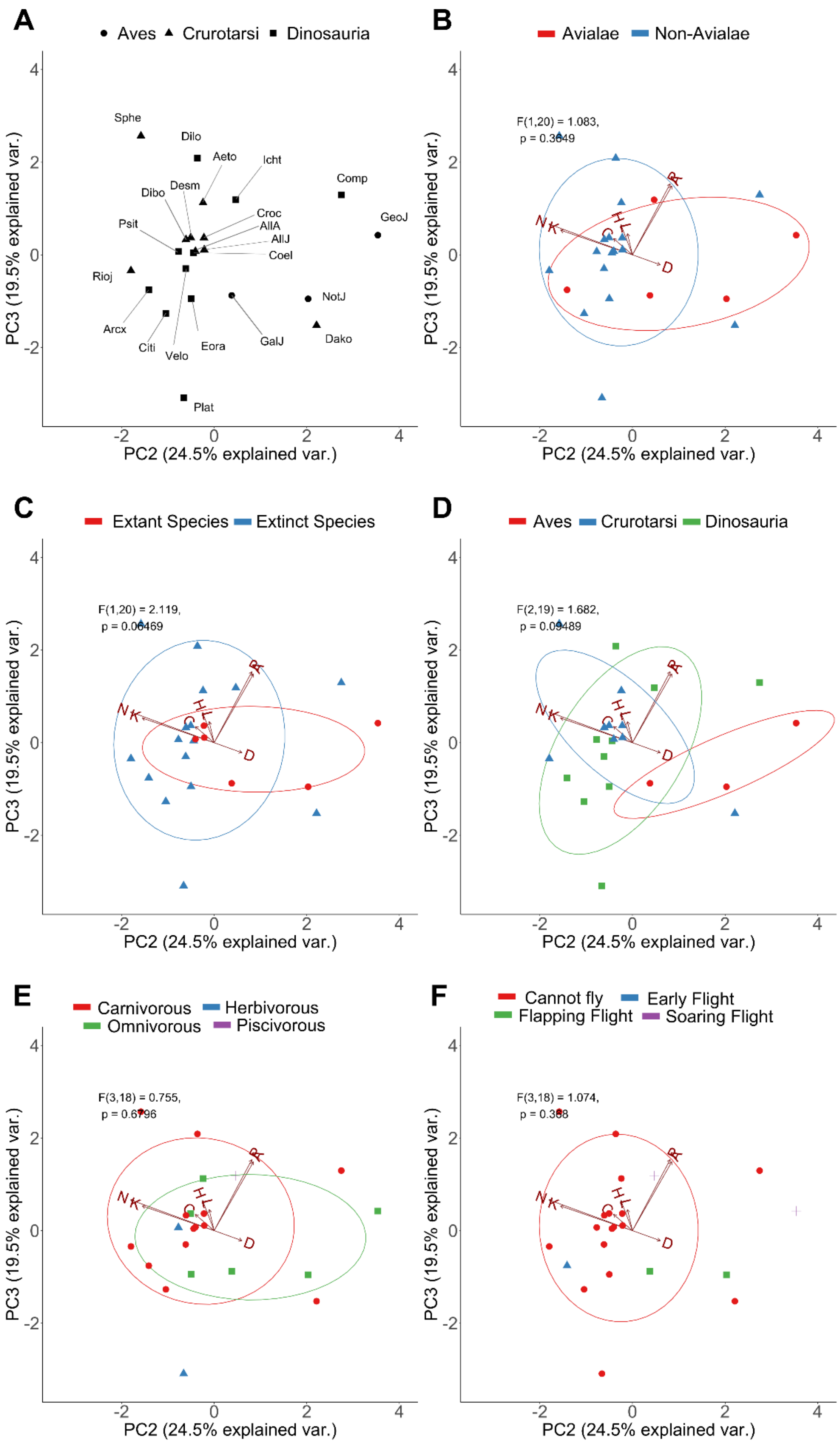
A
Aves Crurotarsi Dinosauria
Sphe
Dilo
Aeto
Icht
Comp
Desm
Dibo
Croc
AllA
AllJ
GeoJ
Psit
Coel
Rioj
NotJ
Arcx
GalJ
Dako
Citi
Velo
Eora
Plat
PC3 (19.5% explained var.)
PC2 (24.5% explained var.)
B
Avialae Non-Avialae
F(1,20) = 1.083,
p = 0.3049
C
Extant Species Extinct Species
F(1,20) = 2.119,
p = 0.06469
D
Aves Crurotarsi Dinosauria
F(2,19) = 1.682,
p = 0.09489
E
Carnivorous Herbivorous
Omnivorous Piscivorous
F(3,18) = 0.755,
p = 0.6796
F
Cannot fly Early Flight
Flapping Flight Soaring Flight
F(3,18) = 1.074,
p = 0.368
N
K
C
H
L
R
D

**Figure S8. Second and third PC of topological parameters for all taxa excluding adult avians.** (A) Skull distribution for each taxon (see labels below). (B) PCA shows an overlap in morphospace occupation between Avialae and non-Avialae species. (C) Extant taxa overlap with extinct taxa. (D) Juvenile avians overlap with crurotarsans and non-avian dinosaurs. (E) Taxa with different dietary requirement overlap with each other. (F) Taxa with different abilities to flight overlap with each other. Ellipses show a normal distribution confidence interval around groups for comparison. Labels: N, number of nodes; K: number of links; D, density of connection; C: mean clustering coefficient; H: heterogeneity of connection; L: mean shortest path length; A: assortativity; P, parcellation. Aeto, *Aetosaurus;* AllA, adult *Alligator*; AllJ, juvenile *Alligator*; Arcx, *Archaeopteryx*; Citi, *Citipati*; Coel, *Coelophysis*; Comp, *Compsognathus*; Croc, *Crocodylus*; Dako, *Dakosaurus*; Desm, *Desmatosuchus*; Dibo, *Dibothrosuchus*; Dilo, *Dilophosaurus*; Eora, *Eoraptor*; GalJ, juvenile *Gallus*; GeoJ, juvenile *Geospiza*; Icht, *Ichthyornis*; NotJ, juvenile *Nothura*; Plat, *Plateosaurus*; Psit, *Psittacosaurus*; Rioj, *Riojasuchus*; Sphe, *Sphenosuchus*; Velo, *Velociraptor*.

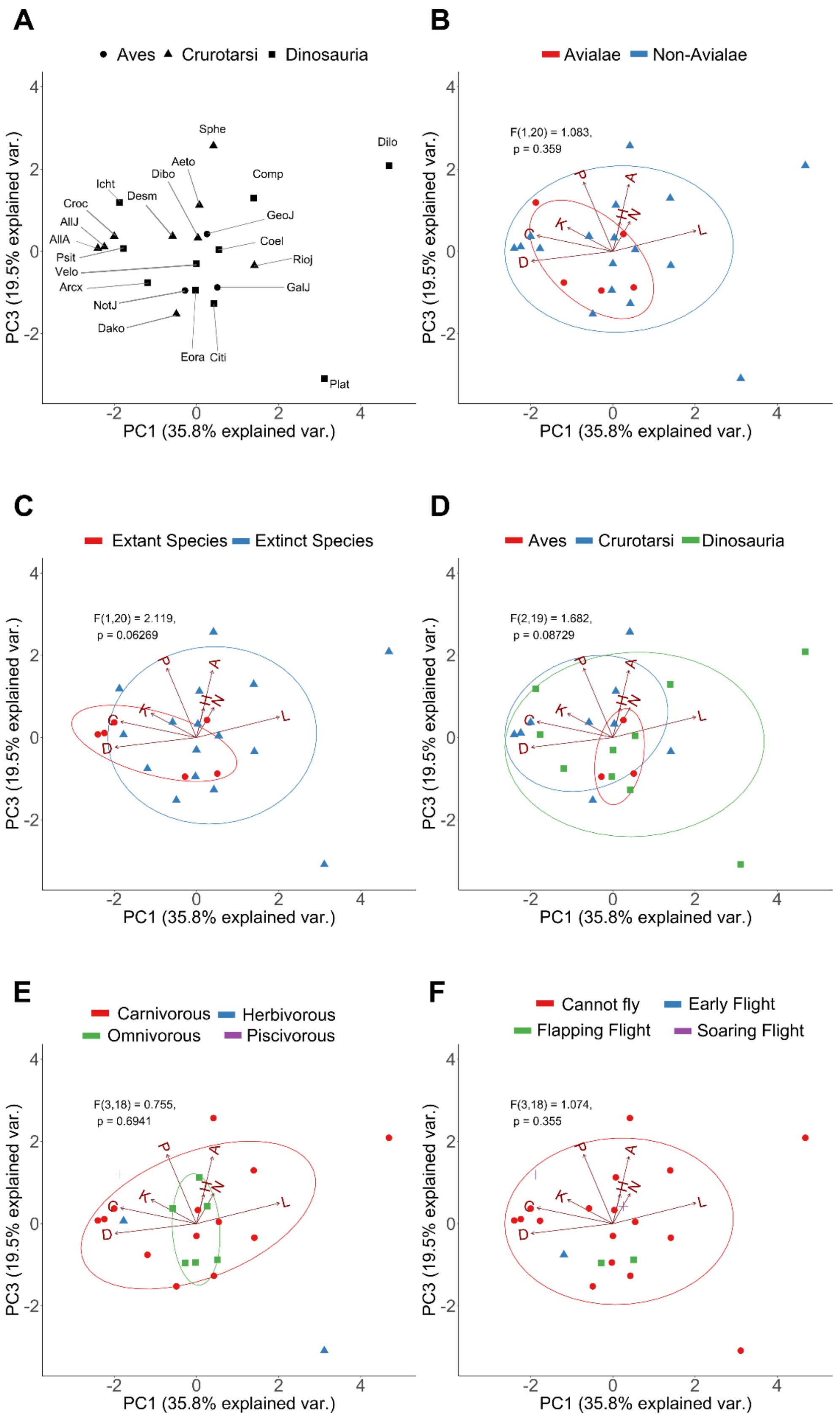

A
Aves
Crurotarsi
Dinosauria
Sphe
Dilo
Aeto
Dibo
Comp
Icht
Desm
Croc
GeoJ
AllJ
Coel
AllA
Psit
Rioj
Velo
Arcx
GalJ
NotJ
Dako
Eora
Citi
Plat
B
Avialae
Non-Avialae
F(1,20) = 1.083,
p = 0.359
C
Extant Species
Extinct Species
F(1,20) = 2.119,
p = 0.06269
D
Aves
Crurotarsi
Dinosauria
F(2,19) = 1.682,
p = 0.08729
E
Carnivorous
Herbivorous
Omnivorous
Piscivorous
F(3,18) = 0.755,
p = 0.6941
F
Cannot fly
Early Flight
Flapping Flight
Soaring Flight
F(3,18) = 1.074,
p = 0.355
PC1 (35.8% explained var.)
PC3 (19.5% explained var.)

**Figure S9. First and third PC of topological parameters for all taxa excluding adult avians.** (A) Skull distribution for each taxon (see labels below). (B) PCA shows an overlap in morphospace occupation between Avialae and non-Avialae species. (C) Extant taxa overlap with extinct taxa. (D) Juvenile avians overlap with crurotarsans and non-avian dinosaurs. (E) Taxa with different dietary requirement overlap with each other. (F) Taxa with different abilities to flight overlap with each other. Ellipses show a normal distribution confidence interval around groups for comparison. Labels: N, number of nodes; K: number of links; D, density of connection; C: mean clustering coefficient; H: heterogeneity of connection; L: mean shortest path length; A: assortativity; P, parcellation. Aeto, *Aetosaurus;* AllA, adult *Alligator*; AllJ, juvenile *Alligator*; Arcx, *Archaeopteryx*; Citi, *Citipati*; Coel, *Coelophysis*; Comp, *Compsognathus*; Croc, *Crocodylus*; Dako, *Dakosaurus*; Desm, *Desmatosuchus*; Dibo, *Dibothrosuchus*; Dilo, *Dilophosaurus*; Eora, *Eoraptor*; GalJ, juvenile *Gallus*; GeoJ, juvenile *Geospiza*; Icht, *Ichthyornis*; NotJ, juvenile *Nothura*; Plat, *Plateosaurus*; Psit, *Psittacosaurus*; Rioj, *Riojasuchus*; Sphe, *Sphenosuchus*; Velo, *Velociraptor*.

*Riojasuchus*

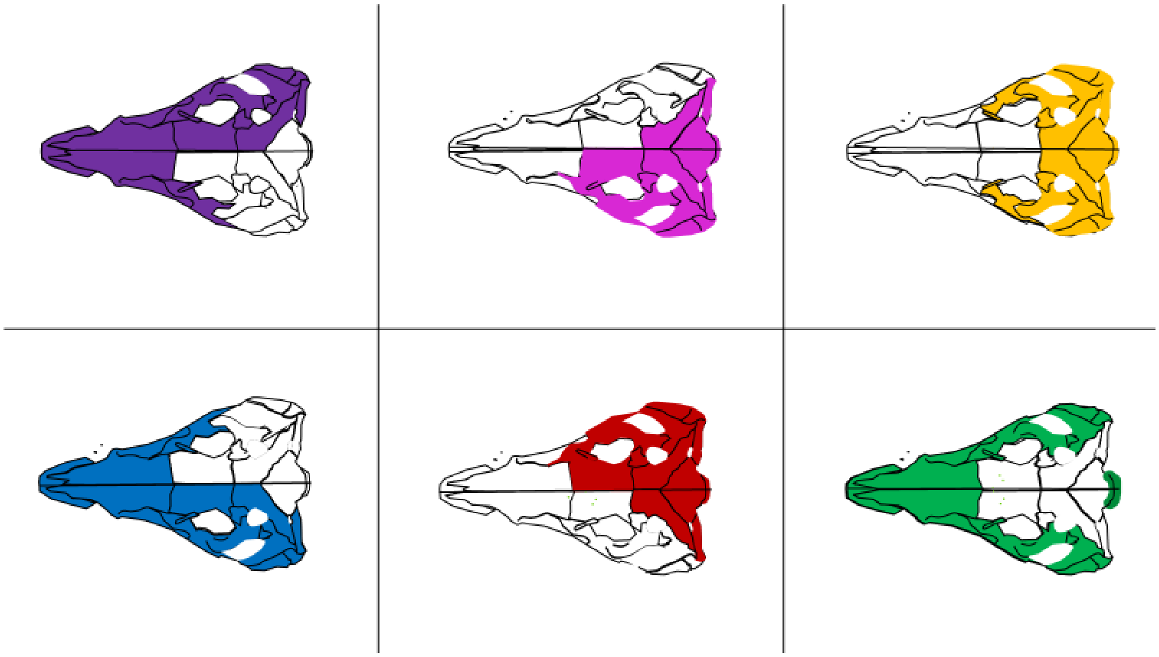

*Aetosaurus*

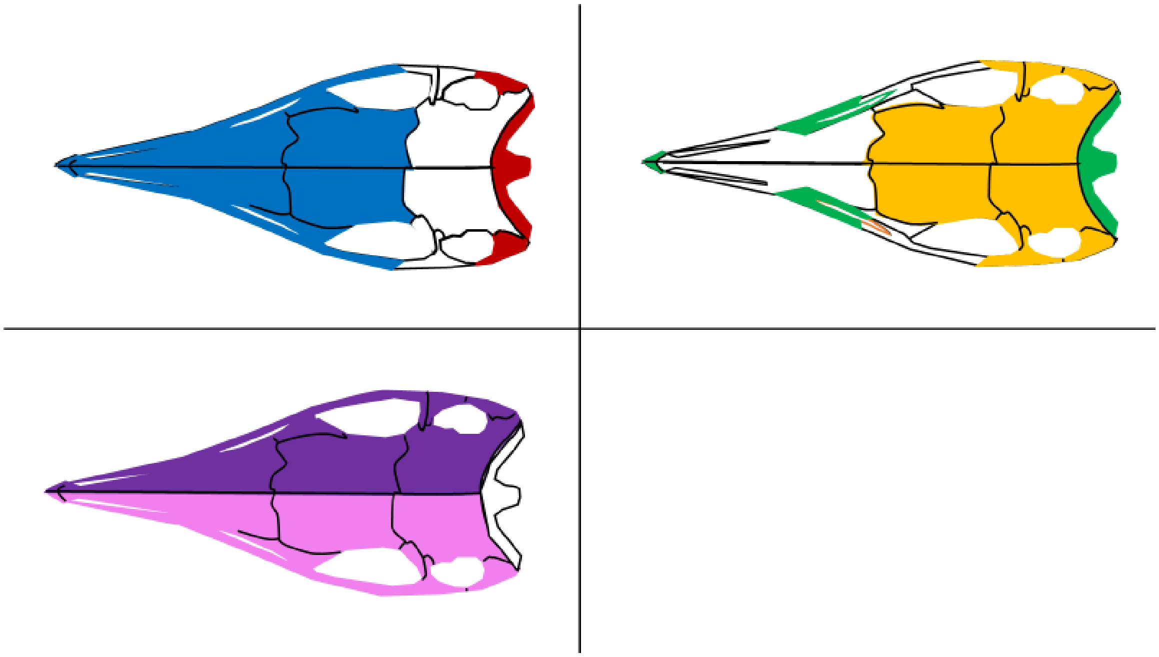

*Desmatosuchus*

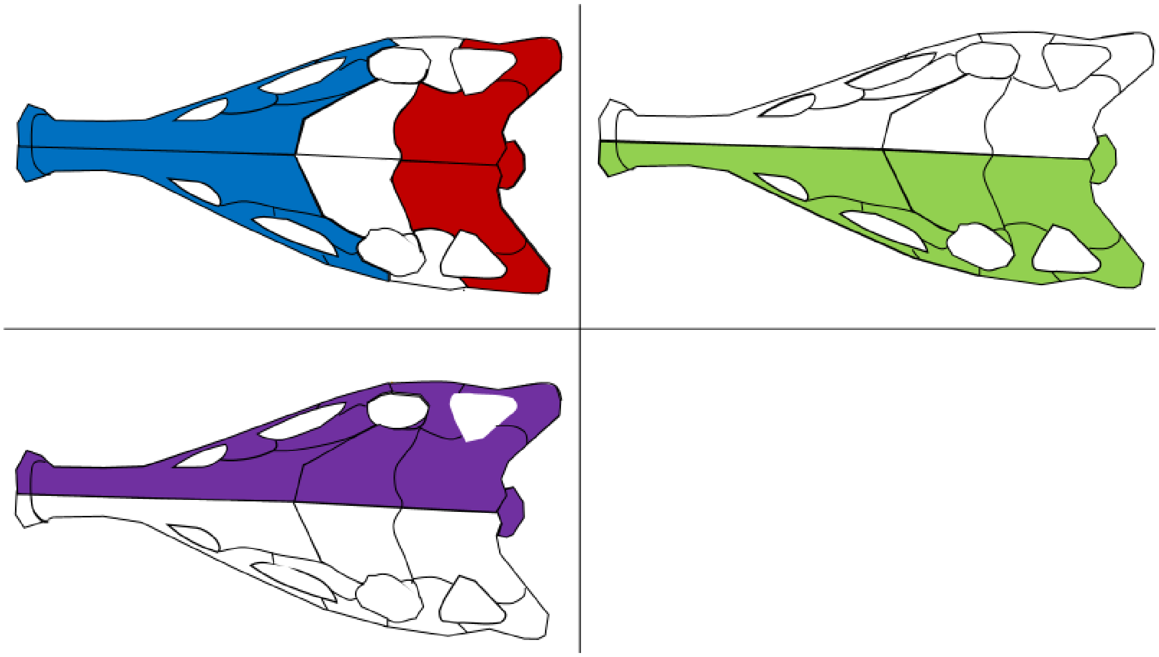

*Sphenosuchus*

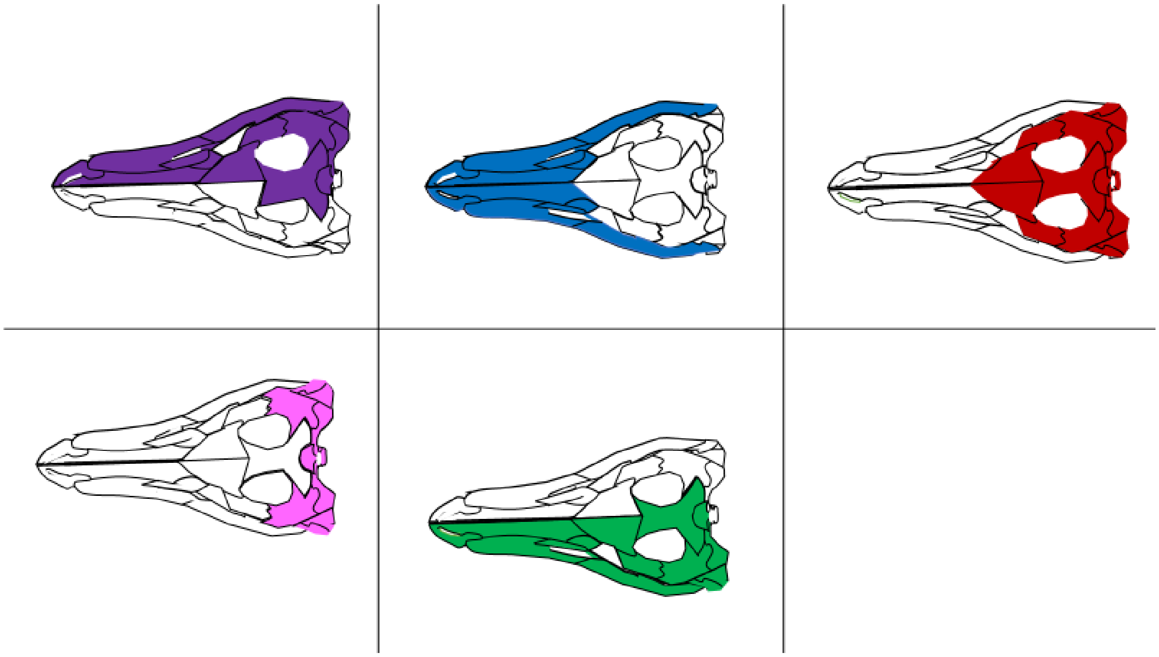

*Dibothrosuchus*

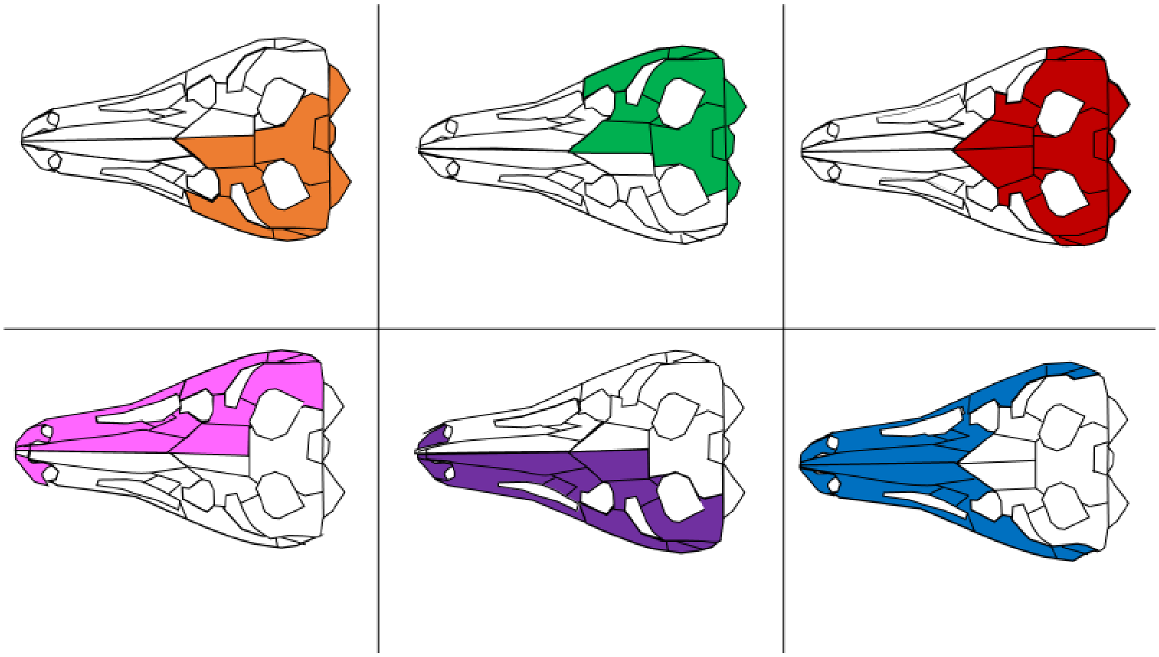

*Dakosaurus*

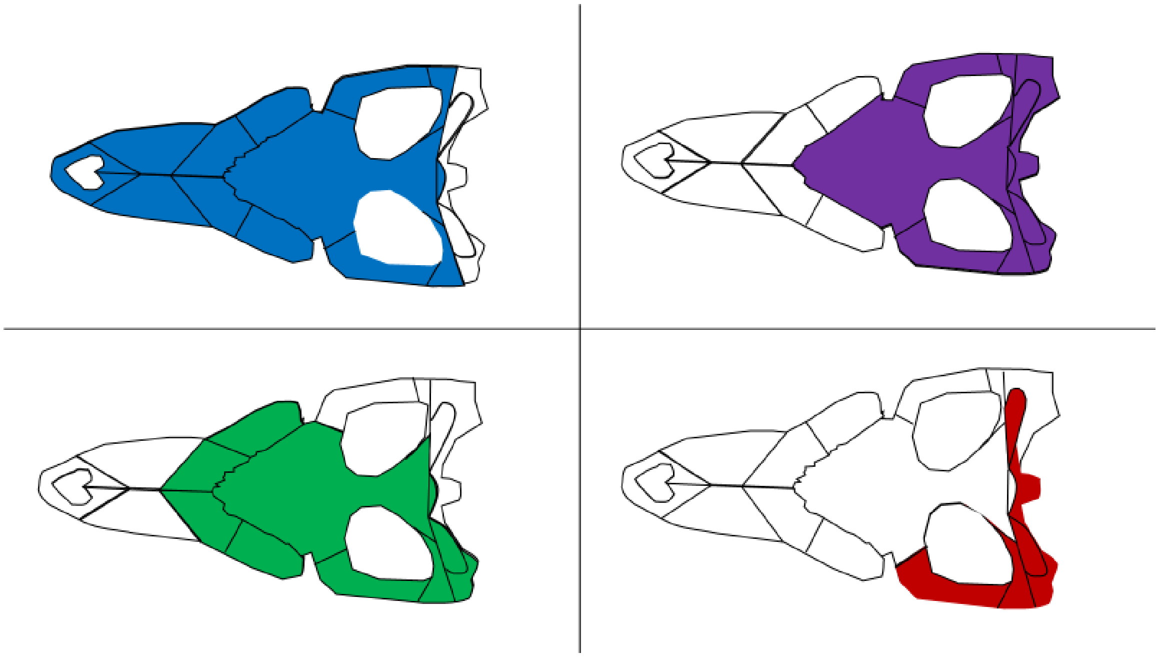

*Alligator* (adult)

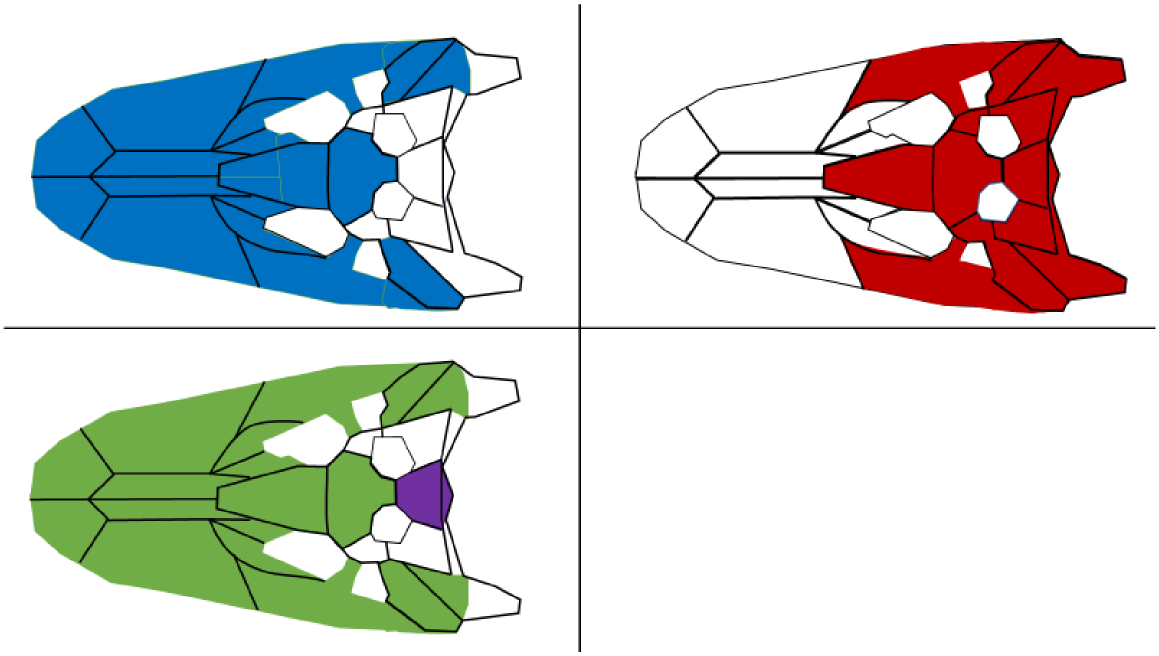

*Alligator* (juvenile)

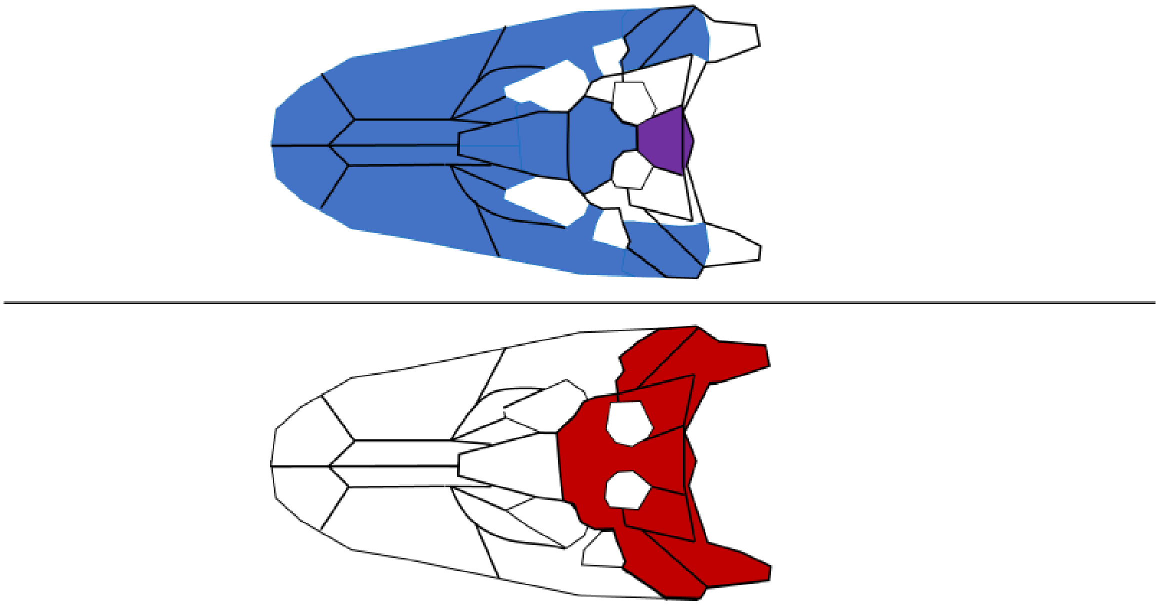

*Crocodylus*

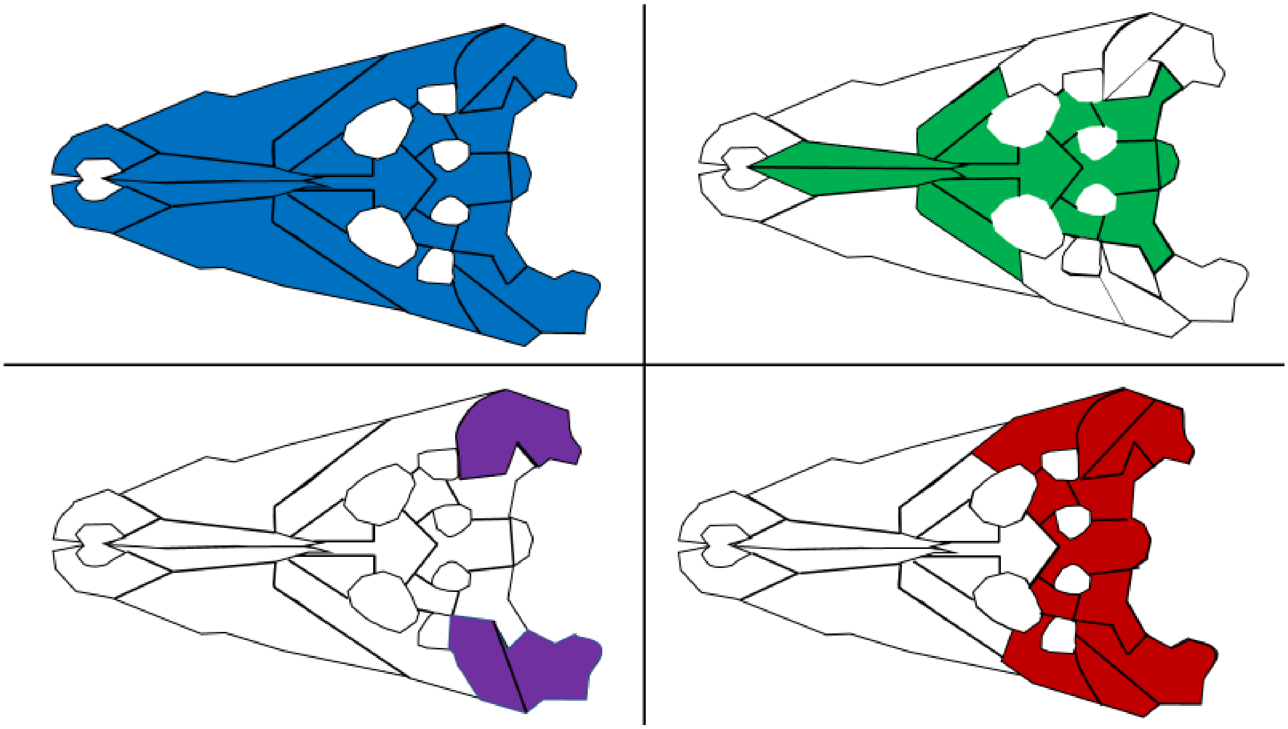

*Psittacosaurus*

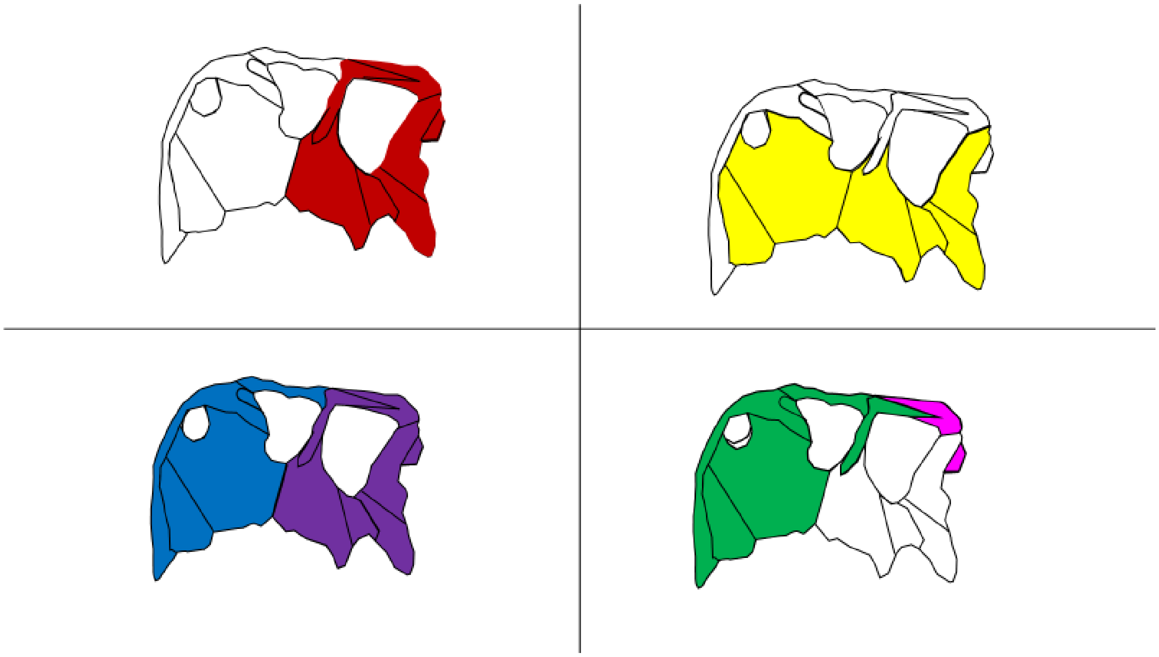

*Eoraptor*

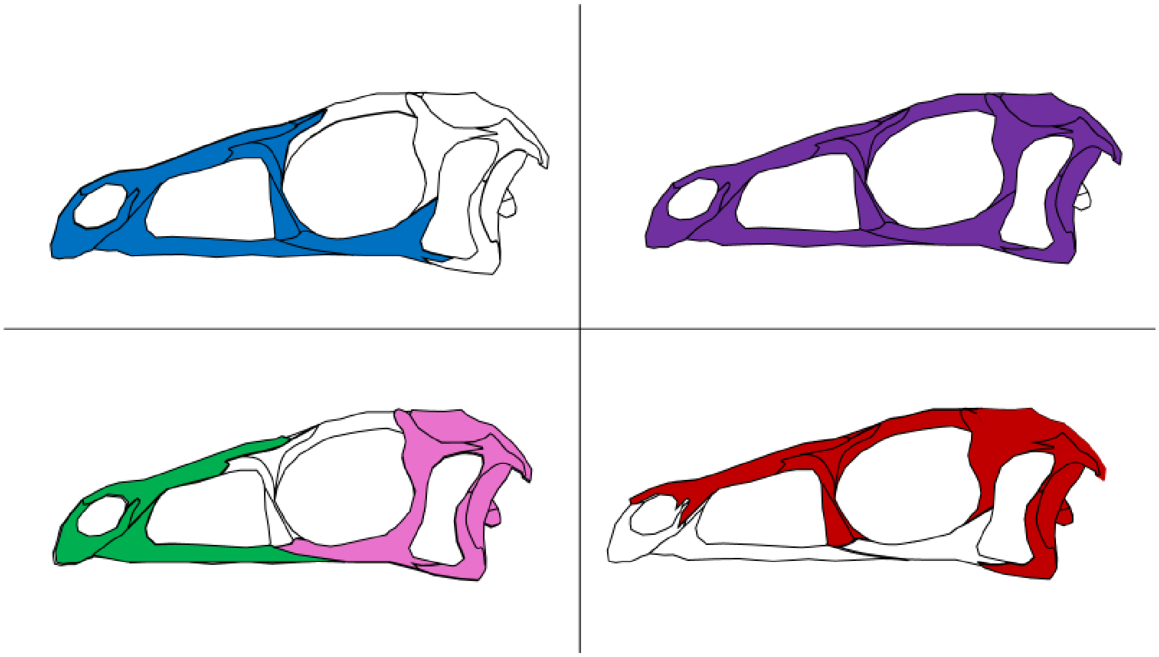

*Plateosaurus*

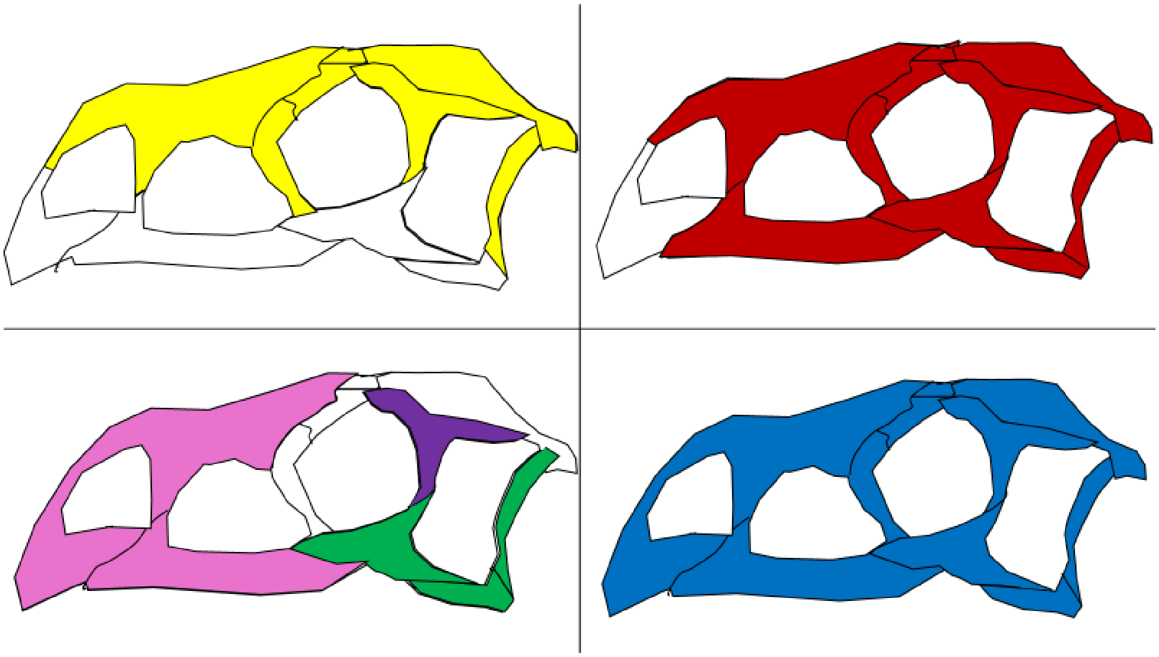

*Coelophysis*

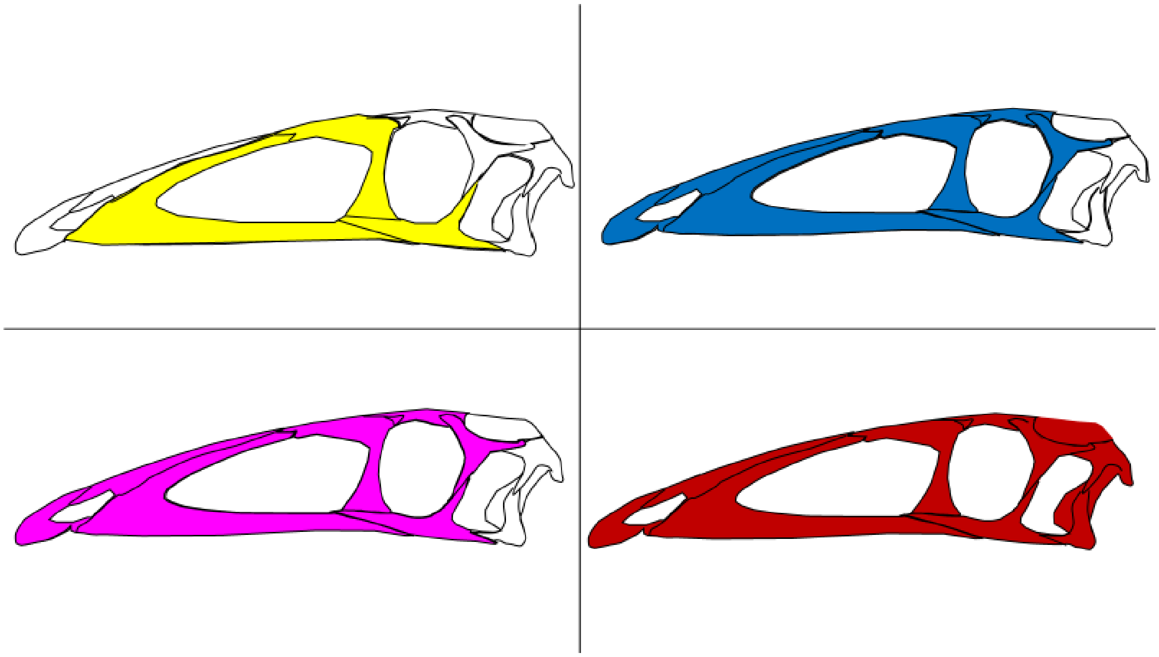

*Dilophosaurus*

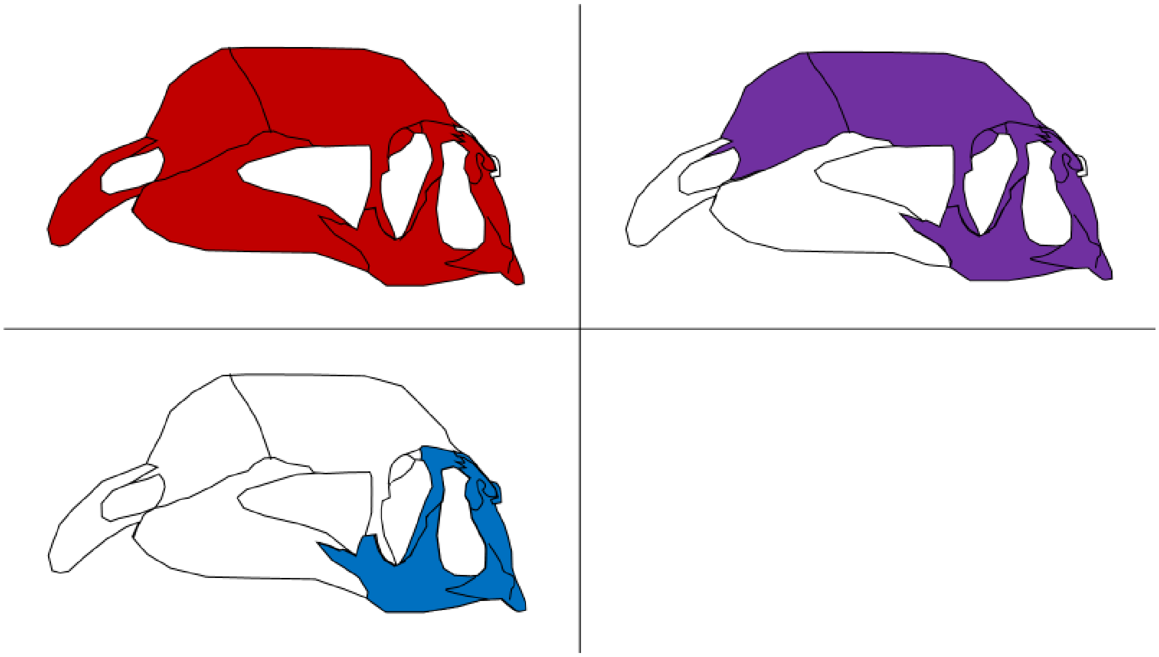

*Compsognathus*

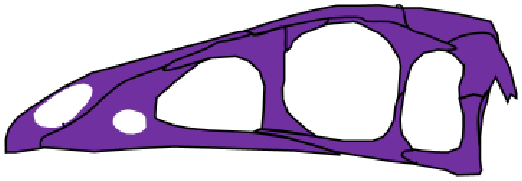

---

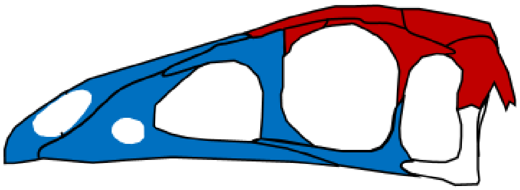

*Citipati*

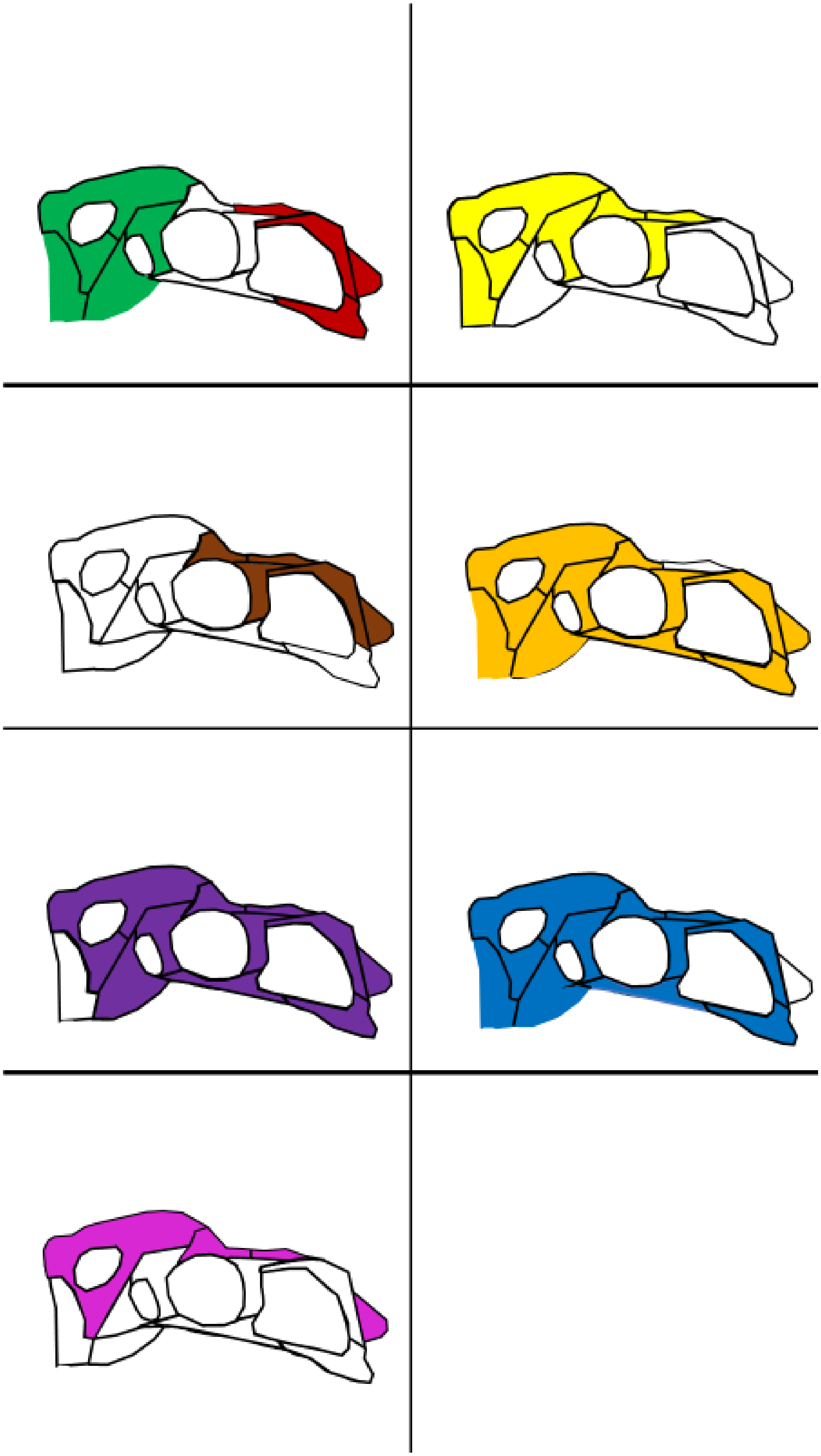

*Velociraptor*

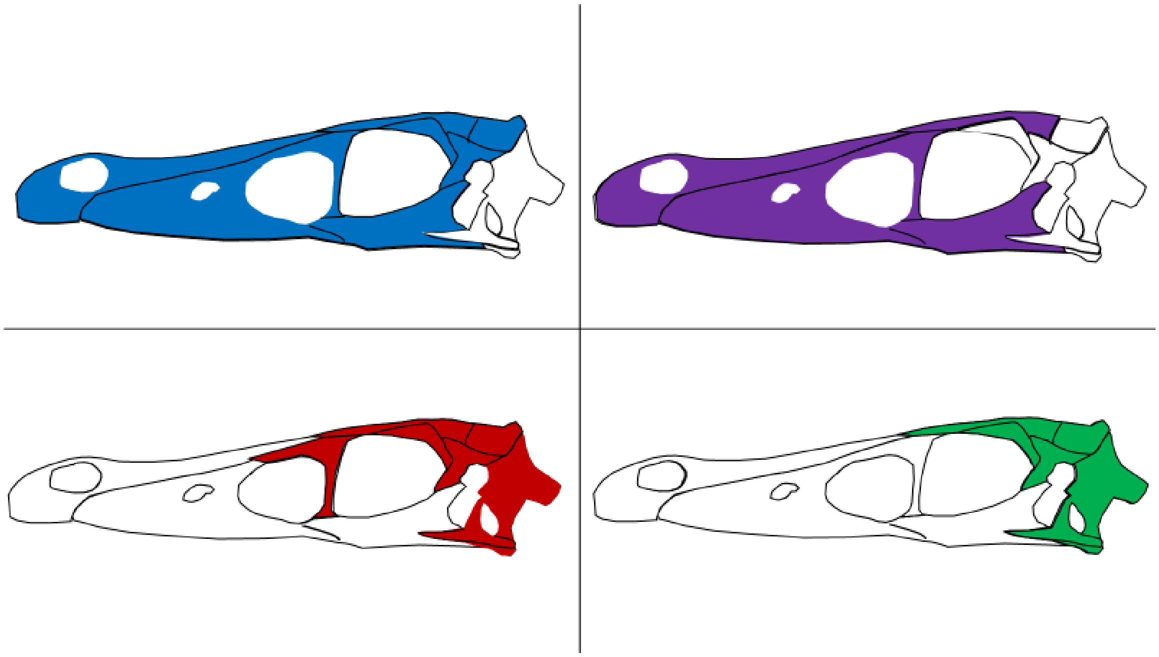

*Archaeopteryx*

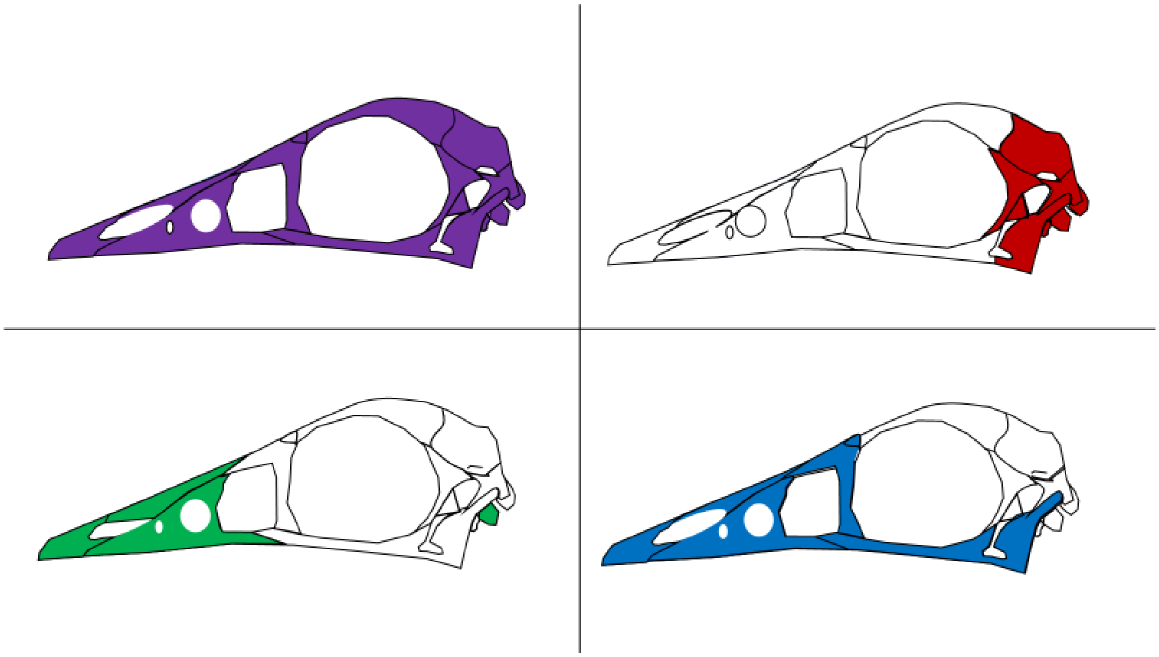

*Ichthyornis*

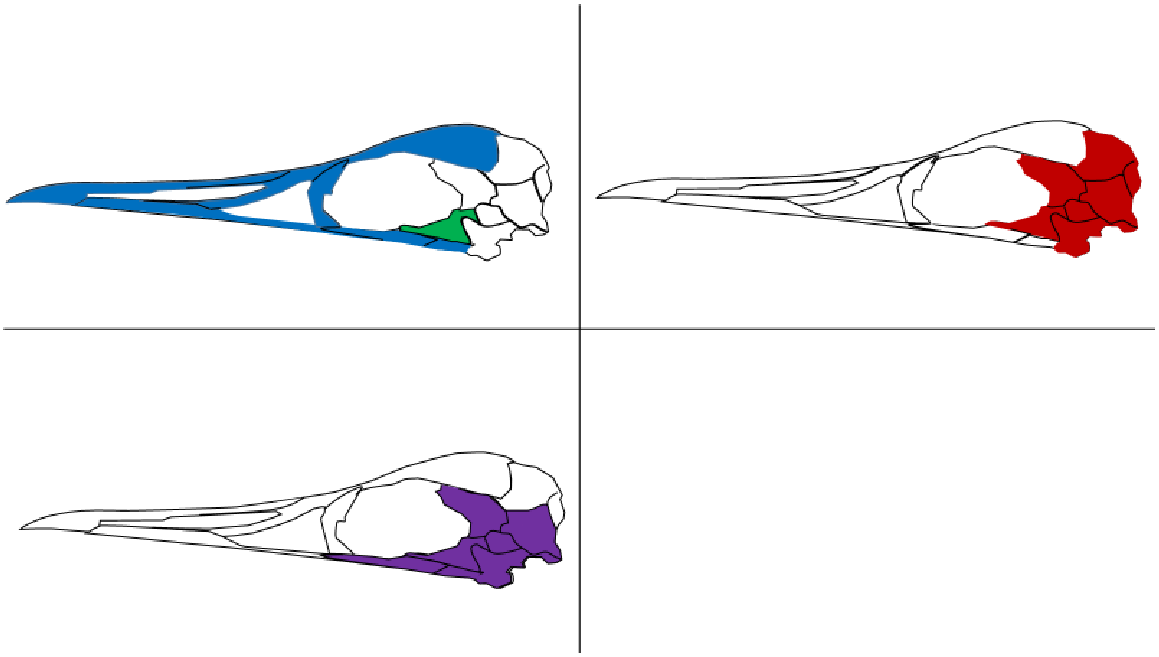

*Nothura (*adult)

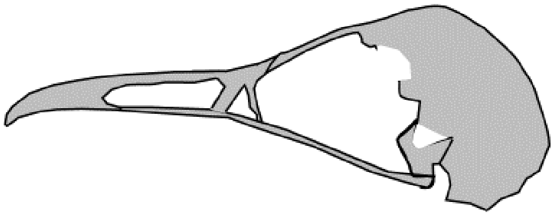

*Nothura* (juvenile)

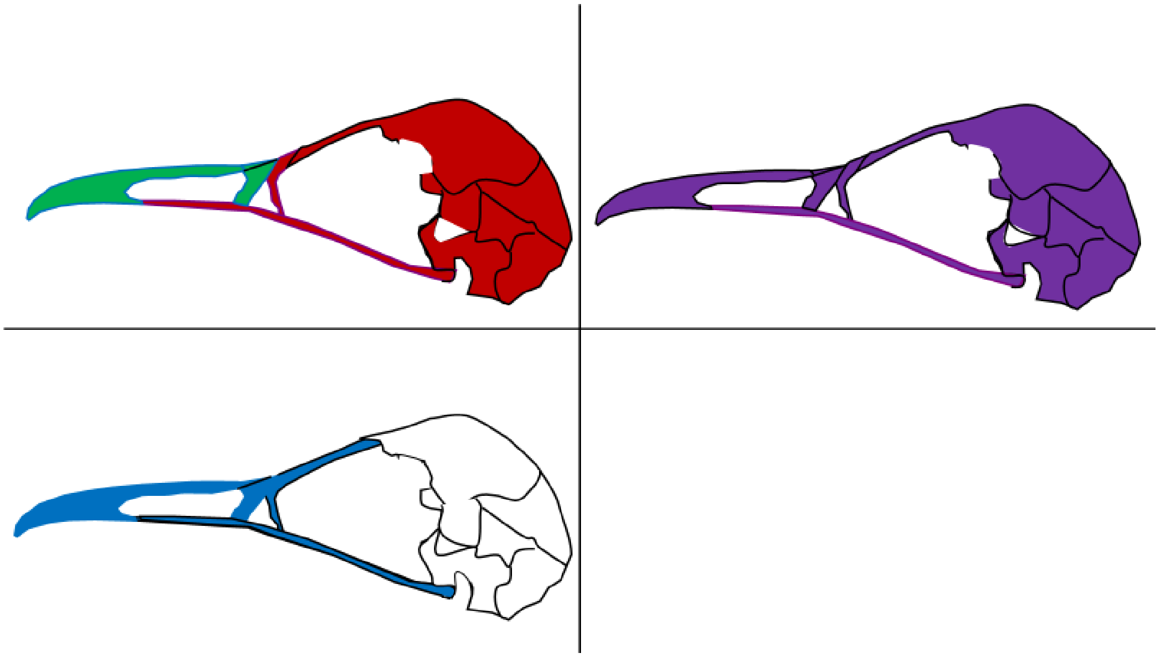

*Gallus* (adult)

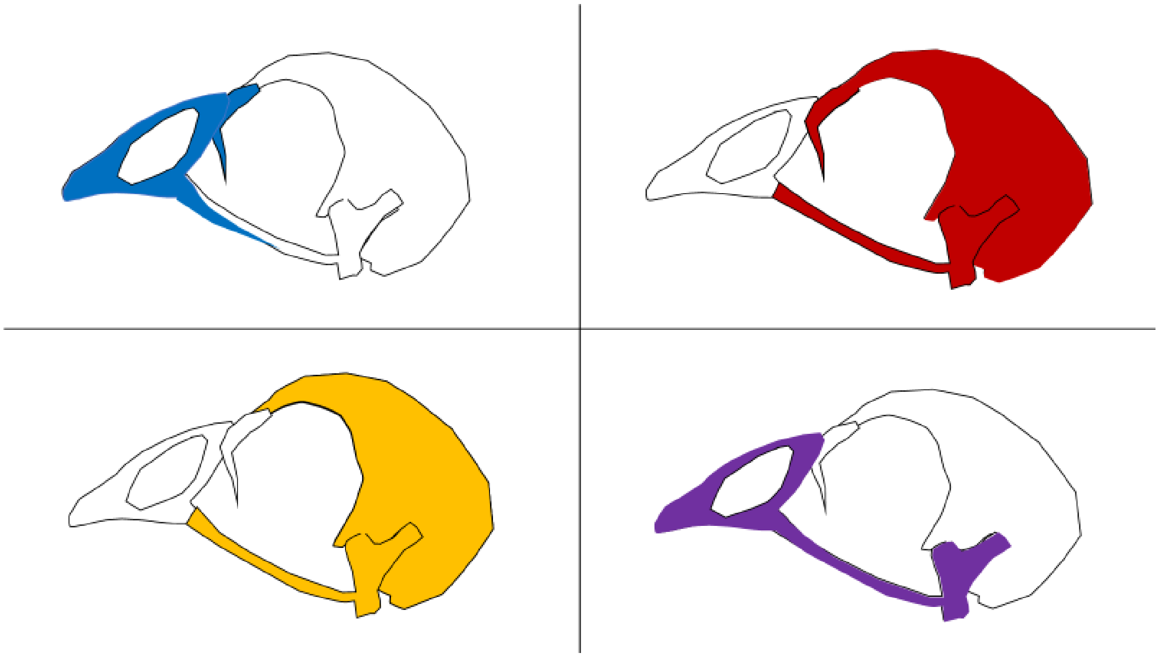

*Gallus* (juvenile)

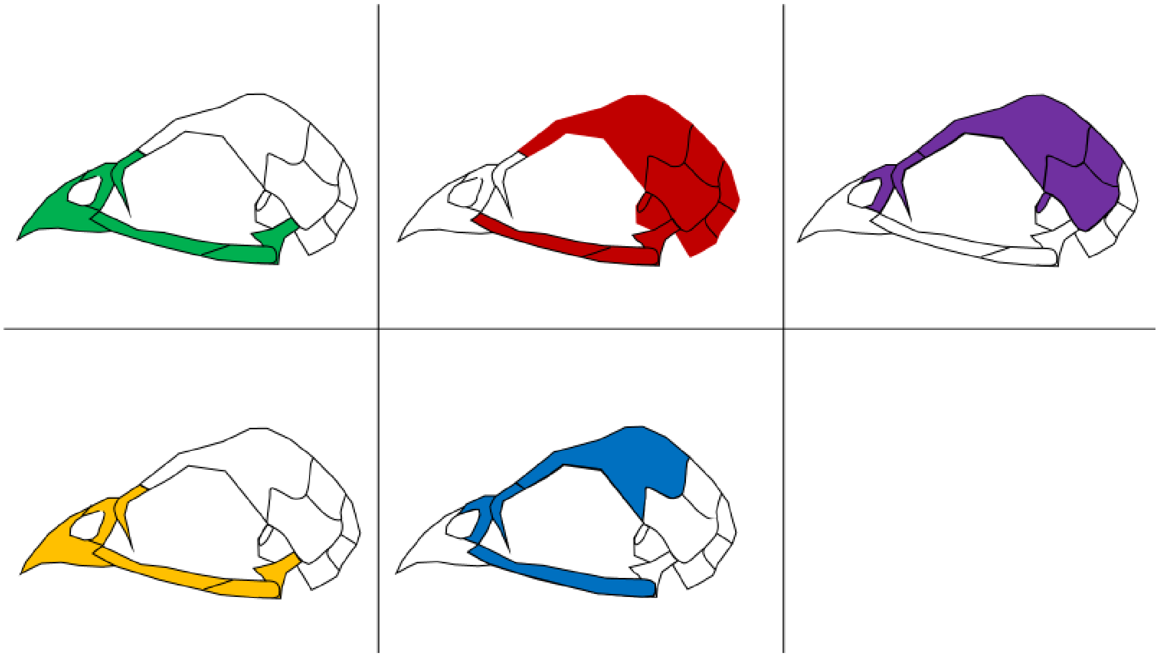

*Geospiza* (adult)

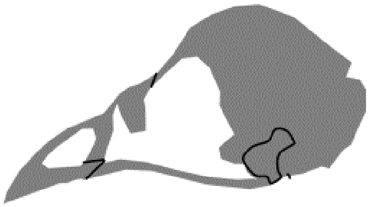

*Geospiza* (juvenile)

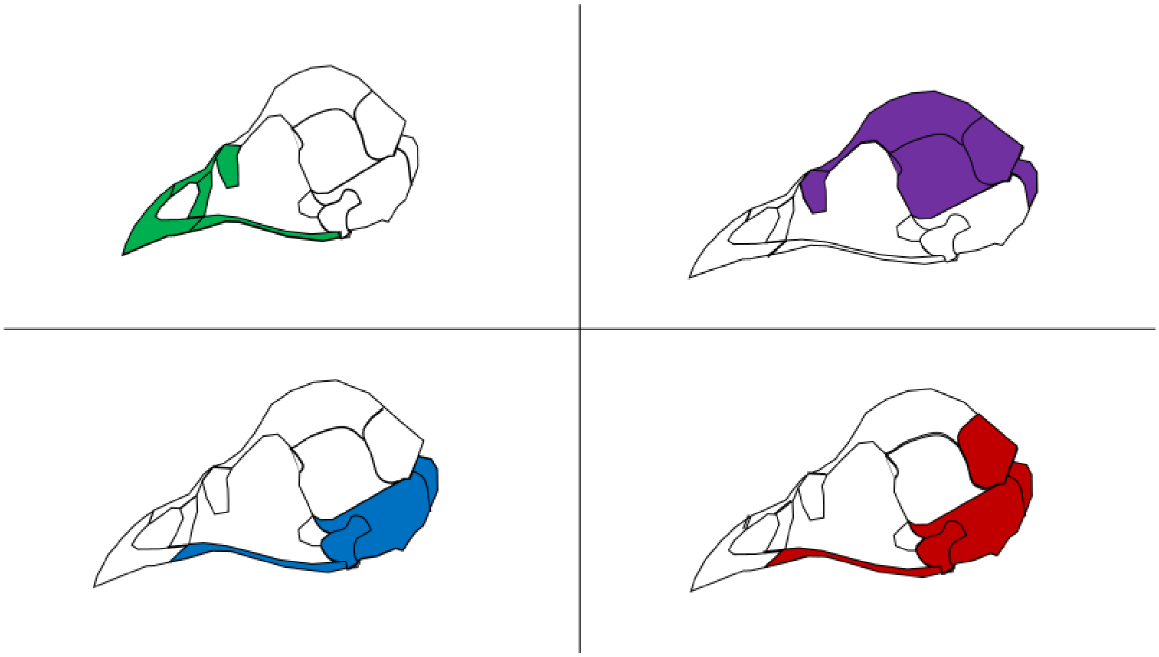

**Figure S10. Node-based modules of archosaurs based on details listed on Table S4.**

**Supplementary Information 1: References and notes about the specimens (arranged alphabetically) used to create the matrices for the analysis.**

*Aetosaurus ferratus* was coded by using the specimen SMNS 5770(S-16, S-18) from Schoch[19].

Adult and juvenile matrices of *Alligator mississippiensis* were coded by Jake Horton[11]. The adult skull was from specimen 1873.2.21.2 from Patrick Campbell at the Natural History Museum, London (NHM) and Iordansky[20]. The juvenile skull was from the 3D model created by Witmer's lab (2011, specimen OUVC 10606, The Visible Interactive Alligator Project at Ohio University and the University of Missouri, url:

https://people.ohio.edu/witmerl/3D_gator.htm;), Witmer and Ridgely[21], and Dufeau and Witmer[22].

*Archaeopteryx lithographica* was coded following the reconstructions by Elzanowski[4] and Rauhut[5] of the London (1st), Munich (7th), Berlin (2nd), and Eichstätt (5th) specimens. No exoccipital, postfrontal, opisthotic, prefrontal, nor supraoccipital were observed. The Eichstätt (5th) specimen was later confirmed by Smith-Paredes et al.[6] to have prefontal which articulates with frontal, nasal, and lacrimal.

*Citipati osmolskae* was coded following the details described by Clark et al.[23]; Clark, Norell and Rowe[24]; and Osmolska et al.[25]. The prefrontal was absent. The following were fused together: basisphenoid, basioccipital, opisthotic, and prootic; laterosphenoid and prootic (labelled as “basisphenoid”). The left and right sides of the following were fused together: supraoccipital, nasal, premaxilla, vomer, and parietal. Clark, Norell and Rowe suggested basipterygoid (although broken) to articulate with pterygoid and basisphenoid[24].

*Coelophysis bauri* was coded following the details described by Witmer[26]; Tykoski and Rowe[27]; Bhullar et al.[28]; Rinehart et al.[29]; and Spielmann et al.[30]. No separate supraoccipital, epiotic, nor coronoid were recorded. The left and right vomers were fused as one. Exoccipital was fused with opisthotic (labelled as “paroccipital process”)

*Compsognathus longipes* was coded by using the holotype B.S.P. A.S. I 563 and MNHN CNJ 79 from Peyer[31]. The basisphenoid, basiooccipital, opisthotic, postfrontal, exoccipital, and supraoccipital were absent.

*Crocodylus moreletii* was coded by Jake Horton[11] based on specimen 1861.4.1.4 from Patrick Campbell at the Natural History Museum, London (NHM) and Iordansky[20].

*Dakosaurus andiniensis* was coded by using the specimen MOZ 6146P from Gasparini, Pol and Spalletti[32] and Pol and Gasparini[33]. It was described not to have separate vomer, laterosphenoids, prootics, postfrontals, and palpebral; premaxilla not in contact with nasal is a common feature in thalattosuchians; and ectopyerygoids were obscured (omitted from analysis). Basisphenoid was described to be very thin and concave-shaped between pterygoid and basioccipital. No separate opisthotics were observed from exoccipital (labelled as “exoccipital”).

*Desmatosuchus haplocerus* was coded by using specimens TTUP 9023, TTUP 9024, and UCMP 27408 from Small[34]. Parasphenoid was fused with basisphenoid (labelled as “basisphenoid”). Parietal was fused with supraoccipital, laterosphenoid, prootic and opisthotic (labelled as “parietal”).

*Dibothrosuchus elaphros* was coded following the reconstruction by Wu and Chaterjee[35] based on specimen IVP V 7907 and holotype CUP 2981, which was also fully described by Simmons[36]. Exoccipital was fused with opisthotic (labelled as “paroccipital process”).

*Dilophosaurus wetherilli* was coded by using the specimens UCMP37302 and UCMP37303. Details were described by Welles[37] and Tykoski and Rowe[27]. It was synonymous to *Megalosaurus wetherilli*. Information about the pterygoid and the absence of vomer were from *Coelophysis* described by Tykoski and Rowe[27]. Postfrontal was not observed. The frontal and parietal were damaged in *Dilophosaurus,* so Welles (1984) reconstructed it without showing the contact between parietal and frontal.

*Eoraptor lunensis* was coded following the details described by Sereno et al.[38] and Sereno[39]. Vomer was attached to nasal and premaxilla. Sereno et al.[38] postulated that the vomer articulates with the pterygoids. Because there was no evidence that the vomer is a paired bone, the vomer is recorded as one bone. The vomer was too fragmented and there was no evidence for vomer-palatine articulation, so we recorded no connection between palatine and vomer. Opisthotic and exoccipital were fused as paroccipital process. Laterosphenoid was too fractured and its articulation to postorbitals was conjectured by Sereno et al.[38].

*Gallus gallus* (Adult) matrix has been borrowed from a previous study by Werneburg et al.[40].

*Gallus gallus* (Juvenile) matrix was coded following the details described by Jollie[41] and van Den Heuvel[42]. The juvenile used in Jollie[41] was 2-3 days old. The following were either fused together or no visible sutures were observed: epiotic and supraoccipital (labelled in this analysis as "supraoccipital"); opisthotic and exoccipital (as "exoccipital"); rostrosphenoid, basisphenoid, basiparasphenoid, alaparasphenoid, and sellaparasphenoid (labelled as "basisphenoid"); lateral ethmoid and prefrontal (as "prefrontal" ); palatine and pterygoids (called "pterygopalatine”).

*Geospiza fortis* (Adult) matrix was coded following the details described by Genbrugge et al.[43] and specimen USNM 345593 from Smithsonian Institution, National Museum of Natural History, Division of Birds, accessed on http://phenome10k.org/geospiza-fortis/ The following were fused together: nasal, premaxilla, and maxilla (labelled in this analysis as "Upper Beak"); palatine and vomer (as "Palate").

*Geospiza fortis* (Juvenile) matrix was coded following the details described by Genbrugge et al.[44]. The juvenile used here was from the group 2 nestling size class 1. The following were fused together: basisphenoid and parasphenoid (as "basisphenoid"); prootic, opisthotic, and exoccipital (as "exoccipital"); Jugal was fused to quadratojugal.

*Ichthyornis dispar* was coded following the details of the holotype YPM 1450 and specimens FHSM VP-18702, ALMNH 3316, KUVP 119673, and BHI 6421 described by Field et al.[45]. There was no postorbital, prefrontal nor postfrontal.

*Nothura maculosa* (Adult) was coded following the details of the specimen MHNT06 from Silveira and Höfling[46]. The following were fused together: paraoccipital, basioccipital, exoccipital, basisphenoid, parietal, supraoccipital, parietal and squamosa (labelled in this analysis as "Braincase"); frontal, lacrimal, nasal, premaxilla, maxilla, frontal, palatine, vomer, laterosphenoid, pterygoid, mesethmoid, and basipterygoid (as "Beak"); maxilla, jugal, and quadratojugal (as "Jugal Bar").

*Nothura maculosa* (Juvenile) was coded following the details of the specimen AZ163 from Silveira and Höfling[46]. Palatine was fused to maxilla here (labelled in this analysis as "maxilla").

*Plateosaurus engelhardti* was coded following the reconstruction of MB.R.1937 and other sauropodomorphs by Button, Barrett and Rayfield[47] and Huene[48]. Huene reconstructed the vomer, palatine, pterygoid and ectopterygoid from *P. longiceps*[48].

Labelling of the bones is based on the drawing by Huene[48] and *P. erlenbergiensis* AMNH FARB 6810 from Prieto-Marquez & Norell[49]. Exoccipital is fused to opisthotic (labelled as "opisthotic"). No postfrontal observed. Button, Barrett and Rayfield[47] mentioned the presence of an epipterygoid without describing its articulations but a separate epipterygoid was not observed in the figures and illustrations from Button, Barrett and Rayfield[50] and Huene[48]. Thus, the epipterygoid was not recorded as a separate bone in the matrix.

*Psittacosaurus lujiatunensis* was coded following the details described by Zhou et al.[51] (2006) and Osborn[52]. Opisthotic and exoccipital were fused together (labelled as “paroccipital process”). Zhou et al.[51] referred to Sereno[53] for more details on the laterosphenoid articulation. Also, specimens in Zhou et al (2006) show conflicting articulations between the left and right maxilla: in (ZMNH M8137 (holotype), the bones are separate but in PKUP V1053 (paratype), the bones are joined together. We recorded the left and right maxilla are fused as one bone.

*Riojasuchus tenuisceps* was coded following the details of the holotype PVL3827 and specimen PVL 3828 described by von Baczko and Desojo[54]. Exoccipital was fused to opisthotic (labelled in this analysis as "opisthotic"). The premaxilla appear to be on a different plane from maxilla but we assume the premaxilla articulates with the maxilla here.

*Sphenosuchus acutus* was coded following the description of the holotype SAM 3014 by Walker[55]. Its quadrates do not meet opisthotic nor latersosphenoid but meet prootic and squamosal. Exoccipitals were fused with opisthotics (labelled as “paroccipital process”).

*Velociraptor mongoliensis* was coded following the description of the specimens GIN 100/24, GIN 100/25, and GIN 100/2000 by Barsbold and Osmolska[56] and Norell et al.[57]. The following were fused together: prefrontal and lacrimal (marked as"prefrontal"); parasphenoid and basisphenoid (marked as "basisphenoid"); basioccipital and opisthotic (marked as "basioccipital"). The suture between exoccipital and supraoccipital was uncertain (these two bones could be fused) and marked as “1”.

**Supplementary Information 2: Comparison between node-based modules and variational modules in archosaurs.**

Previous studies on shape co-variation (i.e., variational modularity) in the skull of archosaurs by Felice and colleagues[8] using EMMLi showed that the composition of the rostral module (premaxilla, maxilla, nasal) and the neurocranium module (parietal, frontal, postorbital, and squamosal) were conserved throughout archosaurs. We found a similar composition for both rostral and cranial modules.

Avians used in this analysis did not have a separate postorbital. The rostral and cranium modules in adult birds used in this analysis and by Felice et al.[8], were comprised of fused bones (Table S4).

We showed that facial bones (such as frontal, prefrontal, lacrimal, and premaxilla) that co-vary more in crocodylians[8] are grouped in at least one node-based module in archosaurs. The frontal bones could be found in both rostral and neurocranial node-based modules.

We did not observe a separated prefrontal bone in *Citipati* to *Geospiza* (Table S6), which agrees with previous observations showing that a separate prefrontal was an ancestral trait preserved in extant crurotarsans and regained in the adult chicken and *Archaeopteryx*[6].

Additionally, Felice and colleagues showed that birds, non-avian dinosaurs, and crocodilians had an occipital module comprised of the same elements (supraoccipital and basioccipital)[8]. Supraoccipital and basioccipital belonged to the same node-based module in most of the archosaurs in this analysis. Supraoccipital was not separated from other bones in *Aetosaurus*. Supraoccipital and basisphenoid were not identified in *Compsognathus*; and they were fused together in adult birds.

**Supplementary Information 3: Comparison of network parameters among Aves, Crurotarsi, and non-avian Dinosauria.**

Network parameters were calculated for each skull network, with median and range (in parenthesis) values calculated for each class (see Table 1).

Number of bones (nodes) and articulations (links) varied greatly across archosaurs: N ranged from 6 in adult *Nothura* to 46 in *Sphenosuchus*; K ranged from 9 in adult *Nothura* to 114 in *Sphenosuchus*. Not all archosaurs with the highest N value had the highest K value, such as *Dilophosaurus* (N: 44, K: 72). Aves had fewer nodes and links (N: 6 to 34, K: 9 to 72), associated with a highly fused skull with fewer visible articulations, compared to Crurotarsi (N: 28 to 46, K: 56 to 114) and to non-avian Dinosauria (N: 32 to 44, K: 61 to 112) (Table S10, in bold).

Density of connections (D) ranged from 0.076 (*Dilophosaurus*) to 0.6 (adult *Nothura*). Among clades, the density in non-avian Dinosauria (0.076 - 0.159) and Crurotarsi (0.095 to 0.159) were lower than Aves (0.128 to 0.6). Thus, Aves had a more connected network despite fewer articulations.

Mean clustering coefficient (C) ranged from 0.085 (*Plateosaurus*) to 0.733 (adult *Nothura*). Among clades, the minimal mean clustering coefficient for non-avian Dinosauria was the lowest (0.085), followed by Aves (0.126) and Crurotarsi (0.266) while the maximal C was non-avian Dinosauria (0.422), Crurotarsi (0.45), and Aves (0.733) in ascending order. It means neighboring bones in Aves and Crurotarsi are more likely to act as clusters and have correlating functions and structures[59–61].

Mean path length (L) ranged from 1.4 (*Nothura)* to 3.981 (*Dilophosaurus)*. The mean path length for Aves was the lowest (1.4 to 2.877), followed by Crurotarsi (2.429 to 3.278) and non-avian Dinosauria (2.509 to 3.981). It shows Aves have higher efficiency to pass biomechanical forces and molecular signals to other bones[62,63] than non-avian dinosaurs.

D, C, and L are parameters that delineate the relationship between nodes (bones) and links (articulations) and can be used to measure complexity[63]. Avian skulls that have higher D and C and lower L are more complex than non-avian dinosaurs.

Heterogeneity of connections (H), which means the difference in the number of connections each bone has, ranged from 0.284 (*Aetosaurus)* to 0.634 (adult *Gallus*). Average H value for Crurotarsi was the lowest (0.284 to 0.471), followed by non-avian

Dinosauria (0.272 to 0.439) and Aves (0.296 to 0.634). High H means that anisomerism, the specialization and anatomical difference of bones had occurred[63,64].

Assortativity of connections (A) ranged from -0.455 (adult *Nothura*) to 0.282 (*Compsognathus*). The negative assortativity in Aves (-0.455 to 0.093) meant bones with were more likely to connect to bones with a different number of articulations than Crurotarsi (-0.217 to 0.185). In contrast, nodes with the same number of connections were likely to connect to each other in non-avian Dinosauria (-0.36 to 0.282). Together, A and H mean modern bird skulls are composed of bones with a different number of articulations while crurotarsans have a more regularly (or symmetrically) shaped skull.

Parcellation (P) ranged from 0 to 0.886 (juvenile *Geospiza*). Aves has the largest range for maximal value for parcellation (0 to 0.886), followed by non-avian Dinosauria (0.096 to 0.841), and Crurotarsi (0.422 - 0.799). Adult *Nothura* and adult *Geospiza* (P: 0) only had one module, as restricted by fewer nodes and articulation.

**Supplementary Information 4: Comparison based on diet.**

To test whether diet had an impact in the structural organization of the archosaur skull, we grouped the taxa of our sample in carnivores, herbivores, and omnivores: *A. mississippiensis, Archaeopteryx lithographica, Citipati osmolskae, Coelophysis bauri, Compsognathus longipes, Crocodylus moreletii, Dakosaurus andiniensis, Dibothrosuchus elaphros, Dilophosaurus wetherilli, Riojasuchus tenuisceps, Sphenosuchus acutus,* and *Velociraptor mongoliensis* are carnivores[65–68]. *Plateosaurus* and *Psittacosaurus* are herbivores[65,69]; modern birds used in this analysis, *Aetosaurus ferratus, Eoraptor lunensis* and *Dilophosaurus haplocerus* are omnivores[8,38,70,71]; *Ichthyornis dispar* was recorded as piscivore because there was no evidence that it had other sources of food[72]. Some carnivores also consumed fish, such as *A. mississippiensis*. Whether aetosaurs, such as *A. ferratus* and *D. haplocerus*, are either herbivores, omnivores, or insectivores is still on debate[71,73–76] and we have classified them as omnivores in our analysis.

No significant difference was found between archosaurs with different dietary requirements when Aves were included ($F_{3,21} = 1.329$, $p = 0.2225$; Fig. S1E), when Aves were excluded ($F_{3,15} = 0.512$, $p = 0.9051$; Fig. S4E), and when adult Aves were excluded ($F_{3,18} = 0.755$, $p = 0.6842$; Fig. S7E; Table S11).

**Supplementary Information 5: Comparison of juvenile avian modules with adult avian bones.**

The density, mean path length, parcellation, and heterogeneity (D: 0.128 to 0.143, L: 2.717 to 2.877, P: 0.446 to 0.886, H:0.296 to 0.48) of juvenile birds were closer to Crurotarsi (D: 0.095 to 0.159, L: 2.429 to 3.278, P: 0.422 to 0.799, H:0.284 to 0.471) and non-avian Dinosauria (D: 0.076 to 0.159, L: 2.509 to 3.981, P: 0.096 to 0.841, H: 0.272 to 0.439), meaning they have a complexity and a network symmetry similar to crurotarsans and its theropod ancestors.

The mean clustering coefficient of juvenile birds (C: 0.263 to 0.348) overlapped with adult birds (C: 0.126 to 0.733), Crurotarsi (C: 0.266 to 0.45) and non-avian Dinosauria (C: 0.085 to 0.422), meaning bones at juvenile stages first cluster with each other became more integrated with each other) before suture fusion. As they age, bird skulls became less symmetrical and their bones became more connected and closer to each other.

For the spotted tinamou, juvenile jugal and quadratojugal, which were in the same modules, were later fused into the jugal bars in adult; the juvenile premaxilla, nasal, parasphenoid, pterygoid, vomer and maxilla, which were also in the same node-based modules, were later fused into the upper beak in adult stage.

For the 2-3 day old chick, the suture was not visible between the lateral ethmoid and the prefrontal and was later divided into separate bones in the same module when matured; the premature basicranium module (comprised of squamosal, frontal, parietal, postfrontal, exoccipital, orbitosphenoid, supraoccipital, basioccipital, and basisphenoid) closely resembled the braincase module in the adult (comprised of the braincase, postfrontal, parasphenoid, and prefrontal). The left jugal, quadrate, and quadratojugal were in the same three modules for both stages (Table S4).

For the medium ground finch, the palatine and vomer were in the same module in the juvenile and later fused in the adult; the juvenile maxilla module (comprised of premaxilla, nasal, maxilla, and quadratojugal) matched the adult beak and jugal bars; the adult braincase, quadrates and pterygoids were in the same module in juvenile: the module comprised of basioccipital, basisphenoid, vomer, supraoccipital, and left and right palatine, pterygoid, quadrate, quadratojugal, and exoccipital (Table S4).

**Supplementary Information 6: Supplementary Reference**

1. Benton, M. J. & Donoghue, P. C. J. Paleontological evidence to date the tree of life. *Mol. Biol. Evol.* **24**, 26–53 (2007).
2. Naish, D. & Barrett, P. M. *Dinosaurs: How They Lived and Evolved.* (The Natural History Museum, London., 2016).
3. Mayr, G. *Avian Evolution: The Fossil Record of Birds and its Paleobiological Significance (Topics in paleobiology series)*. (2017).
4. Elzanowski, A. A novel reconstruction of the skull of Archaeopteryx. *Netherlands J. Zool.* **51**, 207–215 (2001).
5. Rauhut, O. W. M. New observations on the skull of Archaeopteryx. *Palaontologische Zeitschrift* **88**, 211–221 (2014).
6. Smith-Paredes, D. *et al.* Dinosaur ossification centres in embryonic birds uncover developmental evolution of the skull. *Nat. Ecol. Evol.* **2**, 1966–1973 (2018).
7. Piras, P. *et al.* Morphological integration and functional modularity in the crocodilian skull. *Integr. Zool.* **9**, 498–516 (2014).
8. Felice, R. N. *et al.* Evolutionary integration and modularity in the archosaur cranium. *Integr. Comp. Biol.* (2019). doi:10.1093/icb/icz052
9. Sanger, T. J., Mahler, D. L., Abzhanov, A. & Losos, J. B. Roles for modularity and constraint in the evolution of cranial diversity among anolis lizards. *Evolution (N. Y).* **66**, 1525–1542 (2011).
10. Goswami, A. Morphological Integration in The Carnivoran Skull. *Evolution (N. Y).* **60**, 169–183 (2006).
11. Horton, J. An Anatomical Network Analysis of Crocodilian skull ecomorphology and modularity. (Imperial College London, 2018).
12. LJ, R. Phytools: An R package for phylogenetic comparative biology (and other things). *Methods Ecol. Evol.* **3**, 217–223 (2012).
13. Pennell, M. *et al.* geiger v2.0: an expanded suite of methods for fitting macroevolutionary models to phylogenetic trees. *Bioinformatics* **30**, 2216–2218 (2014).
14. Jarvis, E. D. *et al.* Whole-genome analyses resolve early branches in the tree of life of modern birds. *Science* **346**, 1320 LP – 1331 (2014).

29. Rinehart, L. F., Lucas, S. G., Heckert, A. B., Spielmann, J. A. & Celeskey, M. D. The Paleobiology of Coelophysis bauri (Cope) from the Upper Triassic

(Apachean) Whitaker quarry, New Mexico, with detailed analysis of a single quarry block. *New Mex. Museum Nat. Hist. Sci.* (2009).

30. Spielmann, J. A. *et al.* Oldest records of the late triasssic theropod dinosaur. *Nat. Hist.* 384–401 (2007).

31. Peyer, K. A reconsideration of Compsognathus from the Upper Tithonian of Canjuers, southeastern France. *J. Vertebr. Paleontol.* **26**, 879–896 (2006).

32. Gasparini, Z., Pol, D. & Spalletti, L. A. An unusual marine crocodyliform from the jurassic-cretaceous boundary of Patagonia. *Science* **311**, 70–73 (2006).

33. Pol, D. & Gasparini, Z. Skull anatomy of dakosaurus andiniensis (thalattosuchia: Crocodylomorpha) and the phylogenetic position of thalattosuchia. *J. Syst. Palaeontol.* **7**, 163–197 (2009).

34. Small, B. J. The Triassic Thecodontian Reptile Desmatosuchus: Osteology and Relationships. (Texas Tech University, 1985).

45. Field, D. J. *et al.* Complete Ichthyornis skull illuminates mosaic assembly of the

avian head. *Nature* **557**, 96–100 (2018).

53. Sereno, P. C. The ornithischian dinosaur Psittacosaurus from the Lower Cretaceous of Asia and the relationships of the Ceratopsia. (Columbia University, New York, 1987).

75. Heckert, A. B. & Lucas, S. G. Taxonomy, phylogeny, biostratigraphy,

biochronology,paleobiogeography,and evolution of the Late Triassic Aetosauria (Archosauria:Crurotarsi). *Zentralblatt für Geol. und Paläontologie Tl. I, H.* **Heft 11-12**, 1539–1587 (2000).

76. Small, B. J. Cranial anatomy of Desmatosuchus Haplocerus (Reptilia: Archosauria: Stagonolepididae). *Zool. J. Linn. Soc.* **136**, 97–111 (2002).

77. Csardi, G. & Nepusz, T. The igraph software package for complex network research, InterJournal, Complex Systems 1695. (2006).
